# Computational analysis of transcriptome signature repurposes low dose trifluoperazine for the treatment of fragile X syndrome in mouse model

**DOI:** 10.1101/683169

**Authors:** Qi Ding, Ferzin Sethna, Xue-Ting Wu, Zhuang Miao, Ping Chen, Yueqi Zhang, Hua Xiao, Wei Feng, Yue Feng, Xuan Li, Hongbing Wang

**Affiliations:** Department of Physiology, Michigan State University, East Lansing, MI 48824; Department of Genetics Program, Michigan State University, East Lansing, MI 48824; Key Laboratory of Synthetic Biology, Institute of Plant Physiology and Ecology, Shanghai Institutes for Biological Sciences, Chinese Academy of Sciences, Shanghai, 200032, China; Department of Pharmacology and Chemical Biology, Emory University School of Medicine, Atlanta, GA 30322

## Abstract

Fragile X syndrome (FXS), caused by mutations in fragile X mental retardation 1 gene (*FMR1*), is a prevailing genetic disorder of intellectual disability and autism. Currently, there is no efficacious medication for FXS. Here, we use transcriptome landscape as a holistic molecular phenotype/endpoint to identify potential therapeutic intervention. Throug*h in silico* screening with public gene signature database, computational analysis of transcriptome profile in *Fmr1* knockout (KO) neurons predicts therapeutic value of an FDA-approved drug trifluoperazine. Through experimental validation, we find that systemic administration of low dose trifluoperazine at 0.05 mg/kg attenuates multiple FXS- and autism-related behavioral symptoms. Moreover, computational analysis of transcriptome alteration caused by trifluoperazine suggests a new mechanism of action against PI3K (Phosphatidylinositol-4,5-bisphosphate 3-kinase) activity. Consistently, trifluoperazine suppresses PI3K activity and its down-stream targets Akt (protein kinase B) and S6K1 (S6 kinase 1) in neurons. Further, trifluoperazine normalizes the aberrantly elevated activity of Akt and S6K1 and enhanced protein synthesis in FXS mouse. In conclusion, our data demonstrate promising value of gene signature-based computation in identification of therapeutic strategy and repurposing drugs for neurological disorders, and suggest trifluoperazine as a potential practical treatment for FXS.

## INTRODUCTION

Fragile X syndrome (FXS), caused by mutations in the *FMR1* (fragile X mental retardation 1) gene, is the most common form of inherited intellectual disability and a leading cause of autism ^1^. As the most prevailing form of mutation, expansion of CGG repeats in the 5’ un-translated region of the *FMR1* gene inhibits its transcription and thereby preventing the expression of its gene product FMRP (fragile X mental retardation protein). FXS patients exhibit numerous neurological abnormalities including intellectual disability, social impairment, perseveration, and hyperactivity ^2^. Despite recent advances in the understanding of FMRP function, FXS pathophysiology, and identification of potential therapeutic targets ^3^, development of practical therapy has had limited success. As there is no efficacious medication to treat FXS, identification of practical therapeutic intervention is of urgent demand.

Recent studies with transcriptome landscapes in distinct neuronal populations suggest that gene signature may represent a holistic molecular outcome and an indicator for cell type classification and function ^4^. As a molecular phenotype and pathological endpoint, transcriptome alteration has been observed in patient samples from major psychiatric disorders ^5, 6^. Intriguingly, using transcriptome landscape as a holistic outcome may offer an unbiased approach to reveal disease mechanism as well as neuropathology and etiology that are unique or overlapping among psychiatric disorders ^7^. Recent studies also suggest that analysis of disease-associated transcriptome profile may also predict potential therapy or aid treatment development. For example, antipsychotics-induced transcriptome changes are negatively correlated with those identified in schizophrenia samples. In contrast, psychotomimetic phencyclidine triggers transcriptome changes that overlap with the disease profile ^7^. Regarding its potential application in drug discovery, comparison of psychiatric disease-associated transcriptomes with drug-induced transcript alterations in public database can computationally reveal the previously validated drug-disease pairs ^8^. However, whether the transcriptome-based computational approach can help identify new therapeutic or repurposing drugs for new application is of great interest but remains to be examined ^8^.

In this study, we identified significant transcriptome changes in *Fmr1* deficient neurons. We compared the transcriptome signature in *Fmr1* knockout (KO) neuron with those in the connectivity Map (CMAP) database ^9^, which contains over 7000 reference gene expression profiles representing transcriptome changes affected by 1309 compounds/drugs. Computational analysis outcome predicts therapeutic value of an FDA-approved drug trifluoperazine. Using the FXS mouse model, we found significant effects of trifluoperazine on correcting FXS- and autism-associated symptoms. Further analysis with drug-induced transcriptome changes revealed a new pharmacologically inhibitory activity of trifluoperazine against PI3K (Phosphatidylinositol-4,5-bisphosphate 3-kinase). Administration of trifluoperazine normalizes the aberrantly elevated PI3K/Akt (protein kinase B)-S6K1 (S6 kinase 1) signaling in FXS. Our data support the value of computational transcriptome analysis in therapeutic development, and suggest trifluoperazine as a potential practical medication for FXS treatment.

## MATERIALS AND METHODS

### Animals

Male *Fmr1* knockout (KO) mice and their wild type (WT) littermates on C57BL/6 background were obtained from breeding pairs consisting of heterozygous female and WT male mouse. Animals were housed in the Campus Animal Research facility. The Institutional Animal Care and Use Committee approved all procedures. The mice had *ad libitum* access to water and food and were housed under 12 h dark/light cycle.

### Transcriptome analysis of gene expression in *Fmr1* KO neurons

Whole genome transcript changes were determined by RNA-seq with total RNA extracted from primary hippocampal cultures at DIV (days in vitro) 14. Triplicate samples/repeats were analyzed and compared between wild type (WT) and *Fmr1* KO genotype. Total RNA was extracted by the TRIzol method (Invitrogen) followed by purification with RNAeasy kit from Qiagen. Purified total RNA with RIN>8 was subjected to RNA-seq. Raw data were processed through adaptor cleaning (with Fastx Clipper), trimming (with sickle, https://github.com/najoshi/sickle), junk sequence removal, mapping clean reads to the mouse reference genome (version: GRCm38) using TopHat2, and gene construction with Cufflinks ^10^. Differentially expressed genes (DEGs) between WT and *Fmr1* KO samples were assessed with Cuffdiff. The resulting DEGs were ranked based on expression differences. GO (gene ontology) enrichment and KEGG (Kyoto Encyclopedia of Genes and Genomes) pathways were identified with DAVID (Database for Annotation, Visualization and Integrated Discovery; http://david.abcc.ncifcrf.gov/home.jsp) ^11^.

### Computational prediction on drug connectivity by using CMAP database

Transcriptome alterations in *Fmr1* KO neurons, including up- and down-regulated DEGs (differentially expressed genes) were compared with gene signatures in CMAP database (http://www.broadintitute.org/cmap/) ^9^. DEGs were first converted to corresponding human gene probes using Batch Query at NetAffx (http://www.affymetrix.com/analysis/netaffx/index.affx). The detailed list of signature probes for DEGs in *Fmr1* KO neurons is shown in Supplementary Data 1.

To obtain gene expression signature for trifluoperazine, raw microarray data sets in PC3 cells treated with trifluoperazine and corresponding vehicle control were downloaded from CMAP database. The raw data were normalized using robust multi-array average algorithm (RMA) ^12^ from the R package “affy” (v1.44.0) ^13^ (https://www.bioconductor.org/about/). Genes were then ranked based on the expression differences between the drug-treated and vehicle controls. The signature genes were selected using ‘up threshold’ and ‘down threshold’ as defined by CMAP. The detailed list of signature probes for trifluoperazine is shown in Supplementary Data 2.

Gene signature of *Fmr1* KO neuron or trifluoperazine was uploaded to CMAP quick query page to search for compounds that induce similar and oppositional transcriptome changes. CMAP has devised a tool to compute enrichment of up- and down-regulated genes from transcriptome changes in compound/drug-treated cells, and the results for all instances related to a specific compound are combined to form ‘connectivity score’ (i.e. similarity mean) for each compound ^9^. After executing a query, the top hit chemical list was returned, displaying each compound’s associated similarity mean, number of arrays, enrichment, and p value.

### *In vivo* drug administration

Trifluoperazine dihydrochloride (Sigma-Aldrich), SCH23390 (Sigma-Aldrich), and raclopride (Sigma-Aldrich) solution in water was i.p. injected into mice at 0.05 mg/kg, 0.05 mg/kg, and 0.3 mg/kg, respectively. For passive avoidance, trifluoperazine was administered 1 hour before or after training. For all other behavioral paradigms, drug was administered 1 hour before examination. Control mice were treated similarly but injected with vehicle.

To test the effect of repeated drug administration, trifluoperazine at 0.05 mg/kg was injected once per day for 8 days. One hour after the last injection, mice were examined by behavior tests.

### Behavioral tests

To examine sociability, 2- to 3-month old mouse was placed in the center chamber of a 3-chamber social interaction box. Mouse was allowed to freely explore all 3 chambers for 5 min. Mouse showing significant preference for a particular chamber was not used for further testing. Following the exploration, mouse was momentarily restricted (for less than 10 sec) in the center chamber with the entry doors to the side chamber closed. The 10-min sociability test started with placement of a novel stimulus mouse in a wire enclosure and an empty wire enclosure, respectively, in one of the side chambers, followed by opening the chamber-connecting doors. As a social interaction index, total time spent in sniffing the novel stimulus mouse enclosure was recorded.

To perform the marble-burying test, 2- to 3-month old mice were placed in a box (27 cm by 15 cm box with 12-cm high walls) with 7.5 cm depth of bedding for 1 hour prior to the test. The mouse was then briefly removed from the testing box and 15 marbles were evenly arranged in a 5 by 3 pattern on the surface of the bedding. The mouse was reintroduced into the testing box and was allowed to bury marbles for 10 min. At the end of the testing period, the mouse was removed from the box and the number of marbles that were fully buried, partially buried, and left on the surface was counted.

To determine stereotypic digging, 2.5-to 3.5-month old mice were introduced to fresh mouse cage with 1-inch bedding material. During the 5-min examination, latency to the first digging episode, total number of digging episodes, and accumulative time spent in digging were recorded.

To determine stereotypic self-grooming, 2.5-to 3.5-month old mice were placed in a plexiglas cage with 1 cm fresh bedding without nesting material. After a 10-min habituation period, the number of self-grooming episode and duration were scored for 10 min ^14^.

To examine animal activity in light-dark test, 2.5-to 3.5-month old mice were placed in the dark half of the light-dark chamber and the trap door was opened 1 min later. The mice were allowed to move freely between the dark and the lit chambers for 5 min. Time spent in the lit chamber and number of crossing into the lit side were recorded.

To determine animal activity in an open field arena, 2- to 3-month old mice were placed in the center of an open field chamber, and were allowed to move freely for 1 hour. Ambulatory movement distance in the whole arena and in the center area of the open field were determined every 10-min during the 1-hour testing period by the TruScan Photo Beam Activity System (Coulbourn Instruments, Whitehall, PA).

To examine passive avoidance learning, 2.5- to 3.5-month old mice were introduced into the lit half of the passive avoidance chamber (Coulbourn Instruments, Whitehall, PA) and allowed to explore for 1 min before the trap door was opened. The trap door was closed as soon as the mouse entered the dark chamber. A mild foot shock (0.7 mA for 2 sec) was immediately delivered. The mouse was removed from the dark chamber and returned to its home cage 30 sec after the delivery of foot shock. The mice were tested 24 hours after training. During testing, the mouse was put in the lit chamber and crossover latency to the dark chamber was recorded. If mice stayed in the lit chamber for more than 600 secs, they were manually removed from the chamber, and 600 secs was used as their crossover latency. In a stronger training paradigm, 3 electric foot shocks (0.7 mA for 2 secs per shock with 10 sec interval) were delivered during training when animal entered the dark chamber.

To measure audiogenic seizures (AGS), twenty-one- to twenty-four-day old mice were placed in a box (30 cm L by 17 cm W by 12 cm H) with a flat plastic lid. A personal alarm (from Streetwise, item # SWPDAL) was taped to the lid of the box and wired to a DC power supply to keep the sound amplitude constant. The mouse was allowed to acclimatize to the box for 5 min, following which a 120-dB sound was emitted from the alarm for 2 min. The number of mice undergoing seizure within the 2-min period was counted. Audiogenic seizures were considered when wild running and/or clonic/tonic seizures occur.

Some of the animals were re-used in different behavioral tests (see Supplementary Fig. 1). Mice used for open field test were later used for social interaction. Some mice used for marble-burying test were later used for light-dark test. Mice used for stereotypic digging were used later for passive avoidance. The interval between the two behavioral tests was at least 10 days.

### Neuronal cell culture and measurements of protein synthesis

Primary hippocampal neurons were cultured with samples obtained from postnatal day 0 WT and *Fmr1* KO mice. Protein synthesis was determined by the SUnSET method ^15, 16^. DIV 14 hippocampal neurons were pre-treated with trifluoperazine at various concentrations (0.5, 1, 2, 10, and 20 μM as indicated) for 30 min followed by 5 μg/ml puromycin (Sigma, Cat #P8833) treatment for 30 min. Cells were lysed in Buffer H (50 mM β-glycerophosphate, 1.5 mM EGTA, 0.1 mM Na_3_VO_4_, 1 mM DTT). The samples were sonicated and centrifuged. An aliquot of the supernatant was used to determine protein concentration, and the rest was denatured in Laemmli buffer. 20 μg protein was separated by 4-20% SDS-PAGE (Invitrogen) and transferred onto nitrocellulose membranes. The membranes were probed with anti-puromycin antibody (KeraFAST, Cat # EQ0001, 1:1000). The relative amount of loading was determined by β-actin. ImageJ was used to measure the combined signal intensity of proteins with molecular weights ranging from 15 to 250 kDa.

### Examination of drug effects by Western blot

To examine the *in vitro* effects, DIV 14 hippocampal neurons (WT and *Fmr1* KO as indicated) were treated with vehicle, vorinostat (20 μM), trichostatin A (20 μM), LY-294002 (20 *μ*M), and trifluoperazine (at 0.25, 0.5, 1, 2, 10, and 20 μM as indicated) for 1 hour. Immediately after treatment, neurons were lysed in Laemmli buffer (62.5 mM Tris-HCL, pH 6.8, 10% glycerol, 2% SDS, 5% 2-mercaptoethanol, 0.005% bromophenol blue). To examine the *in vivo* effect, 2- to 3-month old wild type (WT) and *Fmr1* KO mice were injected with 0.05 mg/kg trifluoperazine or vehicle. Hippocampus was rapidly dissected 1 hour after injection and homogenized in RIPA buffer (50 mM Tris-HCL, pH 7.6, 150 mM NaCl, 1 mM EDTA, 0.25% sodium deoxycholate, 0.5% NP-40). Protein content in hippocampal homogenates was determined using Bradford’s assay. Same amount of cell extract/lysate was separated by SDS-PAGE followed by transferring to nitrocellulose membranes. The following antibodies were used to detect the corresponding targets. Anti-S6K1 (Cat #2708, 1:1000 dilution), anti-phospho-S6K1 (at Thr389) (Cat # 9234, 1:1000 dilution), anti-phospho-Akt (at Ser473) (Cat # 4060, 1:1000 dilution), anti-Akt (Cat # 9272, 1:1000 dilution), anti-acetylated histone H3 at K9 (Cat #9649P, 1:1000 dilution), anti-acetylated histone H4 at K8 (Cat #2594P, 1:1000 dilution), anti-histone H3 (Cat #4499P, 1:1000 dilution), and anti-histone H4 (Cat #13919P, 1:1000 dilution) were from Cell Signaling. β-actin antibody was from Sigma (Cat #A5441, 1:10,000 dilution). Anti-FMRP (mouse monoclonal antibody 2F5, 1:1000) was a generous gift from Dr. Jennifer Darnell at the Rockefeller University. For quantification purpose, level of phospho-S6K1, phospho-Akt, acetylated-H3, and acetylated-H4 was normalized to the level of total S6K1, Akt, H3, and H4, respectively. The membrane was normally first blotted for the phosphorylated and acetylated proteins. Then, the same membrane was stripped; the effectiveness of stripping/removing the phospho- or acetylation-specific antibodies was checked by blotting with secondary antibody alone. The successfully stripped membrane was then washed and re-probed with antibodies against the corresponding total protein. The Western blot signal was detected by the Odyssey digital imaging system. Relative intensity of the Western blot signal in the no treatment control group, which was quantified using ImageJ (NIH, MD, USA), was defined as 1. Signal in the treatment samples was compared to the control group.

### PI3 kinase assay

Hippocampal neurons (cultured from WT mice) on DIV 14 were treated with trifluoperazine (20 μM) or vehicle for 30 min. Neurons were then washed once with PBS and lysed in 100 μl lysis buffer (50 mM Tris pH 7.4, 40 mM NaCl, 1 mM EDTA, 0.5% Triton, 1.5 mM Na_3_VO_4_, 50 mM NaF, 10 mM sodium pyrophosphate) with proteinase inhibitor (Roche) on ice for 10 min. Lysates were cleared by centrifugation and protein concentration was determined by Bradford method. PI3 kinase activity was measured using PI3 kinase activity ELISA kit from Echelon Biosciences according to the manufacture’s protocol. Briefly, 30 μl of lysate was incubated with 30 μl of 10 μM PIP2 and 20 μM trifluoperazine or vehicle at 37° C for 3 hours. The reaction buffer contained 2 mM DTT and 100 μM ATP. The amount of PIP3 produced by PI3 kinase was determined by ELISA. The relative amounts of PIP3 in the samples were used to determine the activity of PI3 kinase.

### Examination of neuron viability

DIV 14 hippocampal neurons were treated with vehicle or 10 μM or 20 μM trifluoperazine for 1 hour. As a positive control, NMDA treatment (50 μM for 30 min) was used to trigger neuronal cell death. Immediately after treatment, drugs were removed by replacement with conditioned medium. 20 hours after treatment, neurons were fixed with 4% formaldehyde and stained with DAPI. As described in our previous study ^17^, neuron survival (i.e. viability) was quantified by the ratio of neurons showing diffused nuclear DAPI staining to neurons showing condensed nuclear DAPI staining.

### Data collection and statistics analysis

Animals subjected to different treatment were randomly assigned. Sample size was estimated by power analysis (80% power and probability level of 0.05 along with the anticipated effect size) or based on previous reports. Data were collected from two (for biochemical analyses with primary neurons) or more than two litters (for behavioral analyses and *in vivo* biochemical analysis) and combined. Experimenters were blind to the treatments and genotypes, which were decoded before data analyses. Student’s t-test (two-sided) was used to compare data collected from two groups. One-way ANOVA and Tukey’s *post hoc* tests were used to analyze data involving different treatments. Two-way ANOVA and Holm-Sidak *post hoc* tests were to analyze data involving genotype effects and treatment effects. Three-way repeated measures ANOVA was used to analyze the open field data at different time points during the 60-min test. When normal distribution and equal variance were not detected (i.e. crossover latency during testing for passive avoidance), the numerical values were first converted to categorical data (latency of 600 sec or less than 600 secs was assigned to different category) and then analyzed by Fisher’s exact test. Data reporting AGS were analyzed by Chi-square and/or Fisher’s exact test. Data are expressed as mean ± SEM. Differences with *p* value less than 0.05 were considered significant. SPSS 11.5 for Windows (IBM) was used for all data analysis.

The statistical method for CMAP analysis was described previously ^9^. In brief, the *p* value was computed using Kolmogorov-Smirnov statistics. It is estimated empirically by computing the enrichment of 100,000 sets of instances randomly selected from all instances in the result.

### Data Availability

RNA-seq data have been deposited and are available in Gene Expression Omnibus under accession number GSE114015. All individual data point is presented in figures, and disclosed in the source data files. The full gel Western blot images are shown in the source data files. All data associated with this study are available from the corresponding author upon reasonable request.

## RESULTS

### Screening of CMAP with transcriptome signature of *Fmr1* KO neurons identifies trifluoperazine as a potential therapeutic

Transcriptome alteration represents an emerging molecular phenotype and pathological signature of chronic diseases including psychiatric disorders ^5–8^. Here, we used next generation sequencing (NGS) to determine genome-wide changes of gene transcript in *Fmr1* KO hippocampal neurons. Compared to wild type (WT) neurons, 587 gene transcripts were up-regulated and 724 were down-regulated in *Fmr1* KO samples (Fig. 1a; Supplementary Data 3). Enrichment analysis identified significant changes in 138 GO processes (complete list in Supplementary Table 1; top 15 pathways in Fig. 1b and Supplementary Table 2) and 11 KEGG pathways (Fig. 1c; Supplementary Table 3). Some enrichment groups are related to neural development, calcium homeostasis and signaling, p53 signaling, Rap1 signaling, and MAPK signaling (Fig. 1b and 1c), the alterations of which are implicated in FXS ^18–21^. As alterations of numerous transcripts occur in multiple pathways (Supplementary Table 2 and 3), it is challenging to pinpoint which specific DEG (differentially expressed gene) plays a causal role in FXS pathology.

**Figure 1.**
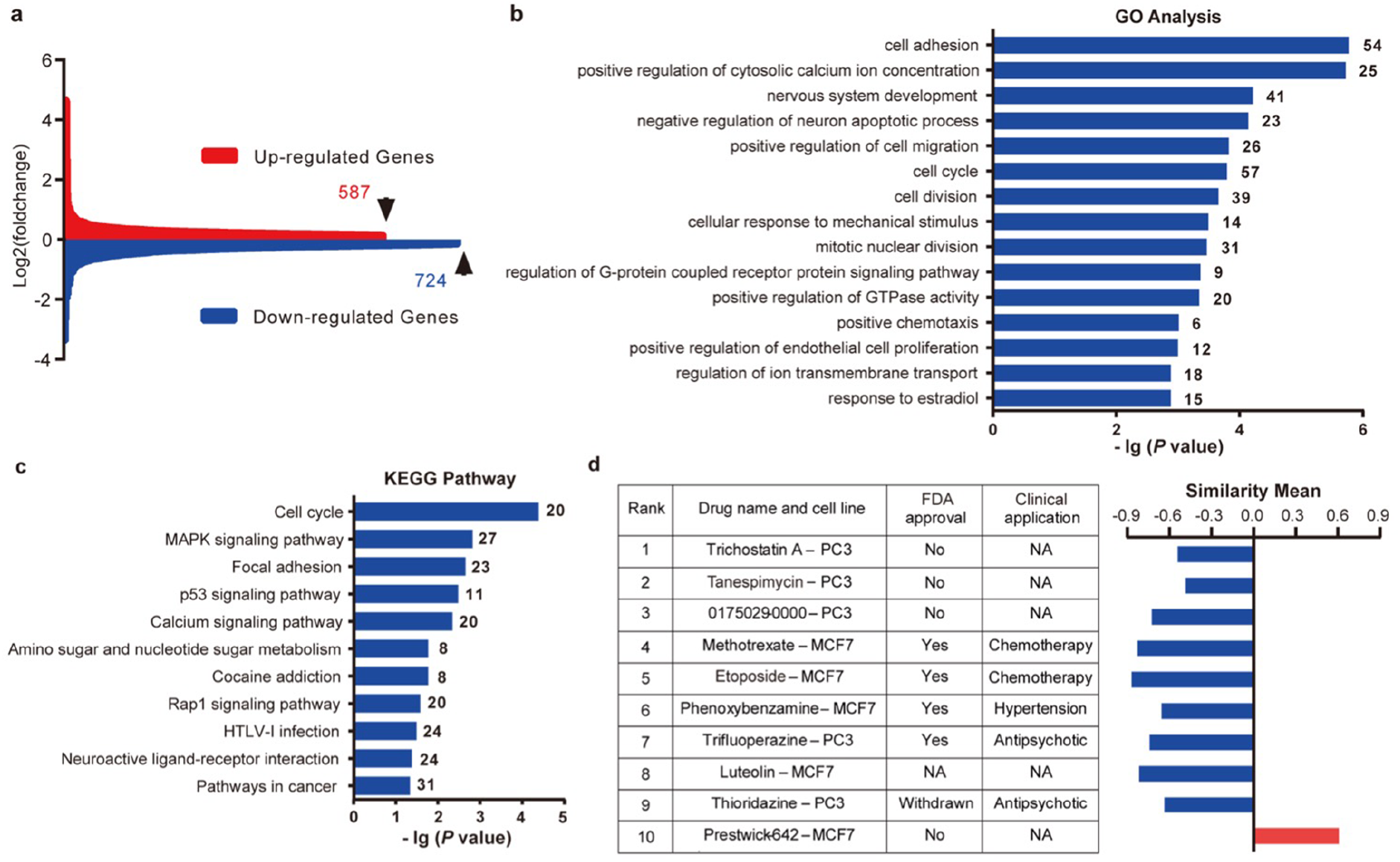
RNA-seq and connectivity Map (CMAP) analysis of differentially expressed genes (DEG) in *Fmr1* knockout (KO) hippocampal neuron. (**a**) Gene expression differences between wild type (WT) and *Fmr1* KO samples. There are 587 up-regulated and 724 down-regulated genes in *Fmr1* KO hippocampal neuron. (**b**) Top 15 GO biology processes that are associated with DEG between WT and *Fmr1*KO samples. (**c**) KEGG pathways that are associated with DEG between WT and *Fmr1* KO samples. (**d**) Top-ranked 10 CMAP compounds/drugs that induce transcriptome alterations oppositional to (indicated by negative similarity mean) or overlapping with (indicated by positive similarity mean) that caused by *Fmr1* deficiency. Rank is determined by *P* value and enrichment score. Drug name and cell line indicate the name of compound used for treatment with specific cell lines in CMAP database. Full information of *P* value, enrichment score, and similarity mean is shown in Supplementary Table 4.

Recent computational analysis with transcriptomes imputed from GWAS data and CMAP drug-induced gene signature suggests an intriguing strategy to identify potential treatment for psychiatric disorders ^8^. The strategy, which identifies therapeutic compounds inducing gene signature oppositional to disease signature (indicated by negative similarity mean), was supported by known medications used for psychiatric conditions. For example, predicted therapeutic compounds for schizophrenia are enriched for antipsychotics ^8^. To examine the value of this unbiased approach, we compared transcriptome signature of *Fmr1* KO neurons with drug-induced signatures in CMAP ^9^. We identified negative and positive correlations between the FXS-drug pair. Among the top 10 hits ranked by p value and enrichment, 9 compounds/drugs induce oppositional transcriptome changes (as indicated by negative similarity mean) (Fig. 1d, Supplementary Table 4), predicating potential therapeutic value. From the 4 FDA-approved drugs, we chose trifluoperazine, which can effectively cross the blood brain barrier (BBB) ^22, 23^ and has been used to treat central nervous system disorders, for experimental validation of therapeutic efficacy. The other 3 FDA-approved compounds have been used for cancer chemotherapy and hypertension treatment, respectively, and were not selected for the current study.

### Trifluoperazine corrects the FXS- and autism-related behavioral symptoms in *Fmr1* KO mice

Previous studies demonstrate that efficacy doses of trifluoperazine used for schizophrenia and anxiety show extrapyramidal side effects in patients ^24^, but low dose trifluoperazine is well tolerated in both human and animal models ^25^. We examined the key FXS-associated behavioral symptoms following low dose i.p. injection (at 0.05 mg/kg).

We first examined the effects of trifluoperazine on sociability, reduction of which prevalently occurs in FXS and autism. In the 3-chamber social interaction test (genotype effect: *F*_1, 35_ = 14.851, p < 0.001; treatment effect: *F*_1, 35_ = 3.475, p = 0.071; genotype X treatment interaction: *F*_1, 35_ = 14.704, p < 0.005), vehicle-injected *Fmr1* KO mice showed less interaction with the social subject (i.e. a novel stimulus mouse) than WT mice (Fig. 2a; *p* < 0.001). Trifluoperazine did not affect WT mice (*p* = 0.189) but improved sociability in *Fmr1* KO mice to the WT level (Fig. 2a; *p* < 0.001).

**Figure 2.**
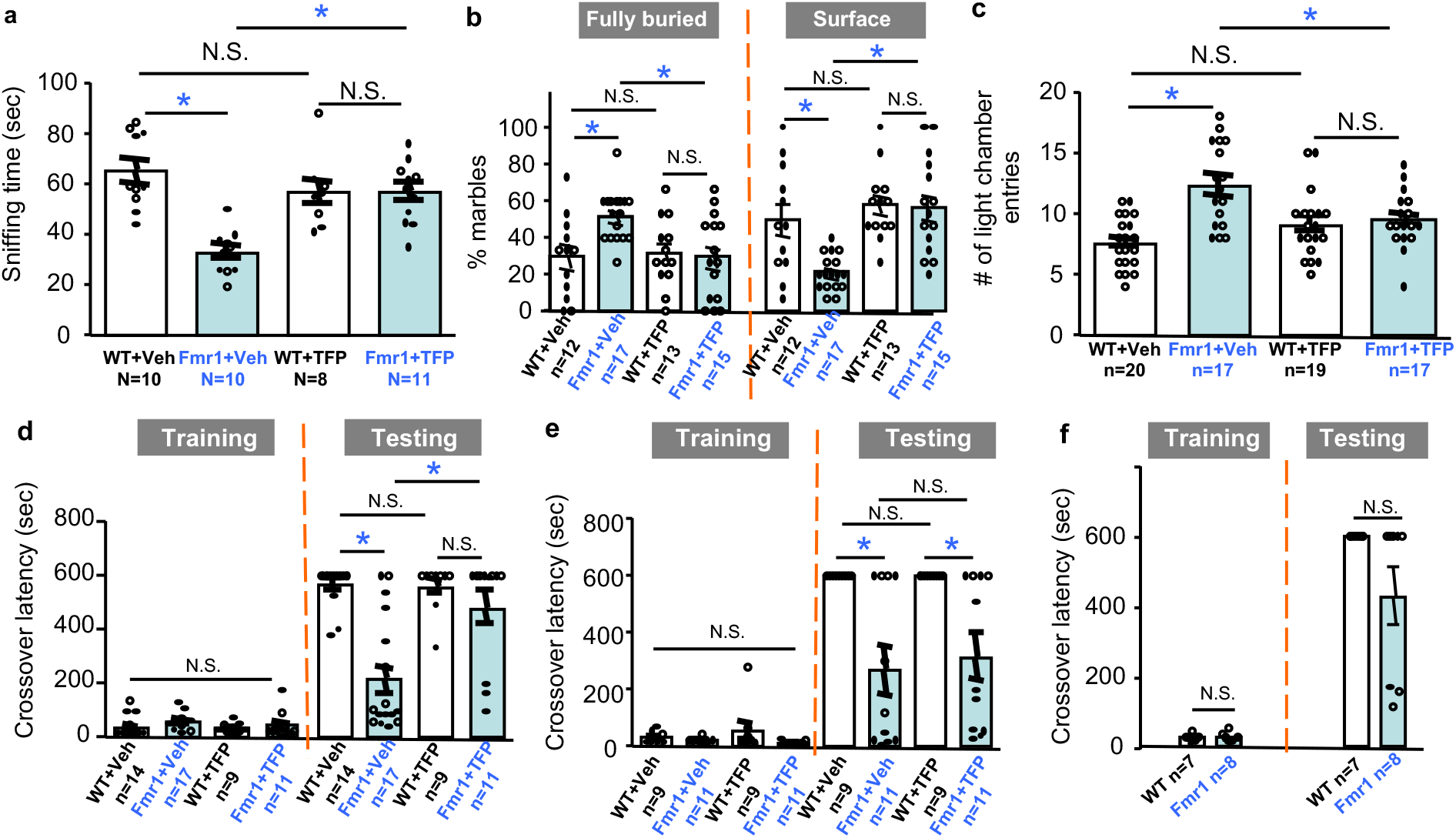
Trifluoperazine corrects FXS- and autism-associated behavioral symptoms in *Fmr1* KO mice. Trifluoperazine (TFP, at 0.05 mg/kg) or vehicle (Veh) was administered into wild type (WT) and *Fmr1* KO mice (*Fmr1*) 1 hour before testing (**a** to **c**). (**a**) TFP improves sociability in *Fmr1* KO and does not affect WT mice. (**b**) The percentage of fully buried marbles and marbles on the surface (as indicated) was determined for WT and *Fmr1* KO mice following TFP or vehicle administration. (**c**) TFP corrects the increased locomotive transition between the lit and dark chamber in *Fmr1* KO mice. The total number of entries to the lit chamber was scored. (**d**) Mice were injected with vehicle or trifluoperazine 60 min before passive avoidance training. Mice were tested 24 hours after training, and crossover latency was scored. (**e**) Mice were injected with vehicle or trifluoperazine 60 min after passive avoidance training. Mice were tested 24 hours after training, and crossover latency was scored. There is no difference in crossover latency among the 4 groups during training (genotype effect: *F*_1, 36_ = 3.218, *p* = 0.081; drug effect: *F*_1, 36_ = 0.309, *p* = 0.581; genotype X drug interaction: *F*_1, 36_ = 1.256, *p* = 0.270). *Fmr1* KO mice showed impaired memory and not corrected by trifluoperazine. (**f**) Mice received 3 mild electric foot shocks during passive avoidance training. Mice were tested 24 hours after training, and crossover latency was scored. For **a**, **b**, and **c**, two-way ANOVA followed by Holm-Sidak test was used to analyze data (*: *p* < 0.05; N.S.: not significant). For **d** and **e**, two-way ANOVA and Fisher’s exact test was used to analyze training and testing data, respectively (*: *p* < 0.05; N.S.: not significant). For **f**, Student’s t-test and Fisher’s exact test was used to analyze training and testing data, respectively (N.S.: not significant).

*Fmr1* KO mice show significant repetitive/stereotypic behavior, recapitulating perseverative symptoms in FXS and autism patients. We examined animal behavior in marble-burying test, which has been considered to measure repetitive behavior and barely involves novelty-induced anxiety ^26^. We found that trifluoperazine had no effect on WT but normalized the excessive marble burying behavior in *Fmr1* KO mice (for fully buried marbles in Fig. 2b; genotype effect: *F*_1, 53_ = 3.726, *p* = 0.059; treatment effect: *F*_1, 53_ = 3.808, *p* = 0.056; genotype X treatment interaction: *F*_1,53_ =5.758, *p* < 0.05; *post-hoc* Holm-Sidak test detected significant difference between vehicle-treated WT and *Fmr1* KO mice) (for marbles on the surface in Fig. 2b; genotype effect: *F*_1, 53_ = 6.255, *p* < 0.02; treatment effect: *F*_1, 53_ = 14.217, *p* < 0.001; genotype X treatment interaction: *F*_1,53_ = 5.383, *p* < 0.03; *post-hoc* Holm-Sidak test detected significant difference between vehicle-treated WT and *Fmr1* KO mice). The number of marbles partially buried by WT and *Fmr1* KO mice was similar (Supplementary Fig. 2; genotype effect: *F*_1, 53_ = 2.01, *p* = 0.162). Due to overall increase in combined number of fully buried and surface marbles, the number of partially buried marbles was decreased in both WT and *Fmr1* KO mice injected with trifluoperazine (Supplementary Fig. 2; drug effect: *F*_1, 53_ = 11.393, *p* < 0.005; genotype-drug interaction: *F*_1, 53_ = 0.096, *p* = 0.758). We further examined *Fmr1* KO mice with light-dark test. The *Fmr1* KO mice showed more transition between the light and dark chambers than WT animals, indicating both hyperactivity and repetitive behavior (Fig. 2c; genotype effect: *F*_1,69_ = 16.69, *p* < 0.001). *Fmr1* KO and WT mice spent comparable time in the light and dark chamber (genotype effect: *F*_1,69_ = 0.428, *p* = 0.515; treatment effect: *F*_1,69_ = 0.046, *p* = 0.830; genotype X treatment interaction: genotype effect: *F*_1,69_ = 0.820, *p* = 0.368) (Supplementary Fig. 3). Trifluoperazine corrected the excessive transition in *Fmr1* KO mice but did not affect WT mice (Fig. 2c; treatment effect: *F*_1, 69_ = 1.276, *p* = 0.263; genotype-treatment interaction: *F*_1, 69_ = 11.73, *p* < 0.005). In other stereotypic behavior tests, *Fmr1* KO mice displayed normal self-grooming (Supplementary Fig. 4) but excessive digging behavior (Supplementary Fig. 5). Trifluoperazine did not show significant effect on correcting excessive digging (Supplementary Fig. 5).

Hyperactivity is observed in both human FXS patients and *Fmr1* KO mice ^27, 28^. In a novel open field arena, we confirmed that *Fmr1* KO mice have enhanced locomotor activity in the whole arena (Supplementary Fig. 6a1 and 6a2; genotype effect: *F*_1, 28_ = 24.226, *p* < 0.0001) as well as in the center area (Supplementary Fig. 6b1 and 6b2; genotype effect: *F*_1, 28_ = 17.106, *p* < 0.0001). We did not detect significant effect of trifluoperazine on locomotion in the whole arena (drug effect: *F*_1, 28_ = 0.762, *p* = 0.39; genotype X drug interaction: *F*_1, 28_ = 1.592, *p* = 0.217) or the center area (drug effect: *F*_1, 28_ = 1.183, *p* = 0.286; genotype X drug interaction: *F*_1, 28_ = 0.376, *p* = 0.545).

FXS is the leading cause of inherited intellectual disability. To determine certain aspect of cognitive deficits, we examined passive avoidance memory. Trifluoperazine or vehicle was administered before training, during which animals received an electric foot shock once entering the dark chamber. 24 hours after training, animals were tested for memory formation, which is implicated by increased latency to cross over and enter the dark chamber. All groups of animals showed similar crossover latency during training (Fig. 2d, genotype effect: *F*_1, 47_ = 2.069, *p* = 0.157; drug effect: *F*_1, 47_ = 2.424, *p* = 0.126; genotype-drug interaction: *F*_1, 47_ = 0.248, *p* = 0.569). During testing, vehicle-treated WT mice showed significantly longer crossover latency than vehicle-treated *Fmr1* KO mice (Fig. 2d, *p* < 0.05, Fisher’s exact test). This confirms the impaired passive avoidance memory in the FXS mice ^27, 28^. Although trifluoperazine did not have an effect on WT animals, crossover latency in trifluoperazine-treated *Fmr1* KO mice significantly increased to the WT level (Fig. 2d, *p* < 0.05, Fisher’s exact test). We expect that the improved memory in *Fmr1* KO mice is not due to trifluoperazine effect on locomotion. This is because the memory test was performed 24 hours after trifluoperazine administration. When tested in a similar context (i.e. movement in the light-dark box test), the effect of trifluoperazine on locomotion is evident 1 hour (Fig. 2c) but not 24 hours after drug administration (Supplementary Fig. 7; genotype effect: *F*_1, 22_ = 18.631, *p* < 0.001; drug effect: *F*_1, 22_ = 0.325, *p* = 0.574; genotype X drug interaction: *F*_1, 22_ = 0.166, *p* = 0.688 for the number of transition between the lit and dark chambers). Interestingly, when trifluoperazine was administered after training, it had no significant effect on correcting the defective passive avoidance memory in *Fmr1* KO mice (Fig. 2e). This suggests an intriguing possibility that lack of FMRP causes impaired learning but not memory consolidation. Supportively, when trained by a stronger learning paradigm, during which 3 consecutive electric foot shocks were delivered, *Fmr1* KO mice showed normal passive avoidance memory (Fig. 2f).

Hyper sensitivity and susceptibility to seizures are common in FXS patients. Here, we confirmed that WT mice do not show audiogenic seizure (AGS), which is a major behavioral phenotype in FXS mouse model ^29^. While *Fmr1* KO mice receiving trifluoperazine displayed reduction in AGS, the drug effect is marginal but not significant (Supplementary Fig. 8; *p* = 0.113 for wild running and *p* = 0.273 for seizure).

### Effects of repeated trifluoperazine administration on behavioral symptoms

One potential problem related to treatment is drug desensitization following repeated administration. Trifluoperazine or vehicle was administered once per day for 8 days. We examined light-dark test and social interaction 1 hour after the last drug administration. While the vehicle-treated *Fmr1* KO mice showed more transitions between the two chambers, the trifluoperazine-treated group showed fewer transitions, which were comparable to the WT level (Fig. 3a1; genotype effect: *F*_1, 25_ = 27.303, *p* < 0.001; drug effect: *F*_1, 25_ = 27.303, *p* < 0.001; genotype X drug interaction: *F*_1, 25_ = 11.613, *p* < 0.005). Repeated trifluoperazine had no effect on the time spent in each chamber (Fig. 3a2; genotype effect: *F*_1, 25_ = 0.287, *p* = 0.597; drug effect: *F*_1, 25_ = 0.019, *p* = 0.892; genotype X drug interaction: *F*_1, 25_ = 0.079, *p* = 0.781).

**Figure 3.**
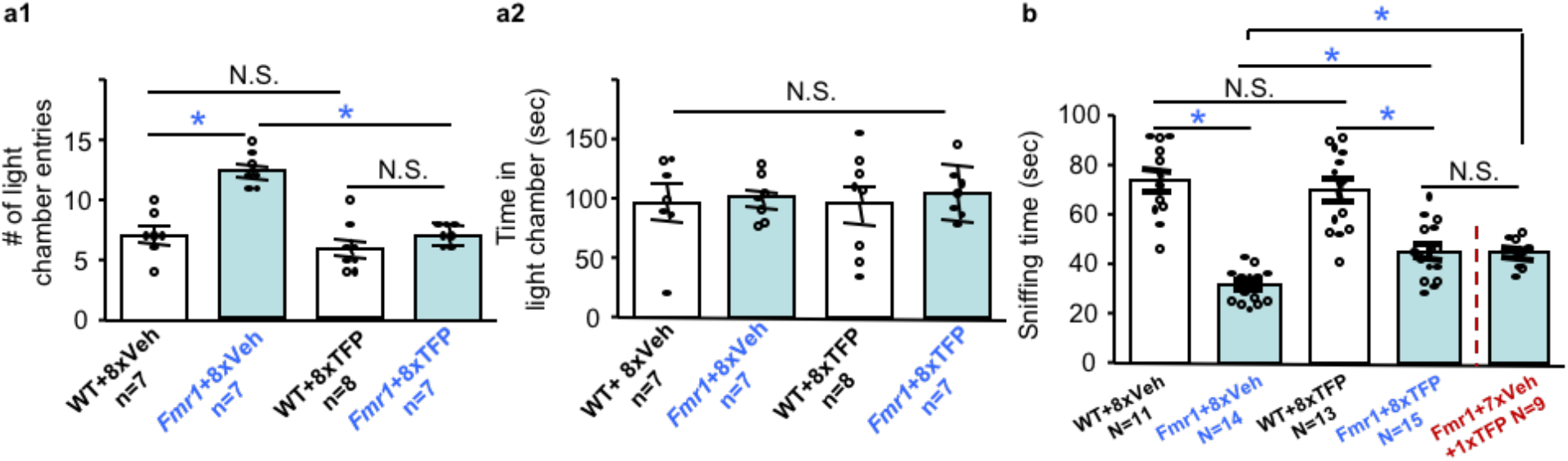
Repeated administration of trifluoperazine (TFP) does not cause desensitization to the drug. Wild type (WT) and *Fmr1* KO mice (Fmr1) were injected with TFP (0.05 mg/kg) or vehicle daily for 8 days. Mice were subjected to light-dark test (**a**) or social interaction test (**b**) 1 hour after the last injection. In a separate *Fmr1* KO cohort, mice were treated with 7 daily injection of vehicle and 1 TFP injection on day 8. Then, social interaction was examined 1 hour after TFP injection (**b**). For light-dark test, number of locomotive transition between the light and dark chamber (**a1**) and time spent in the light chamber (**a2**) are presented. For social interaction, total time spent in sniffing the novel stimulus mouse enclosure is presented (**b**). Two-way ANOVA followed by Holm-Sidak test was used to analyze the data. Student’s t-test was used to compare *Fmr1* KO mice receiving 7 vehicle and 1 TFP injection with other groups in **b**. *: *p* < 0.05; N.S.: not significant.

In another independent behavioral paradigm, repeated trifluoperazine improved social interaction in *Fmr1* KO mice (Fig. 3b). However, the treated-*Fmr1* KO mice still showed less social interaction than WT mice (Fig. 3b; genotype effect: *F*_1, 49_ = 83.685, *p* < 0.001; drug effect: *F*_1, 49_ = 1.896, *p* = 0.175; genotype X drug interaction: *F*_1, 49_ = 5.985, *p* < 0.05). This is likely due to that sociability in *Fmr1* KO mice is more sensitive to the repeated injection procedure (Supplementary Fig. 9). With another cohort of *Fmr1* KO mice, we found that 7 repeated vehicle injections followed by 1 trifluoperazine injection had similar effect to that of 8 repeated trifluoperazine injections (Fig. 3b). These data indicate that *Fmr1* KO mice are not desensitized to trifluoperazine following repeated exposures to the drug.

### Trifluoperazine normalizes the elevated protein synthesis in *Fmr1* KO neurons

Enhanced basal protein synthesis is thought to be the core cellular abnormality associated with FXS ^1, 3, 30^. Here, we labeled newly synthesized proteins in neurons with puromycin using the SUnSET method ^15, 16^. We observed enhanced protein synthesis in *Fmr1* KO neurons compared to WT neurons (Fig. 4a and 4b; genotype effect: *F*_1, 60_ = 21.249, *p* < 0.001). Treatment with trifluoperazine specifically reduced the level of puromycin-labeled proteins in *Fmr1* KO neurons to the WT level; it did not affect protein synthesis in WT neurons (Fig. 4; drug effect: *F*_1, 60_ = 2.286, *p* = 0.057; genotype X drug interaction: *F*_5, 60_ = 3.733, *p* < 0.01).

**Figure 4.**
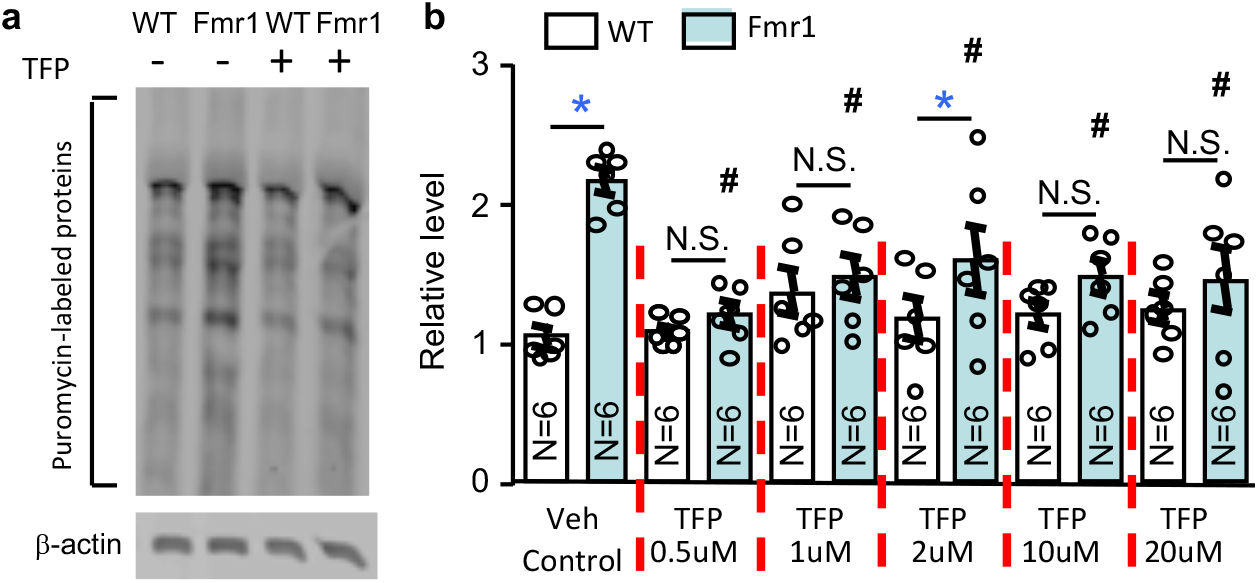
Trifluoperazine corrects enhanced basal translation in *Fmr1* KO neurons. Basal protein synthesis was determined in wild type (WT) and *Fmr1* KO (Fmr1) mouse neurons by the SUnSET method. DIV 14 primary hippocampal neurons were pre-incubated with vehicle (Veh) or trifluoperazine (TFP) (at various concentrations as indicated) for 30 min before the addition of puromycin into the culture media. Equal amounts of protein were loaded on the gel for Western blot analysis with anti-puromycin antibody. (**a**) Representative image shows that the level of puromycin-labeled proteins is higher in *Fmr1* KO than WT neurons, and trifluoperazine dampens protein synthesis in *Fmr1* KO but not WT neurons. (**b**) Quantification shows that the enhanced protein synthesis in *Fmr1* KO but not WT neurons was suppressed by trifluoperazine. The relative level of newly synthesized puromycin-labeled protein in the vehicle-treated WT group was defined as 1, and all samples were normalized to this group. *: *p* < 0.05 between the two genotypes; N.S.: not significant; #: *p* < 0.05 between the TFP- and vehicle-treated *Fmr1* KO group; determined by two-way ANOVA followed by Holm-Sidak test.

### The effect of trifluoperazine is unlikely due to its pharmacological action against dopamine receptors

Regarding the mechanism of action (MOA), trifluoperazine is a phenothiazine derivative and has inhibition activity against dopamine receptors including both D1- and D2-like receptors (D1R/D2R). To directly test the possibility that therapeutic effect of trifluoperazine is mediated through its activity against D1R/D2R, we examined the *in vivo* effects of other well-characterized antagonists on FXS-associated behavior symptoms. In the social interaction test, the D1R antagonist SCH23390 and D2R antagonist raclopride both increased social interaction in WT mice (Fig. 5a; genotype effect: *F*_*1, 58*_ = 121.132, *p* < 0.0001; treatment effect: *F*_*2, 58*_ = 3.447, *p* < 0.05; genotype X treatment interaction: *F*_*2, 58*_ = 2.592, *p* = 0.084) (*p* < 0.05 for SCH23390; *p* < 0.005 for raclopride; *post hoc* Holm-Sidak test). They did not affect *Fmr1* KO mice (*p* = 0.777 for SCH23390, *p* = 0.825 for raclopride; *post hoc* Holm-Sidak test). In the marble bury test, vehicle- and drug-treated *Fmr1* KO mice displayed excessive marble burying activity compared to their corresponding WT groups (for the fully buried marbles in Fig. 5b1; genotype effect: *F*_*1, 83*_ = 18.66, *p* < 0.0001; drug effect: *F*_*2, 83*_ = 5.637, *p* < 0.01; genotype-drug interaction: *F*_*2, 83*_ = 0.107, *p* =0.899) (for the surface marbles in Fig. 5b2; genotype effect: *F*_*1, 83*_ = 47.509, *p* < 0.0001; drug effect: *F*_*2, 83*_ = 11.150, *p* < 0.0001; genotype-drug interaction: *F*_*2, 83*_ = 2.660, *p* = 0.076). SCH23390 non-specifically decreased the % of fully buried marbles in both WT and *Fmr1* KO mice (Fig. 5b1). For % of surface marbles, SCH23390 had effect on WT but not *Fmr1* KO mice (Fig. 5b2). The D2R antagonist raclopride affected neither WT nor *Fmr1* KO mice (Fig. 5b). These data show that the D1R/D2R antagonists have different *in vivo* effects comparing to trifluoperazine. Interestingly, a previous study showed hypo-dopaminergic function in FXS and that a dopamine receptor agonist rescued behavioral abnormality in *Fmr1* KO mice ^31^. Thus, therapeutic effects of trifluoperazine are unlikely mediated through its inhibition action against dopamine receptors.

**Figure 5.**
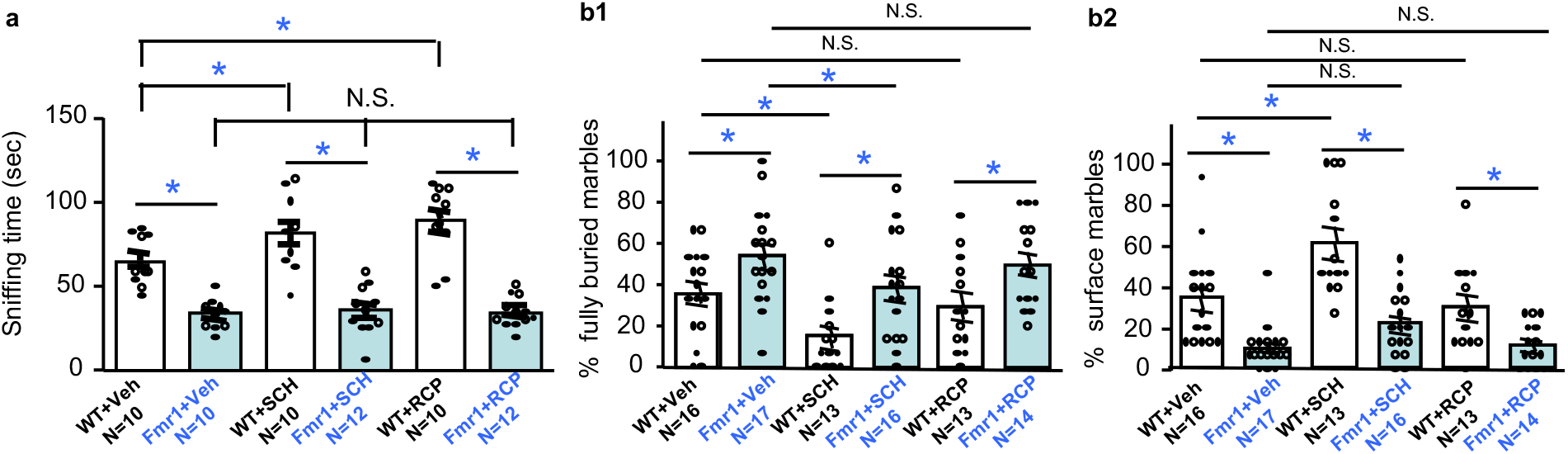
Effects of D1R and D2R antagonists on sociability and repetitive behavior. Wild type (WT) and *Fmr1* KO mice (Fmr1) were i.p. injected with vehicle (Veh), or the D1R antagonist SCH23390 (SCH, 0.05 mg/kg), or the D2R antagonist raclopride (RCP, 0.3 mg/kg). One hour after injection, mice were examined by the 3-chamber social interaction (**a**) or marble bury test (**b**). Direct interaction with the novel stimulus mouse (as indicated by sniffing time) is presented in **a**. For marble bury activity, % of fully buried marble and marbles on the surface is presented in **b1** and **b2**, respectively. *: *p* < 0.05; N.S.: not significant; determined by two-way ANOVA followed by Holm-Sidak test.

### CMAP analysis predicts new pharmacological activities of trifluoperazine

To identify potential new pharmacological activity of trifluoperazine, we screened CMAP for similarity compounds that induce transcriptome alterations overlapping with trifluoperazine-induced changes. Querying CMAP database using trifluoperazine signature in PC3 cell (Supplementary Data 2), which displays oppositional profile to that of *Fmr1* KO neurons (Fig. 1d), revealed compounds with positive and negative similarity scores (Supplementary Table 5). Similarity compounds were ranked in a descending order according to *p* value, enrichment and similarity mean (connectivity/similarity score). Among the top 10 hits, 4 compounds are phenothiazine derivatives and used as typical antipsychotics (Table 1). The other 2 classes of compound over-represented among the top 10 hits are HDAC (histone deacetylase) inhibitors (i.e. vorinostat and trichostatin A) and PI3K signaling inhibitors (i.e. sirolimus and LY-294002). Further, another well-known PI3K inhibitor wortmannin also triggers transcriptome changes that significantly overlap with trifluoperazine-induced changes. Wortmannin-induced gene signatures in different cell lines are ranked at 35 (MCF7 cell, *p*=0.00038, enrichment=0.608, and similarity mean=0.244) and 74 (PC3 cell, *p*=0.00455, enrichment=0.95, and similarity mean=0.389), respectively (Supplementary Table 5). These results predict an intriguing possibility that trifluoperazine may process new pharmacological activity as HDAC and/or PI3K signaling inhibitor.

**Table 1.** Top 10 trifluoperazine similarity compounds. Signature probes of trifluoperazine in PC3 cell are used as query to search similarity compounds in CMAP database. Compound with positive and negative “similarity mean” triggers overlapping and oppositional transcriptome changes, respectively, comparing to the change caused by trifluoperazine in PC3 cell. Rank, p value, enrichment score, and similarity mean are described in Supplementary Table 4. Compound name and cell line indicate the name of compound used for treatment with specific cell lines in CMAP database. Number of arrays (N) is the number of analyzed arrays obtained from all instances of the corresponding compound. MOA: mechanism of action.

| Rank | Compound name and cell line | Similarity mean | N | Enrich -ment | P value | Application/MOA |
| --- | --- | --- | --- | --- | --- | --- |
| 1 | fluphenazine - PC3 | 0.558 | 3 | 0.997 | 0 | typical antipsychotics/derivatives of phenothiazine |
| 2 | thioridazine - PC3 | 0.569 | 5 | 0.994 | 0 | typical antipsychotics/derivatives of phenothiazine |
| 3 | vorinostat - MCF7 | 0.333 | 7 | 0.92 | 0 | HDAC inhibitor |
| 4 | sirolimus - PC3 | 0.344 | 8 | 0.871 | 0 | PI3K/Akt/mTOR inhibitor |
| 5 | prochlorperazine - MCF7 | 0.369 | 9 | 0.867 | 0 | typical antipsychotics/derivatives of phenothiazine |
| 6 | trichostatin A - PC3 | 0.347 | 55 | 0.862 | 0 | HDAC inhibitor |
| 7 | trifluoperazine - MCF7 | 0.371 | 9 | 0.787 | 0 | typical antipsychotics/derivatives of phenothiazine |
| 8 | geldanamycin - MCF7 | 0.294 | 10 | 0.753 | 0 | HSP inhibitor |
| 9 | LY-294002 - PC3 | 0.315 | 12 | 0.748 | 0 | PI3K/Akt/mTOR inhibitor |
| 10 | trichostatin A - MCF7 | 0.29 | 92 | 0.729 | 0 | HDAC inhibitor |

### Trifluoperazine inhibits PI3K-Akt-S6K1 signaling cascade but does not affect histone acetylation in neurons

We first examined the effect of trifluoperazine on histone acetylation. We confirmed that treatment with vorinostat (Fig. 6a) or trichostatin A (Fig. 6b) increased acetylation of histone H3 and H4. We did not observe any significant effects of trifluoperazine on these readouts of HDAC activity (Fig. 6c).

**Figure 6.**
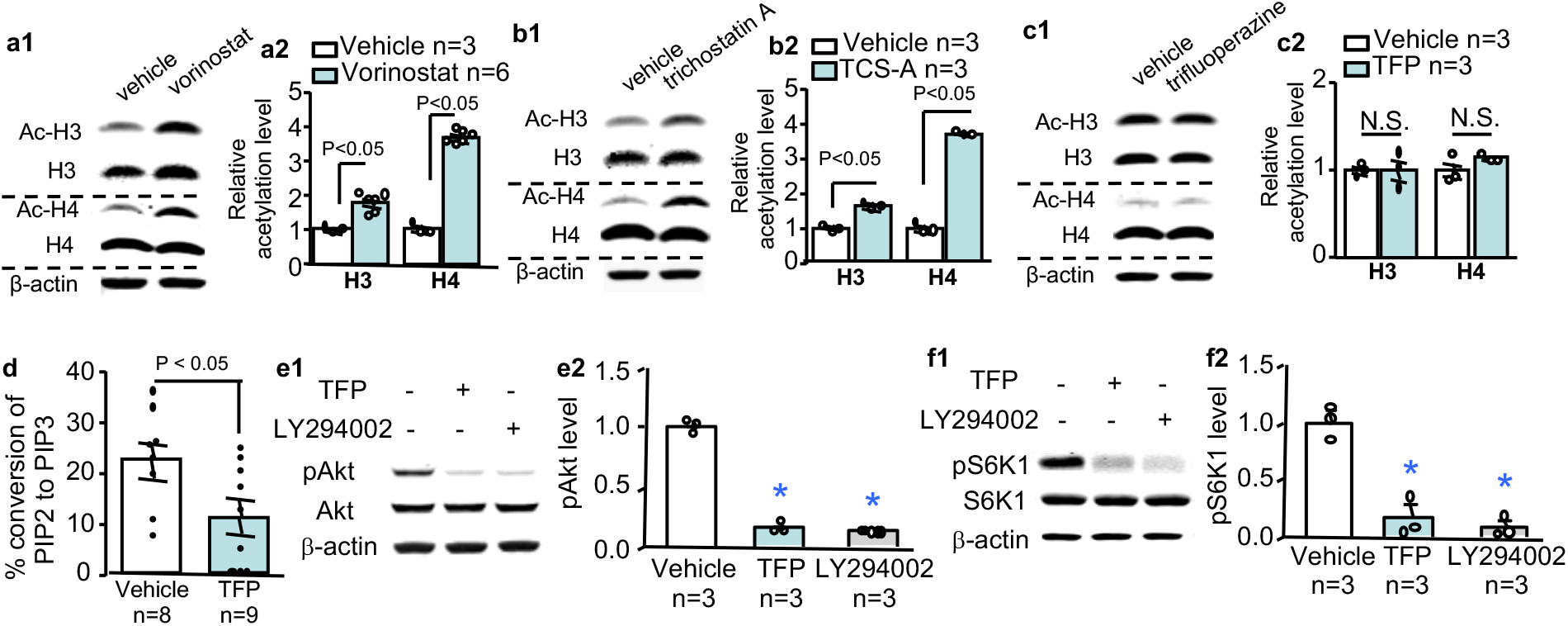
Trifluoperazine suppresses the PI3K-Akt-S6K1 signaling cascade but does not affect histone acetylation in hippocampal neurons. (**a** to **c**) Hippocampal neurons cultured from WT mice were treated with vorinostat (20 μM) (**a**), trichostatin A (20 μM) (**b**), and trifluoperazine (20 μM) (**c**) for 1 hour, and then harvested for Western blot analysis. Level of the acetylated histone H3 and H4 (Ac-H3 and Ac-H4) in vehicle- and drug-treated sample was normalized to total H3 and H4, respectively. (**d**) PI3 kinase activity in lysates from primary WT hippocampal neurons treated with 20 μM trifluoperazine (TFP) or vehicle was determined by ELISA, which measures the concentration of PIP3 converted from PIP2. (**e** and **f**) Primary WT hippocampal neurons were treated with trifluoperazine (TFP) (20 μM) or LY-294002 (20 μM) for 1 h. Samples harvested immediately after drug or vehicle treatment were analyzed by Western blot for the level of pAkt (normalized to total Akt) (**e**) and pS6K1 (normalized to total S6K1) (**f**). The relative protein level in the vehicle-treated control group was defined as 1 (**a2**, **b2**, **c2**, **e2**, and **f2**). P value was determined by student’s t-test (**a** to **d**) or one-way ANOVA and *post hoc* Tukey’s test (**e** and **f**). *: *p* < 0.05 between the control/vehicle and the indicated group. N.S.: not significant.

We next examined the effect of trifluoperazine on PI3K, elevated activity of which has been found in FXS samples ^32, 33^. Enzymatic assay demonstrates that trifluoperazine significantly suppresses PI3K activity in hippocampal neurons (Fig. 6d). We further examined the major PI3K downstream target Akt, and found that, similar to LY-294002, trifluoperazine also dampened Akt activity, as indicated by reduction of phosphorylation (Fig. 6e, treatment effect: *F*_2, 6_ = 1200.316, *p* < 0.001, one-way ANOVA). We next examined the activity of S6K1, which has been implicated as PI3K/Akt downstream target and involved in regulating protein translation machinery and hyper-active in *Fmr1* KO mice and FXS patients ^15, 34, 35^. We found that both LY-294002 and trifluoperazine suppressed S6K1 phosphorylation at Thr389 (a target site of PI3K ^36, 37^) (Fig. 6f, treatment effect: *F*_2, 6_ = 37.441, *p* < 0.001, one-way ANOVA).

One complication is that, as implicated in mesangial cells, long-term trifluoperazine treatment has been shown to cause cell death, which may indirectly affect the activity of survival-supporting molecules such as Akt ^38^. In our experimental set-up, 1-hour trifluoperazine treatment did not affect neuronal viability (Supplementary Fig. 10; genotype effect: *F*_1, 39_ = 3.547, *p* = 0.067; trifluoperazine effect: *F*_2, 39_ = 1.580, *p* = 0.219; genotype X trifluoperazine interaction: *F*_2, 39_ = 0.024, *p* = 0.976). These data demonstrate that trifluoperazine processes a new and previously undescribed MOA against the PI3K-Akt-S6K1 signaling cascade in neurons.

### Trifluoperazine normalizes the aberrantly elevated Akt-S6K1 signaling in *Fmr1* KO mice

Previous studies have shown that enhanced Akt-S6K1 signaling may be causal for certain aspects of symptoms in *Fmr1* KO mice ^15, 39^. We confirmed that, with comparable levels of total Akt (Fig. 7a and 7b1; genotype effect: *F*_1, 20_ = 0.460, *p* = 0.505; drug effect: *F*_1, 20_ = 1.041, *p* = 0.320; genotype X drug interaction: *F*_1, 20_ = 0.407, *p* = 0.531) and S6K1 (Fig. 7a and 7c1; genotype effect: *F*_1, 20_ = 0.044, *p* = 0.836; drug effect: *F*_1, 20_ = 0.140, *p* = 0.713; genotype X drug interaction: *F*_1, 20_ = 0.045, *p* = 0.834), the levels of pAkt (Fig. 7a and 7b2, genotype effect: *F*_*1, 20*_ = 27.793, *p* < 0.0001) and pS6K1 (Fig. 7a and 7c2, genotype effect: *F*_*1, 20*_ = 79.221, *p* < 0.0001) are elevated in the hippocampus of *Fmr1* KO mice. Administration of trifluoperazine (0.05 mg/kg) in *Fmr1* KO mice normalized the elevated pAkt (Fig. 7a and 7b2; drug effect: *F*_*1, 20*_ = 25.791, *p* < 0.0001; genotype X drug interaction: *F*_*1, 20*_ = 22.512, *p* < 0.0001) and pS6K1 (Fig. 7a and 7c2; drug effect: *F*_*1, 20*_ = 40.388, *p* < 0.0001; genotype X drug interaction: *F*_*1, 20*_ = 35.893, *p* < 0.0001) to the WT level. Together, these results demonstrate that trifluoperazine can normalize the aberrantly elevated PI3K-Akt-S6K1 signaling in *Fmr1* KO neurons. Further, Akt activity (as indicated by pAkt) in *Fmr1* KO neurons is more sensitive to trifluoperazine than in WT neurons (Fig. 8). We found that trifluoperazine at 0.25, 0.5, and 1 μM dampened pAkt in *Fmr1* KO (Fig. 8b) but not in WT neurons (Fig. 8a). Akt activity in WT neurons was significantly suppressed by 2 and 10 μM trifluoperazine (Fig. 8a).

**Figure 7.**
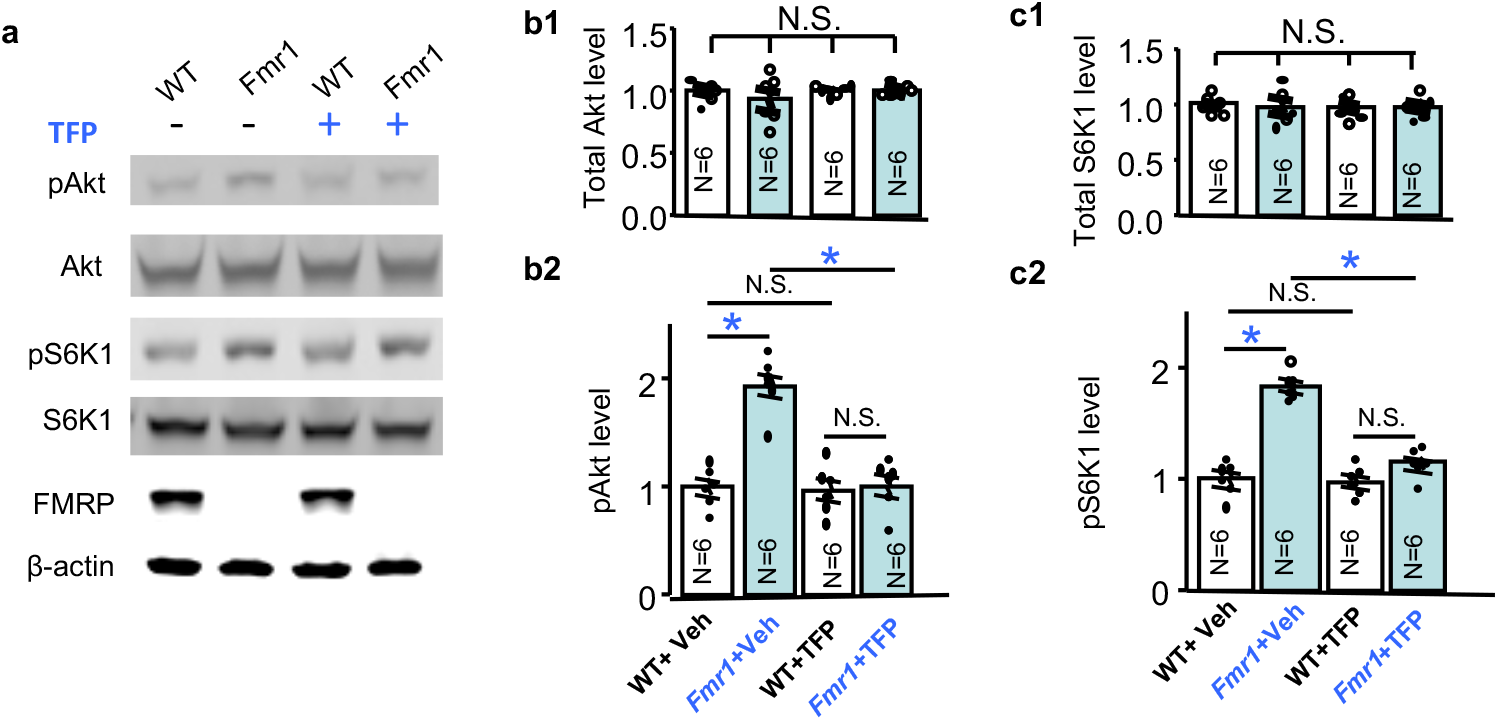
Trifluoperazine (TFP) corrects the aberrantly elevated Akt and S6K1 activity in *Fmr1* KO mice. Wild type (WT) and *Fmr1* KO (Fmr1) mice were injected with vehicle or trifluoperazine (0.05 mg/kg) 1 hour before hippocampus was harvested. The levels of phosphorylated Akt (pAkt), total Akt, phosphorylated S6K1 (pS6K1 at Thr389), and total S6K1 (**a**) were determined by Western blot. Protein loading was determined by the level of β-actin (**a**). The expression of FMRP in WT and *Fmr1* KO samples was confirmed by Western blot (**a**). Representative images are shown in **a**. For quantification, the level of total Akt (**b1**) and total S6K1 (**c1**) was normalized to the level of β-actin. The level of pAkt (**b2**) and pS6K1 (**c2**) was normalized to the level of total Akt and total S6K1, respectively. The relative level of Akt (or pAkt) and S6K1 (or pS6K1) in the vehicle-treated wild type (WT) group was defined as 1, and all samples were normalized to this group. *: *p* < 0.05; N.S.: not significant; determined by two-way ANOVA followed by Holm-Sidak test.

**Figure 8.**
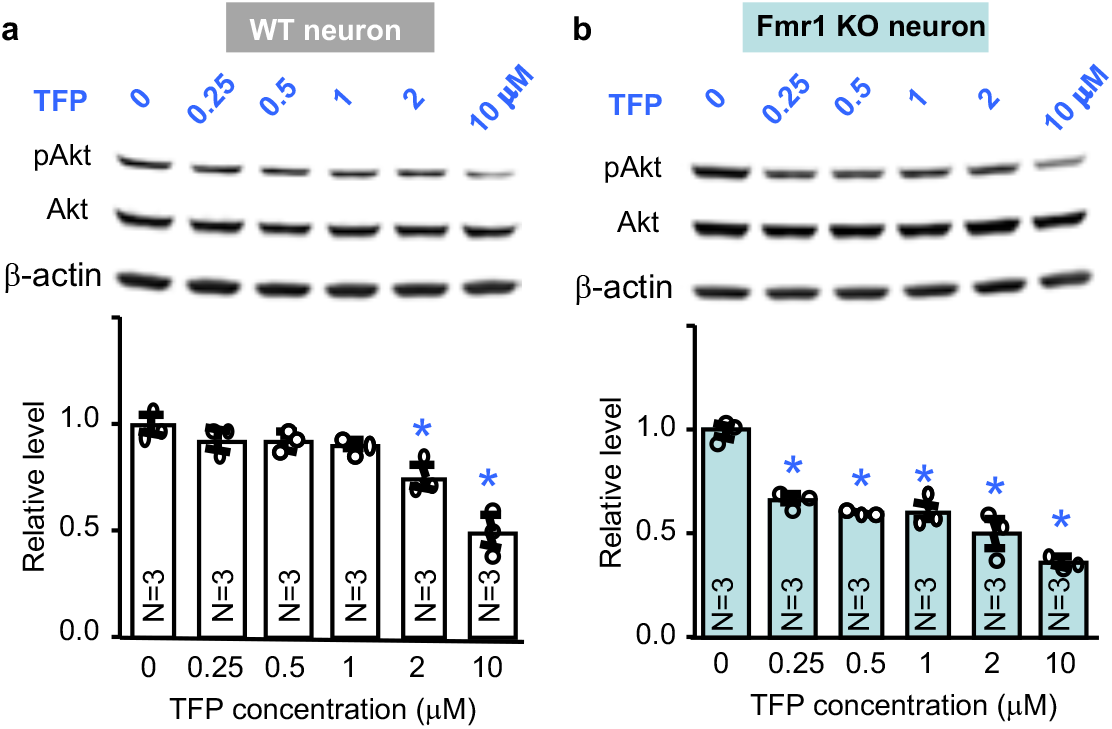
*Fmr1* KO neurons are more sensitive to trifluoperazine treatment. DIV 14 wild type (WT) (**a**) and *Fmr1* KO neurons (**b**) were treated with trifluoperazine (TFP) at various concentrations (as indicated). Following the 1 h treatment, levels of pAkt, total Akt, and β-actin were determined by Western blot. The level of pAkt is normalized to the level of total Akt. The relative level of pAkt in the vehicle-treated control group is defined as 1, and all samples are normalized to this group. *: *p* < 0.05 between the indicated group and the vehicle-treated control group; determined by one-way ANOVA followed by *post hoc* Tukey’s test. *F*_*5, 12*_ = 17.777, *p* < 0.0001 for **a**; *F*_*5, 12*_ = 33.545, *p* < 0.0001 for **b**.

## DISCUSSION

Compared to *de novo* therapeutic development, repurposing FDA-approved drugs offers several benefits including reduction of cost and potentially sooner clinical application. Regarding examination of FDA-approved drugs for FXS treatment, recent studies mainly focused on the known disease mechanism and used hypothesis-driven approaches. For instance, therapeutic testing with metformin and lovastatin, which show promising efficacy in both FXS mouse model ^40, 41^ and small scale open-label human studies ^42, 43^, is theoretically based on their pharmacological actions against mTORC1 and ERK1/2 (extracellular signal-regulated kinase 1/2), the activity of which is abnormally elevated in FXS mouse model and patients ^3^. In this study, we used whole transcriptome analysis as an unbiased approach to identify potential therapeutics. Gene signature analysis with normal neurons and psychiatric disorder-affected specimens suggests an intriguing possibility that transcriptome landscape and cell function are associated and may mutually affect each other. Related to disease mechanism, genomic alteration is considered as an emerging molecular phenotype and pathological endpoint. Intriguingly, recent computational comparison of gene signatures associated with psychiatric disorders and drug treatment suggests an attractive but yet to-be-validated approach for therapy development and drug repurposing ^7, 8^. Here, we compared transcriptome alterations caused by *Fmr1* deficiency and CMAP drugs. The computational analysis identified trifluoperazine as a potential therapeutic. It is important to note that transcriptome data in the CMAP database were collected from cancer cell lines rather than neurons. Interestingly, we were able to validate the CMAP-predicted effect of trifluoperazine in neurons and FXS mouse model. Two recent computational studies also found that the CMAP data from cancer cells may be effectively used to predict therapy or mechanism for neurological disorders ^7, 8^.

Trifluoperazine, belonging to the phenothiazine group, is a well-known antipsychotic. Interestingly, some antipsychotics such as haloperidol, risperidone, and aripiprazole have been noted to alleviate irritability and aggression in autism and FXS patients ^44–46^. The main side effects include extrapyramidal symptoms (EPS) and sedation. In particular for trifluoperazine, long-term use in the 6 to 40 mg/day range exerts EPS ^24^. However, in theory, side effects depend on drug dose, and human studies demonstrate that EPS can be successfully managed through dose adjustment. Here, we observed robust efficacy of low dose trifluoperazine at 0.05 mg/kg, which, according to the well-accepted “FDA formula” ^47^, is equivalent to 0.004 mg/kg for human (or 0.24 mg for a person with 60 kg body weight). This dose is highly applicable and tolerated, as it is significantly lower than 5 mg/day, which, judged by the Extrapyramidal Symptom Rating Scale, does not cause adverse side effects in humans ^25^. Notably, low dose trifluoperazine (i.e. 1 mg, which is about 4X higher than 0.24 mg) is also used in combination with antidepressant in human patients, and sold under the brand names of Parmodalin and Jatrosom N.

The primary application of trifluoperazine in schizophrenia patients shows strong effects on correcting the positive symptoms possibly through its inhibition on dopamine receptors. It is known that the anti-dopamine effects of trifluoperazine decrease locomotor activity but does not improve cognitive function. We found that trifluoperazine at 0.05 mg/kg does not affect locomotor activity in WT mice in the light-dark and open field test. In *Fmr1* KO mice, trifluoperazine normalizes the excessive locomotor transition in the light-dark test but not in the open field paradigm. Notably, 0.05 mg/kg trifluoperazine rescued the cognitive impairment in *Fmr1* KO mice. Interestingly, certain aspects of FXS-associated abnormality are caused by decreased dopamine function and dopamine agonist corrects certain phenotypes in FXS mouse ^31^. We further examined the effect of other known D1R and D2R antagonists, and found no therapeutic efficacy in correcting FXS-associated abnormalities. These lines of evidence suggest that the therapeutic effects of low dose trifluoperazine are unlikely due to its anti-dopamine activity.

Our study further used computational screening with CMAP to identify a potential pharmacological mechanism underlying the therapeutic effects of trifluoperazine. Intriguingly, although gene signature induced by trifluoperazine shows similarity to those induced by the known HDAC and PI3K inhibitors, we found trifluoperazine effect on PI3K activity but not histone acetylation. However, as our assay only examined acetylation of H3 and H4 at specific lysine residuals (i.e. K9 of H3 and K8 of H4), more exhaustive investigation may be needed to totally exclude potential trifluoperazine activity against HDAC. Since all available small molecule drugs have promiscuous pharmacological targets, we do not exclude the possibility that the therapeutic effects of trifluoperazine are mediated by regulating other signaling pathways in addition to the inhibition of the PI3K-Akt-S6K1 cascade. Nevertheless, our results do suggest that drugs with similar effects on transcriptome signature may have similar MOA (mechanism of action) and warrant experimental validation.

Notably, some of the top-ranked trifluoperazine similarity compounds also induce transcriptome alterations that are oppositional to that caused by *Fmr1* deficiency. For example, gene signatures of trichostatin A (in PC3 cell, ranked at #6) and thioridazine (in PC3 cell, ranked at #2) show positive similarity to trifluoperazine signature (Table 1). They show negative similarity to *Fmr1* KO neuron signature, and are ranked at #1 and #9, respectively (Fig. 1d). Although these compounds are either not approved by FDA or withdrawn from clinical use, it would be interesting to test whether they can correct certain pathological aspects of FXS in animal models. Along the same line, our results advocate future studies to repurpose other CMAP-identified trifluoperazine similarity drugs (Supplementary Table 5).

Intriguingly, our unbiased computational prediction of trifluoperazine efficacy and its action against PI3K activity are strikingly coincident with a previously suggested disease mechanism. Recent studies have found that, in the absence of FMRP, expression levels of PI3K activator PIKE and p110ß subunit are elevated in *Fmr1* deficient cells ^32, 34^, leading to elevated PI3K activity and over-activation of the Akt-S6K1 signaling cascade ^15, 35^. Therapeutic value of correcting the altered PI3K-Akt-S6K1 signaling in FXS is strongly suggested by that genetic removal/reduction of S6K1, PIKE, and p110ß subunit of PI3K can rescue the elevated overall protein synthesis and multiple behavioral abnormalities in *Fmr1* KO mice ^15, 39, 48^. A more recent study showed that systemic injection of a novel p110β-specific inhibitor GSK2702926A, a similarly structured compound of which is under investigation in a cancer clinical trial, rescues cellular and behavioral abnormalities in *Fmr1* KO mice ^49^. To realize therapeutic value of targeting the PI3K-Akt-S6K1 signaling cascade, a significant advancement is to find a clinically suitable approach to effectively dampen PI3K-Akt-S6K1 activity in FXS. By using the FDA-approved trifluoperazine, we demonstrate that acute administration of a drug with PI3K inhibiting activity is sufficient to rescue multiple aspects of FXS-associated pathology, supporting the practical value of targeting PI3K-Akt-S6K1 signaling. Considering that human FXS samples also display elevated PI3K-Akt-S6K1 signaling ^33, 35^, our data encourage future investigation to determine the efficacy of trifluoperazine in human patients.

We acknowledge that trifluoperazine does not correct all FXS-associated symptoms that were examined in this study. Lack of “absolute efficacy” is common for drugs used for complex neurological disorders. It is noticed that targeting ERK1/2 or PI3K by lovastatin, metformin, and GSK2702926A rescues some but not all symptoms ^40, 41, 49^. Such limitation may be due to multiple pathological alterations in FXS; combinatorial targeting multiple disease factors represents an emerging idea to achieve more robust therapeutic outcome ^50^. Alternatively, adjustment of dose and treatment duration may also be considered for future studies. Furthermore, although FXS is a main cause for autism and intellectually disability (ID), we do not expect that trifluoperazine would show general therapeutic effects in other autism and ID models.

We acknowledge that FDA has just approved the first PI3K inhibitor idelalisib in July 2014 for the treatment of leukemia. Idelalisib (not included in CMAP database) shows high specificity against the p110δ isoform of PI3K, which is expressed mainly in hemocyte and marginally in limited brain regions (according to Allen Brain Atlas). Considering that the dysfunction in FXS involves numerous brain structures, the therapeutic efficacy of idelalisib may be limited. Another complication is that its pharmacokinetic profile has only been examined for blood absorption and excretion. Whether idelalisib can cross the blood brain barrier has not been determined and not known ^51^.

In summary, our study supports the value of holistic transcriptome analysis in therapeutic discovery and repurposing FDA-approved drugs for FXS treatment. Experimental validation of trifluoperazine efficacy in FXS mouse model suggests a practical treatment strategy. Our results along with a recent computational study ^8^ also encourage broader application of transcriptome screening with public databases in guiding drug discovery for other neurological disorders.

## Acknowledgements

We thank Drs. James Malter (UT Southwestern Medical Center) and Cara Westmark (University of Wisconsin-Madison) for providing the *Fmr1* KO mice.

## Author contributions

H.W. initiated the study. Q.D., F. S., W.F., Y.F., and H.W. designed the experiments and analyzed the data. X.W., Z.M., P.C., Y.Z., H.X., and X.L. performed bioinformatics analysis. H.W., Q.D., and F.S. wrote the paper.

---

**Supplementary Figure 1.**
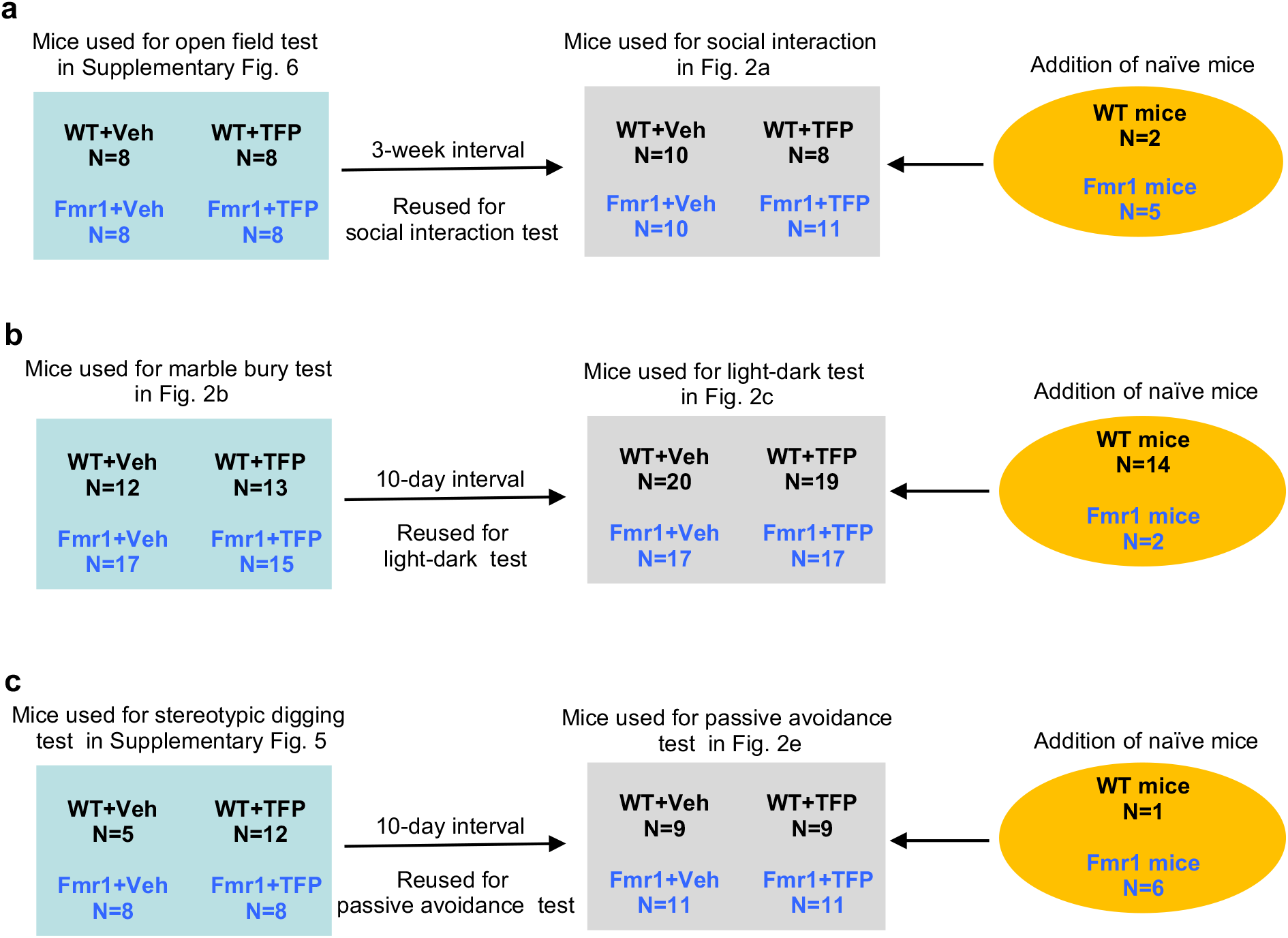
Animals that are reused for different behavioral tests. **a.** Wild type (WT) and *Fmr1* knockout (Fmr1) mice were first used for open field test, and then reused for social interaction test with the addition of naïve mice. **b.** WT and *Fmr1* mice were first used for marble bury test, and then reused for light-dark test with the addition of naïve mice. **c.** WT and *Fmr1* mice were first used for stereotypic digging test, and then reused for passive avoidance test with the addition of naïve mice. Veh: vehicle. TFP: trifluoperazine.

**Supplementary Figure 2.**
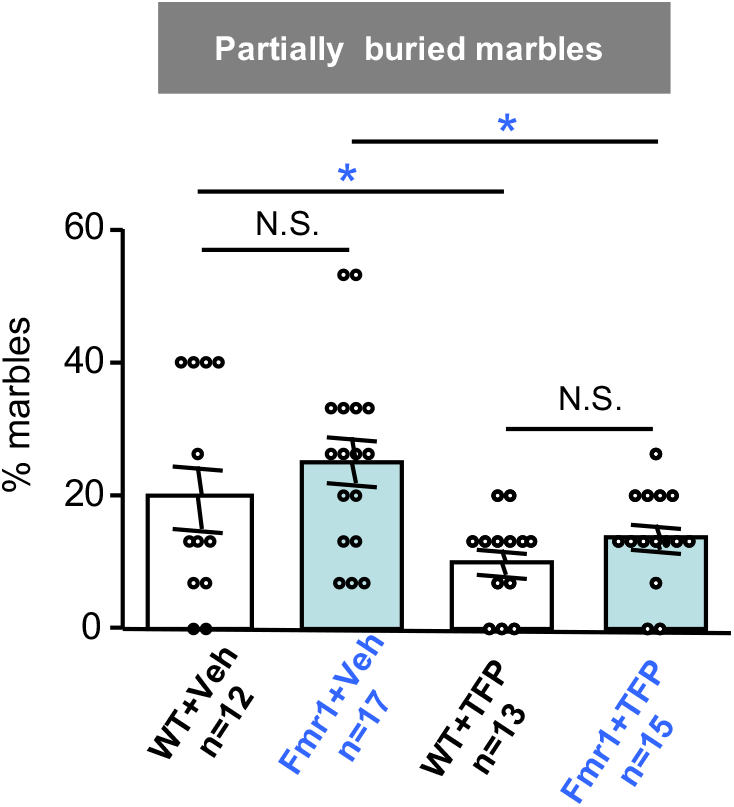
Trifluoperazine (TFP) non-specifically decreases partially buried marbles in wild type (WT) and *Fmr1* KO (Fmr1) mice. Veh: vehicle. *: *p* < 0.05; N.S.: not significant; determined by two-way ANOVA followed by Holm-Sidak test.

**Supplementary Figure 3.**
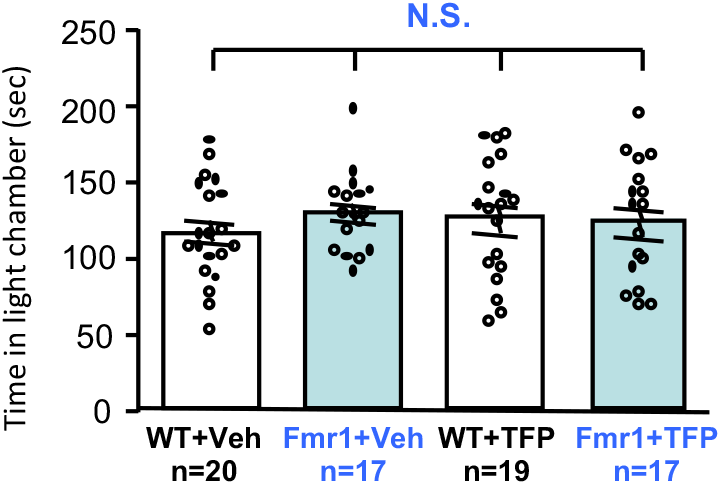
Wild type (WT) and *Fmr1* KO (Fmr1) mice spent similar time in the lit chamber during the light-dark test. Vehicle (Veh)- or trifluoperazine (TFP)-injected mice were subjected to light-dark test for 5 min. Time spent in the lit chamber was scored. N.S.: not significant; determined by two-way ANOVA.

**Supplementary Figure 4.**
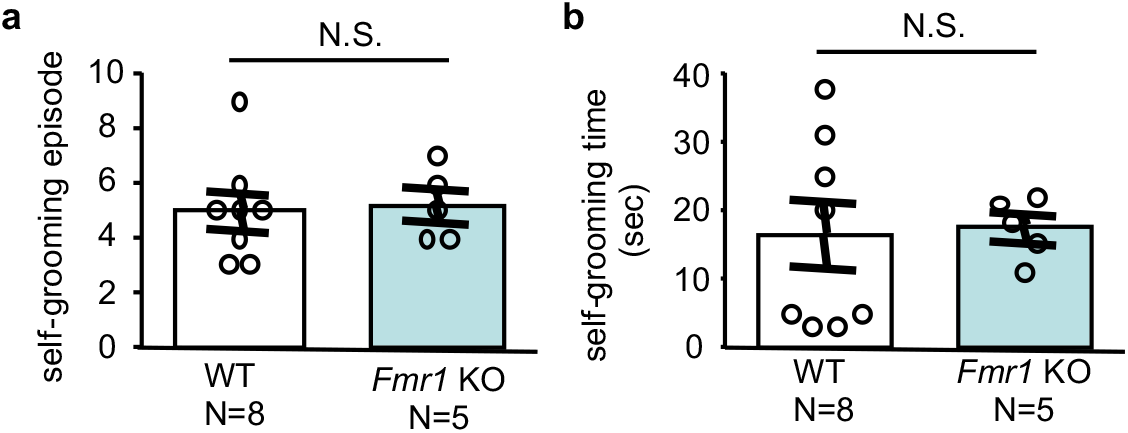
Wild type (WT) and *Fmr1* knockout (KO) mice show comparable self-grooming. WT and KO mice were examined for self-grooming. The number of grooming session (i.e. episode) is presented in **a**. Total grooming time is presented in **b**. N.S.: not significant, determined by Student’s t-test.

**Supplementary Figure 5.**
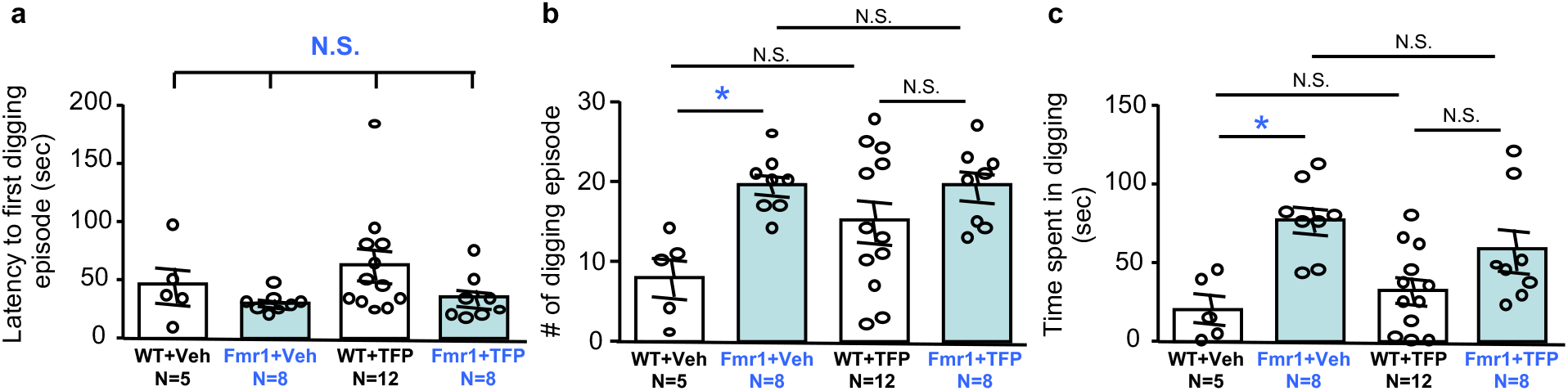
Trifluoperazine (TFP) does not significantly affect stereotypic digging behavior. (**a**) Wild type (WT) and *Fmr1* KO (Fmr1) mice show comparable latency to the first digging behavior, and are not affected by TFP. Genotype effect: *F*_1, 29_ = 3.395, *p* = 0.076; drug effect: *F*_1, 29_ = 0.783, *p* = 0.383; genotype-drug interaction: *F*_1, 29_ = 0.305, *p* = 0.585. (**b**) *Fmr1* KO mice show more digging episode, and are not affected by TFP. Genotype effect: *F*_1, 29_ = 11.264, *p* < 0.005; drug effect: *F*_1, 29_ = 2.005, *p* = 0.167; genotype-drug interaction: *F*_1, 29_ = 2.313, *p* = 0.139. (**c**) *Fmr1* KO mice show more total digging time, and are not affected by TFP. Genotype effect: *F*_1, 29_ = 15.809, *p* < 0.001; drug effect: *F*_1, 29_ = 0.138, *p* = 0.713; genotype-drug interaction: *F*_1, 29_ = 2.319, *p* = 0.139. Veh: vehicle. *: *p* < 0.05; N.S.: not significant; determined by two-way ANOVA followed by Holm-Sidak test.

**Supplementary Figure 6.**
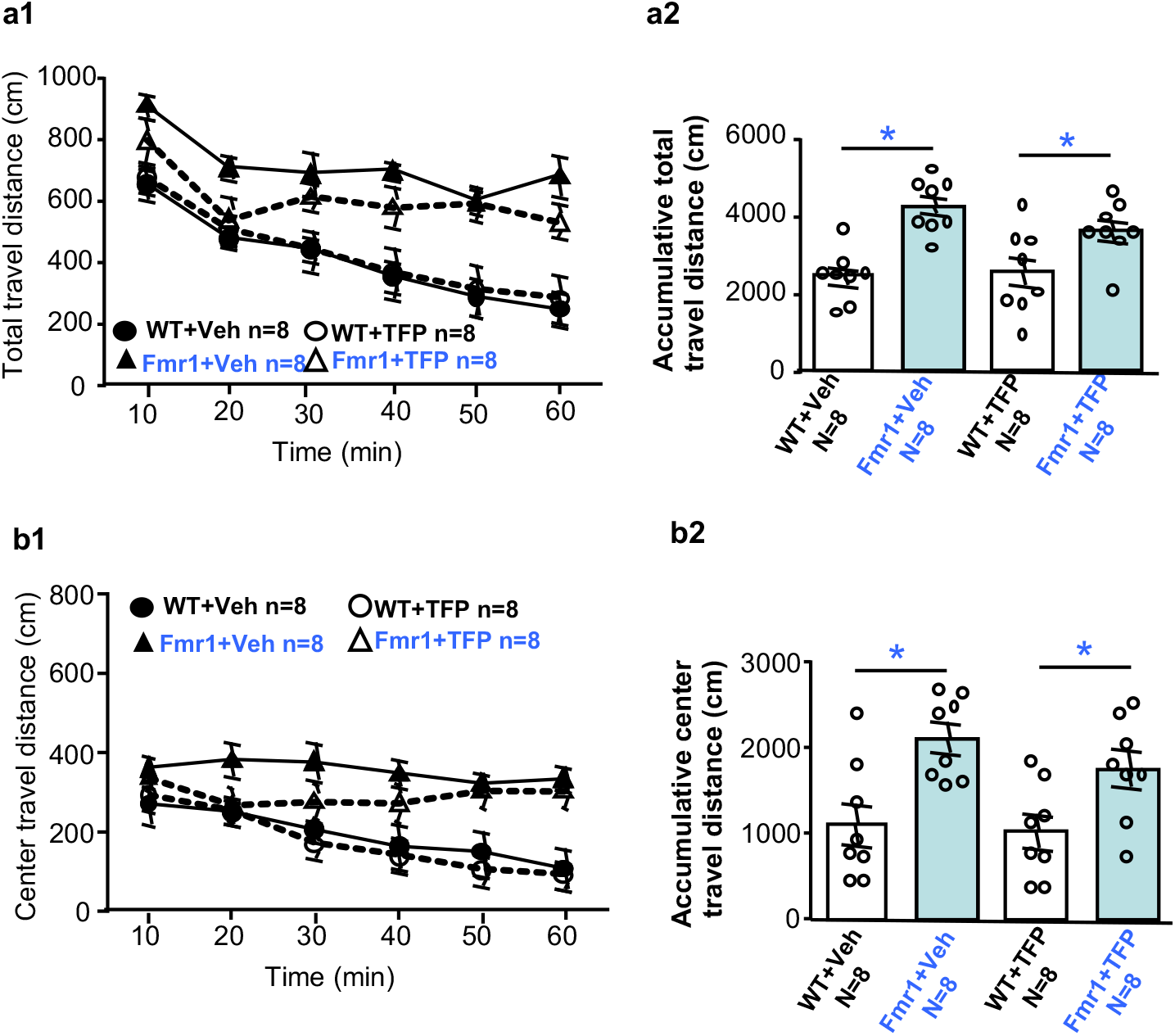
Trifluoperazine (TFP) has no effect on locomotor activity in open field. One hour after trifluoperazine administration, mice were subjected to an open field test. During the 1-hour test, ambulatory travel distance in the whole arena (**a**) or in the center area (**b**) for each of the 10-min bin (**a1** and **b1**) and during the whole 1-hour testing (i.e. accumulative travel distance in **a2** and **b2**) are presented. Veh: vehicle. *: *p* < 0.05; determined by two-way ANOVA followed by Holm-Sidak test.

**Supplementary Figure 7.**
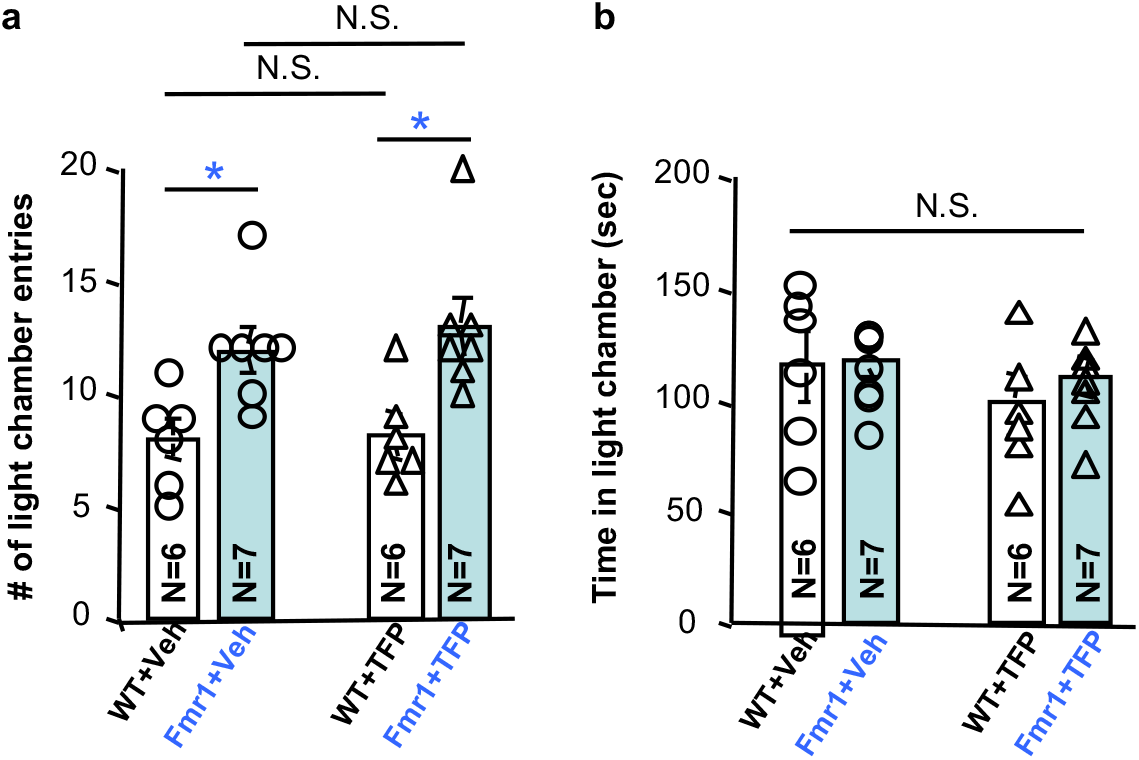
The effect of trifluoperazine on locomotion does not persist beyond 24 hours after drug administration. Wild type (WT) and *Fmr1* KO (Fmr1) mice were injected with trifluoperazine (TFP) (0.05 mg/kg) or vehicle (Veh). 24 hours later, these four groups of mice were subjected to light-dark test. The number of locomotive transition between the lit and dark chamber is presented in **a**. *: *p* < 0.05; N.S.: not significant; determined by two-way ANOVA followed by Holm-Sidak test. Total time stayed in the lit chamber is presented in **b**. N.S.: no significant difference among these four groups. Genotype effect: *F*_1, 22_ = 0.213, *p* = 0.649; drug effect: *F*_1, 22_ = 1.307, *p* = 0.265; genotype X drug interaction: *F*_1, 22_ = 0.552, *p* = 0.466; determined by two-way ANOVA.

**Supplementary Figure 8.**
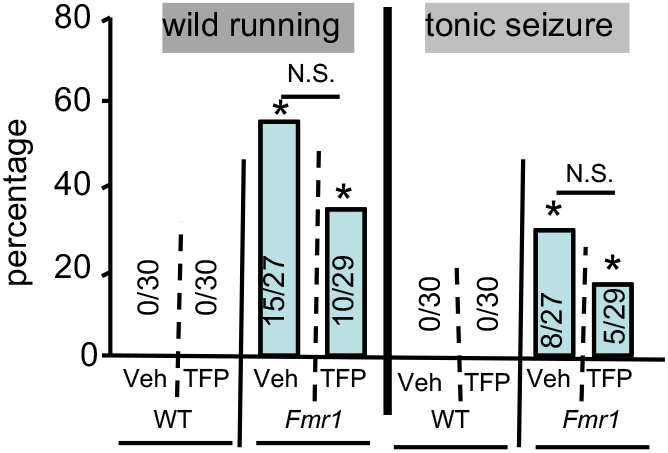
Trifluoperazine does not significantly affect audiogenic seizures. Audiogenic seizures were tested with 21- to 24-day old mice. The percentage of animals showing wild running and seizures is presented for wild type (WT) and *Fmr1* KO mice (*Fmr1*) receiving vehicle (Veh) or trifluoperazine (TFP). *: *p* < 0.05 between genotypes (determined by Fisher’s exact test); N.S.: not significant (determined by Chi-square analysis).

**Supplementary Figure 9.**
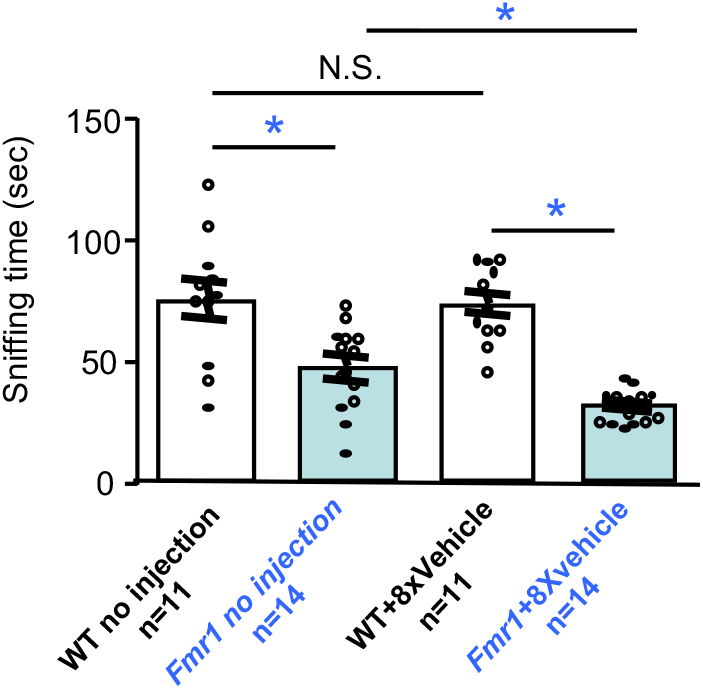
The repeated injection procedure reduces sociability in *Fmr1* KO mice. Wild type (WT) and *Fmr1* KO (Fmr1) mice receiving no injection or 8 daily injection of vehicle were examined in the 3-chamber social interaction test. Time spent in sniffing the novel stimulus mouse enclosure was compared. Regardless of the injection procedure, WT mice show higher sociability than *Fmr1* KO mice (genotype effect: *F*_1, 46_ = 49.614, *p* < 0.001; injection effect: *F*_1, 46_ = 3.117, *p* = 0.084; genotype X injection interaction: *F*_1, 46_ = 1.971, *p* = 0.167). *: *p* < 0.05; N.S.: not significant; determined by two-way ANOVA followed by *post-hoc* Holm-Sidak test.

**Supplementary Figure 10.**
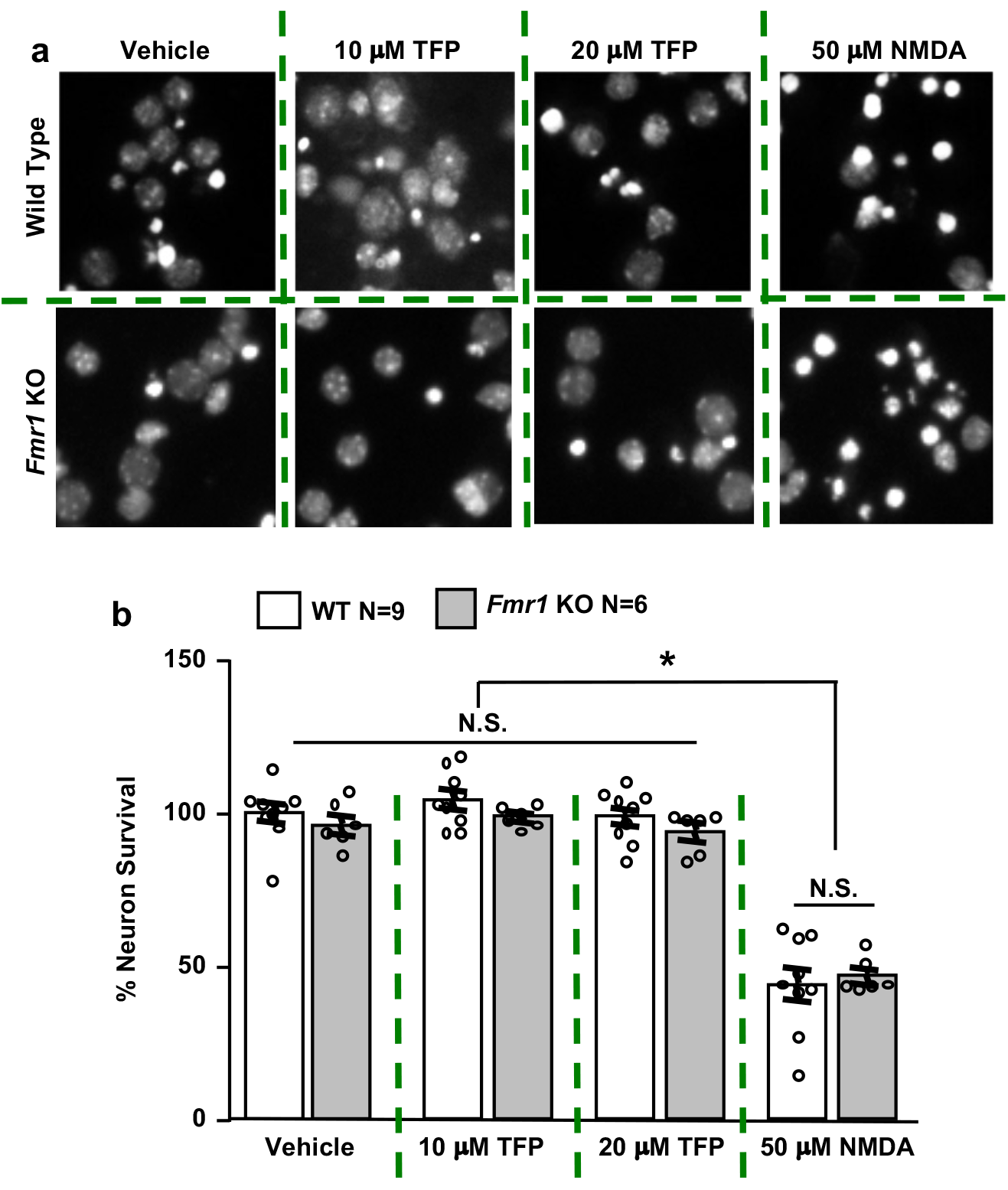
Trifluoperazine (TFP) does not affect neuron viability. DIV 14 wild type (WT) and *Fmr1* KO hippocampal neurons were treated with vehicle, 10 μM TFP, 20 μM TFP, or 50 μM NMDA. Representative DAPI staining is shown in **a**. Quantification of viability is shown in **b**. N.S.: not significant, determined by two-way ANOVA between vehicle and TFP-treated groups. When NMDA treatment is included, there is significant NMDA effect on neuron viability (*: *p* < 0.05; N.S.: not significant; determined by two-way ANOVA followed by Holm-Sidak test).

## Supplementary Table Legends

**Supplementary Table 1** Complete list of GO biology processes associated with differentially expressed genes (DEG) in Fmr1 knockout hippocampal neurons. The list of GO categories is arranged in an ascending order of P values (P<0.05 is considered significant). Count represents the number of DEG detected in the specified GO process. % represents the percentage of total number of genes involved in the specified GO process.

**Supplementary Table 2** Genes in the top 15 GO processes associated with DEGs in Fmr1 knockout (KO) hippocampal neurons. Changes of transcript level (ratio of *Fmr1* KO to wild type value) are expressed as Log2 values.

**Supplementary Table 3** Genes in the KEGG pathways associated with differentially expressed genes (DEG) in *Fmr1* knockout (KO) hippocampal neurons. KEGG pathways are ranked and listed in an ascending order of P values (P < 0.05 is considered significant, as shown in Fig. 1C). Changes of transcript level (ratio of *Fmr1* KO to wild type value) are expressed as Log2 values.

**Supplementary Table 4** Ranking of CMAP compounds, which trigger similar transcriptome alterations (including positive and negative correlations) to that in *Fmr1* knockout hippocampal neurons. Ranking is arranged in an ascending order of p-value, and then a descending order of (absolute) enrichment when two p-values are identical. Compound name and cell line indicate the name of compound used for treatment with specific cell lines in CMAP database. The similarity mean is the arithmetic mean of the similarity/connectivity scores for all instances of the specific named compound. Number of arrays (N) is the number of analyzed arrays obtained from all instances of the corresponding compound/drug. Enrichment score is a measure of enrichment of DEG between *Fmr1* KO samples and all instances of the specific CMAP compound. It is computed using the Kolmogorov-Smirnov statistic as described 38. Compound with negative similarity mean triggers oppositional transcriptome changes comparing to the change caused by FMRP deficiency in Fmr1 knockout neurons. Compound with positive similarity mean triggers transcriptome changes overlapping with that caused by FMRP deficiency in *Fmr1* knockout neurons.

**Supplementary Table 5** Ranking of CMAP compounds, which trigger similar transcriptome alterations (including positive and negative correlations) to that triggered by trifluoperazine. Signature probes of trifluoperazine in PC3 cell are used as query. Compound with positive and negative similarity mean triggers overlapping and oppositional transcriptome changes, respectively, comparing to the change caused by trifluoperazine in PC3 cell. Rank, p value, enrichment score, and similarity mean are described in Supplementary Table 4. Compound name and cell line indicate the name of compound used for treatment with specific cell lines in CMAP database. Number of arrays (N) is the number of analyzed arrays obtained from all instances of the corresponding compound/drug.

## Supplementary Data Legends

**Supplementary Data 1.** Summary of signature probes corresponding to up-regulated genes (page 1) and down-regulated genes (page 2) in *Fmr1* knockout hippocampal neurons.

**Supplementary Data 2.** Summary of signature probes corresponding to up-regulated genes (page 1) and down-regulated genes (page 2) in PC3 cell treated with trifluoperazine.

**Supplementary Data 3.** List of differentially expressed genes (DEGs) between wild type and Fmr1 knockout neurons. WT: samples from wild type hippocampal neuron. FMR: samples from *Fmr1* knockout (KO) hippocampal neuron. DEG list includes transcripts that are significantly up- or down-regulated in *Fmr1* KO samples (P <0.05). Transcripts that show FPKM (Fragments Per Kilobase of transcript per Million mapped reads) less than 1 in both WT and FMR samples are excluded.

## REFERENCES

1. Santoro, M.R., Bray, S.M. & Warren, S.T. Molecular mechanisms of fragile X syndrome: a twenty-year perspective. Annu Rev Pathol 7, 219–245 (2012).

2. Bagni, C., Tassone, F., Neri, G. & Hagerman, R. Fragile X syndrome: causes, diagnosis, mechanisms, and therapeutics. J Clin Invest 122, 4314–4322 (2012).

3. Sethna, F., Moon, C. & Wang, H. From FMRP function to potential therapies for fragile X syndrome. Neurochem Res 39, 1016–1031 (2014).

4. Tasic, B., et al. Adult mouse cortical cell taxonomy revealed by single cell transcriptomics. Nat Neurosci 19, 335–346 (2016).

5. Voineagu, I., et al. Transcriptomic analysis of autistic brain reveals convergent molecular pathology. Nature 474, 380–384 (2011).

6. Maycox, P.R., et al. Analysis of gene expression in two large schizophrenia cohorts identifies multiple changes associated with nerve terminal function. Mol Psychiatry 14, 1083–1094 (2009).

7. Gandal, M.J., et al. Shared molecular neuropathology across major psychiatric disorders parallels polygenic overlap. Science 359, 693–697 (2018).

8. So, H.C., et al. Analysis of genome-wide association data highlights candidates for drug repositioning in psychiatry. Nat Neurosci 20, 1342–1349 (2017).

9. Lamb, J., et al. The Connectivity Map: using gene-expression signatures to connect small molecules, genes, and disease. Science 313, 1929–1935 (2006).

10. Trapnell, C., et al. Differential gene and transcript expression analysis of RNA-seq experiments with TopHat and Cufflinks. Nat Protoc 7, 562–578 (2012).

11. Huang da, W., Sherman, B.T. & Lempicki, R.A. Systematic and integrative analysis of large gene lists using DAVID bioinformatics resources. Nat Protoc 4, 44–57 (2009).

12. Irizarry, R.A., et al. Exploration, normalization, and summaries of high density oligonucleotide array probe level data. Biostatistics 4, 249–264 (2003).

13. Gautier, L., Cope, L., Bolstad, B.M. & Irizarry, R.A. affy--analysis of Affymetrix GeneChip data at the probe level. Bioinformatics 20, 307–315 (2004).

14. Silverman, J.L., Tolu, S.S., Barkan, C.L. & Crawley, J.N. Repetitive self-grooming behavior in the BTBR mouse model of autism is blocked by the mGluR5 antagonist MPEP. Neuropsychopharmacology 35, 976–989 (2010).

15. Bhattacharya, A., et al. Genetic removal of p70 S6 kinase 1 corrects molecular, synaptic, and behavioral phenotypes in fragile X syndrome mice. Neuron 76, 325–337 (2012).

16. Schmidt, E.K., Clavarino, G., Ceppi, M. & Pierre, P. SUnSET, a nonradioactive method to monitor protein synthesis. Nat Methods 6, 275–277 (2009).

17. Zhou, X., Hollern, D., Liao, J., Andrechek, E. & Wang, H. NMDA receptor-mediated excitotoxicity depends on the coactivation of synaptic and extrasynaptic receptors. Cell Death Dis 4, e560 (2013).

18. Wang, X., et al. Activation of the extracellular signal-regulated kinase pathway contributes to the behavioral deficit of fragile x-syndrome. J Neurochem 121, 672–679 (2012).

19. Michalon, A., et al. Chronic pharmacological mGlu5 inhibition corrects fragile X in adult mice. Neuron 74, 49–56 (2012).

20. Osterweil, E.K., Krueger, D.D., Reinhold, K. & Bear, M.F. Hypersensitivity to mGluR5 and ERK1/2 leads to excessive protein synthesis in the hippocampus of a mouse model of fragile X syndrome. J Neurosci 30, 15616–15627 (2010).

21. Sethna, F., et al. Enhanced expression of ADCY1 underlies aberrant neuronal signalling and behaviour in a syndromic autism model. Nature communications 8, 14359 (2017).

22. Schmalzing, G. Metabolism and disposition of trifluoperazine in the rat. II. Kinetics after oral and intravenous administration in acutely and chronically treated animals. Drug Metab Dispos 5, 104–115 (1977).

23. Albert, K., et al. In vivo 19F nuclear magnetic resonance spectroscopy of trifluorinated neuroleptics in the rat. NMR Biomed 3, 120–123 (1990).

24. Marques, L.O., Lima, M.S. & Soares, B.G. Trifluoperazine for schizophrenia. Cochrane Database Syst Rev, CD003545 (2004).

25. Molokie, R.E., et al. Mechanism-driven phase I translational study of trifluoperazine in adults with sickle cell disease. Eur J Pharmacol (2013).

26. Thomas, A., et al. Marble burying reflects a repetitive and perseverative behavior more than novelty-induced anxiety. Psychopharmacology (Berl) 204, 361–373 (2009).

27. Qin, M., Kang, J. & Smith, C.B. Increased rates of cerebral glucose metabolism in a mouse model of fragile X mental retardation. Proc Natl Acad Sci U S A 99, 15758–15763 (2002).

28. Ding, Q., Sethna, F. & Wang, H. Behavioral analysis of male and female Fmr1 knockout mice on C57BL/6 background. Behav Brain Res 271, 72–78 (2014).

29. Musumeci, S.A., et al. Audiogenic seizures susceptibility in transgenic mice with fragile X syndrome. Epilepsia 41, 19–23 (2000).

30. Qin, M., Kang, J., Burlin, T.V., Jiang, C. & Smith, C.B. Postadolescent changes in regional cerebral protein synthesis: an in vivo study in the FMR1 null mouse. J Neurosci 25, 5087–5095 (2005).

31. Wang, H., et al. FMRP acts as a key messenger for dopamine modulation in the forebrain. Neuron 59, 634–647 (2008).

32. Gross, C., et al. Excess phosphoinositide 3-kinase subunit synthesis and activity as a novel therapeutic target in fragile X syndrome. J Neurosci 30, 10624–10638 (2010).

33. Gross, C. & Bassell, G.J. Excess protein synthesis in FXS patient lymphoblastoid cells can be rescued with a p110beta-selective inhibitor. Molecular medicine 18, 336–345 (2012).

34. Sharma, A., et al. Dysregulation of mTOR signaling in fragile X syndrome. J Neurosci 30, 694–702 (2010).

35. Kumari, D., et al. Identification of fragile X syndrome specific molecular markers in human fibroblasts: a useful model to test the efficacy of therapeutic drugs. Hum Mutat 35, 1485–1494 (2014).

36. Lehman, J.A., Calvo, V. & Gomez-Cambronero, J. Mechanism of ribosomal p70S6 kinase activation by granulocyte macrophage colony-stimulating factor in neutrophils: cooperation of a MEK-related, THR421/SER424 kinase and a rapamycin-sensitive, m-TOR-related THR389 kinase. J Biol Chem 278, 28130–28138 (2003).

37. Zhou, X., Lin, D.S., Zheng, F., Sutton, M.A. & Wang, H. Intracellular calcium and calmodulin link brain-derived neurotrophic factor to p70S6 kinase phosphorylation and dendritic protein synthesis. J Neurosci Res 88, 1420–1432 (2010).

38. Wang, B., Luo, Y., Zhou, X. & Li, R. Trifluoperazine induces apoptosis through the upregulation of Bax/Bcl2 and downregulated phosphorylation of AKT in mesangial cells and improves renal function in lupus nephritis mice. Int J Mol Med 41, 3278–3286 (2018).

39. Gross, C., et al. Increased Expression of the PI3K Enhancer PIKE Mediates Deficits in Synaptic Plasticity and Behavior in Fragile X Syndrome. Cell Rep 11, 727–736 (2015).

40. Osterweil, E.K., et al. Lovastatin corrects excess protein synthesis and prevents epileptogenesis in a mouse model of fragile X syndrome. Neuron 77, 243–250 (2013).

41. Gantois, I., et al. Metformin ameliorates core deficits in a mouse model of fragile X syndrome. Nat Med 23, 674–677 (2017).

42. Dy, A.B.C., et al. Metformin as targeted treatment in fragile X syndrome. Clin Genet 93, 216–222 (2018).

43. Caku, A., Pellerin, D., Bouvier, P., Riou, E. & Corbin, F. Effect of lovastatin on behavior in children and adults with fragile X syndrome: an open-label study. Am J Med Genet A 164A, 2834–2842 (2014).

44. Posey, D.J., Stigler, K.A., Erickson, C.A. & McDougle, C.J. Antipsychotics in the treatment of autism. J Clin Invest 118, 6–14 (2008).

45. Erickson, C.A., et al. A prospective open-label study of aripiprazole in fragile X syndrome. Psychopharmacology (Berl) 216, 85–90 (2011).

46. Berry-Kravis, E. & Potanos, K. Psychopharmacology in fragile X syndrome--present and future. Ment Retard Dev Disabil Res Rev 10, 42–48 (2004).

47. Reagan-Shaw, S., Nihal, M. & Ahmad, N. Dose translation from animal to human studies revisited. FASEB journal: official publication of the Federation of American Societies for Experimental Biology 22, 659–661 (2008).

48. Gross, C., et al. Selective role of the catalytic PI3K subunit p110beta in impaired higher order cognition in fragile X syndrome. Cell Rep 11, 681–688 (2015).

49. Gross, C., et al. Isoform-selective phosphoinositide 3-kinase inhibition ameliorates a broad range of fragile X syndrome-associated deficits in a mouse model. Neuropsychopharmacology 44, 324–333 (2019).

50. Davenport, M.H., Schaefer, T.L., Friedmann, K.J., Fitzpatrick, S.E. & Erickson, C.A. Pharmacotherapy for Fragile X Syndrome: Progress to Date. Drugs 76, 431–445 (2016).

51. Ramanathan, S., Jin, F., Sharma, S. & Kearney, B.P. Clinical Pharmacokinetic and Pharmacodynamic Profile of Idelalisib. Clinical pharmacokinetics (2015).

