## Supplementary Data 1 for "Computational analysis of transcriptome signature repurposes low dose trifluoperazine for the treatment of fragile X syndrome in mouse model"

**probes for up-regulated genes**

| Probe Set ID | Gene Symbol |
|---|---|
| 200951_s_at | CCND2 |
| 200952_s_at | CCND2 |
| 200953_s_at | CCND2 |
| 201028_s_at | CD99 |
| 201029_s_at | CD99 |
| 201291_s_at | TOP2A |
| 201292_at | TOP2A |
| 201430_s_at | DPYSL3 |
| 201431_s_at | DPYSL3 |
| 201559_s_at | CLIC4 |
| 201560_at | CLIC4 |
| 201710_at | MYBL2 |
| 201755_at | MCM5 |
| 201954_at | ARPC1B |
| 201983_s_at | EGFR |
| 201984_s_at | EGFR |
| 202094_at | BIRC5 |
| 202095_s_at | BIRC5 |
| 202111_at | SLC4A2 |
| 202240_at | PLK1 |
| 202468_s_at | CTNNAL1 |
| 202589_at | TYMS |
| 202711_at | EFNB1 |
| 202718_at | IGFBP2 |
| 202781_s_at | INPP5K |
| 202782_s_at | INPP5K |
| 202870_s_at | CDC20 |
| 203131_at | PDGFRA |
| 203184_at | FBN2 |
| 203369_x_at | PDLIM7 |
| 203370_s_at | PDLIM7 |
| 203477_at | COL15A1 |
| 203504_s_at | ABCA1 |
| 203505_at | ABCA1 |
| 203525_s_at | APC |
| 203526_s_at | APC |
| 203527_s_at | APC |
| 203677_s_at | TARBP2 |
| 203702_s_at | TTLL4 |
| 203703_s_at | TTLL4 |
| 203792_x_at | PCGF2 |
| 203793_x_at | PCGF2 |
| 203819_s_at | IGF2BP3 |
| 203820_s_at | IGF2BP3 |
| 203821_at | HBEGF |
| 203967_at | CDC6 |
| 203968_s_at | CDC6 |
| 204033_at | TRIP13 |
| 204069_at | MEIS1 |
| 204082_at | PBX3 |

**probes for down-regulated genes**

| Probe Set ID | Gene Symbol |
|---|---|
| 1729_at | TRADD |
| 200606_at | DSP |
| 200696_s_at | GSN |
| 200872_at | S100A10 |
| 200908_s_at | RPLP2 |
| 200909_s_at | RPLP2 |
| 200916_at | TAGLN2 |
| 201008_s_at | TXNIP |
| 201009_s_at | TXNIP |
| 201010_s_at | TXNIP |
| 201060_x_at | STOM |
| 201061_s_at | STOM |
| 201062_at | STOM |
| 201079_at | SYNGR2 |
| 201141_at | GPNMB |
| 201215_at | PLS3 |
| 201231_s_at | ENO1 |
| 201261_x_at | BGN |
| 201262_s_at | BGN |
| 201274_at | PSMA5 |
| 201301_s_at | ANXA4 |
| 201302_at | ANXA4 |
| 201315_x_at | IFITM2 |
| 201348_at | GPX3 |
| 201400_at | PSMB3 |
| 201601_x_at | IFITM1 /// IFITM2 |
| 201605_x_at | CNN2 |
| 201779_s_at | RNF13 |
| 201780_s_at | RNF13 |
| 201850_at | CAPG |
| 201854_s_at | ATMIN |
| 201855_s_at | ATMIN |
| 201891_s_at | B2M |
| 201893_x_at | DCN |
| 201944_at | HEXB |
| 202196_s_at | DKK3 |
| 202263_at | CYB5R1 |
| 202267_at | LAMC2 |
| 202283_at | SERPINF1 |
| 202340_x_at | NR4A1 |
| 202411_at | IFI27 |
| 202518_at | BCL7B |
| 202555_s_at | MYLK |
| 202605_at | GUSB |
| 202659_at | PSMB10 |
| 202699_s_at | TMEM63A |
| 202700_s_at | TMEM63A |
| 202735_at | EBP |
| 202783_at | NNT |
| 202784_s_at | NNT |

| | | | |
|---|---|---|---|
| 204146_at | RAD51AP1 | 202834_at | AGT |
| 204244_s_at | DBF4 | 202862_at | FAH |
| 204315_s_at | GTSE1 | 202877_s_at | CD93 |
| 204317_at | GTSE1 /// TRMU | 202878_s_at | CD93 |
| 204318_s_at | GTSE1 | 202910_s_at | ADGRE5 |
| 204431_at | TLE2 | 202912_at | ADM |
| 204437_s_at | FOLR1 | 202949_s_at | FHL2 |
| 204484_at | PIK3C2B | 202953_at | C1QB |
| 204603_at | EXO1 | 203009_at | BCAM |
| 204709_s_at | KIF23 | 203028_s_at | CYBA |
| 204775_at | CHAF1B | 203065_s_at | CAV1 |
| 204854_at | P3H3 | 203119_at | CCDC86 |
| 204948_s_at | FST | 203217_s_at | ST3GAL5 |
| 205000_at | DDX3Y | 203313_s_at | TGIF1 |
| 205001_s_at | DDX3Y | 203423_at | RBP1 |
| 205029_s_at | FABP7 | 203430_at | HEBP2 |
| 205030_at | FABP7 | 203436_at | RPP30 |
| 205288_at | CDC14A | 203442_x_at | EML3 |
| 205593_s_at | PDE9A | 203443_at | EML3 |
| 205666_at | FMO1 | 203501_at | CPQ |
| 206070_s_at | EPHA3 | 203507_at | CD68 |
| 206071_s_at | EPHA3 | 203522_at | CCS |
| 206080_at | PLCH2 | 203554_x_at | PTTG1 |
| 206092_x_at | RTEL1 /// RTEL1-TNFRSF6B | 203557_s_at | PCBD1 |
| 206107_at | RGS11 | 203618_at | FAIM2 |
| 206148_at | IL3RA | 203619_s_at | FAIM2 |
| 206322_at | SYN3 | 203636_at | MID1 |
| 206333_at | MSI1 | 203637_s_at | MID1 |
| 206364_at | KIF14 | 203671_at | TPMT |
| 206700_s_at | KDM5D | 203672_x_at | TPMT |
| 206709_x_at | GPT | 203697_at | FRZB |
| 206717_at | MYH8 | 203698_s_at | FRZB |
| 206817_x_at | CELF3 | 203704_s_at | RREB1 |
| 206822_s_at | L3MBTL1 | 203740_at | MPHOSPH6 |
| 206823_at | L3MBTL1 | 203762_s_at | DYNC2LI1 |
| 206825_at | OXTR | 203763_at | DYNC2LI1 |
| 206832_s_at | SEMA3F | 203786_s_at | TPD52L1 |
| 206864_s_at | HRK | 203876_s_at | MMP11 |
| 206865_at | HRK | 203878_s_at | MMP11 |
| 206944_at | HTR6 | 203904_x_at | CD82 |
| 206960_at | LPAR4 | 203910_at | ARHGAP29 |
| 206970_at | CNTN2 | 204014_at | DUSP4 |
| 207030_s_at | CSRP2 | 204015_s_at | DUSP4 |
| 207085_x_at | CSF2RA | 204041_at | MAOB |
| 207123_s_at | MATN4 | 204073_s_at | MYRF |
| 207147_at | DLX2 | 204115_at | GNG11 |
| 207230_at | CDON | 204149_s_at | GSTM4 |
| 207242_s_at | GRIK1 | 204167_at | BTD |
| 207309_at | NOS1 | 204223_at | PRELP |
| 207310_s_at | NOS1 | 204316_at | RGS10 |
| 207322_at | ITSN1 | 204319_s_at | RGS10 |
| 207331_at | CENPF | 204360_s_at | NAGLU |

| | | | |
|---|---|---|---|
| 207345_at | FST | 204378_at | BCAS1 |
| 207480_s_at | MEIS2 | 204428_s_at | LCAT |
| 207528_s_at | SLC7A11 | 204472_at | GEM |
| 207598_x_at | XRCC2 | 204621_s_at | NR4A2 |
| 207694_at | POU3F4 | 204622_x_at | NR4A2 |
| 207828_s_at | CENPF | 204624_at | ATP7B |
| 208050_s_at | CASP2 | 204629_at | PARVB |
| 208056_s_at | CBFA2T3 | 204747_at | IFIT3 |
| 208067_x_at | UTY | 204748_at | PTGS2 |
| 208252_s_at | CHST3 | 204834_at | FGL2 |
| 208258_s_at | GAS2L1 | 204875_s_at | GMDS |
| 208296_x_at | TNFAIP8 | 204916_at | RAMP1 |
| 208345_s_at | POU3F1 | 205007_s_at | CIB2 |
| 208795_s_at | MCM7 | 205008_s_at | CIB2 |
| 208808_s_at | HMGB2 | 205013_s_at | ADORA2A /// SPECC1L |
| 209172_s_at | CENPF | 205084_at | BCAP29 |
| 209234_at | KIF1B | 205090_s_at | NAGPA |
| 209297_at | ITSN1 | 205113_at | NEFM |
| 209298_s_at | ITSN1 | 205190_at | PLS1 |
| 209370_s_at | SH3BP2 | 205200_at | CLEC3B /// EXOSC7 |
| 209371_s_at | SH3BP2 | 205205_at | RELB |
| 209398_at | HIST1H1C | 205229_s_at | COCH |
| 209561_at | THBS3 | 205238_at | TRMT2B |
| 209642_at | BUB1 | 205326_at | RAMP3 |
| 209681_at | SLC19A2 | 205384_at | FXYD1 |
| 209729_at | GAS2L1 | 205404_at | HSD11B1 |
| 209730_at | SEMA3F | 205454_at | HPCA |
| 209756_s_at | MYCN | 205548_s_at | BTG3 |
| 209757_s_at | MYCN | 205549_at | PCP4 |
| 209780_at | PHTF2 | 205586_x_at | VGF |
| 209784_s_at | JAG2 | 205590_at | RASGRP1 |
| 209811_at | CASP2 | 205641_s_at | TRADD |
| 209812_x_at | CASP2 | 205659_at | HDAC9 |
| 209834_at | CHST3 | 205679_x_at | ACAN |
| 209921_at | SLC7A11 | 205755_at | ITIH3 |
| 209960_at | HGF | 205768_s_at | SLC27A2 |
| 209961_s_at | HGF | 205769_at | SLC27A2 |
| 210051_at | RAPGEF3 | 205779_at | RAMP2 |
| 210260_s_at | TNFAIP8 | 205818_at | BRINP1 |
| 210306_at | L3MBTL1 | 205825_at | PCSK1 |
| 210322_x_at | UTY | 205856_at | SLC14A1 |
| 210334_x_at | BIRC5 | 205857_at | SLC18A2 |
| 210340_s_at | CSF2RA | 205874_at | ITPKA |
| 210341_at | MYT1 | 205879_x_at | RET |
| 210440_s_at | CDC14A | 205911_at | PTH1R |
| 210441_at | CDC14A | 205934_at | PLCL1 |
| 210475_at | POU3F1 | 205957_at | PLXNB3 |
| 210495_x_at | FN1 | 205984_at | CRHBP |
| 210614_at | TTPA | 206089_at | NELL1 |
| 210713_at | ITSN1 | 206106_at | MAPK12 |
| 210742_at | CDC14A | 206115_at | EGR3 |
| 210743_s_at | CDC14A | 206231_at | KCNN1 |

| | | | |
|---|---|---|---|
| 210755_at | HGF | 206271_at | TLR3 |
| 210913_at | CDH20 | 206277_at | P2RY2 |
| 210983_s_at | MCM7 | 206280_at | CDH18 |
| 210984_x_at | EGFR | 206281_at | ADCYAP1 |
| 210997_at | HGF | 206291_at | NTS |
| 210998_s_at | HGF | 206373_at | ZIC1 |
| 211011_at | COL19A1 | 206382_s_at | BDNF |
| 211040_x_at | GTSE1 | 206384_at | CACNG3 |
| 211089_s_at | NEK3 | 206404_at | FGF9 |
| 211126_s_at | CSRP2 | 206528_at | TRPC6 |
| 211140_s_at | CASP2 | 206560_s_at | MIA |
| 211149_at | UTY | 206577_at | VIP |
| 211164_at | EPHA3 | 206617_s_at | RENBP |
| 211250_s_at | SH3BP2 | 206721_at | CCDC181 |
| 211286_x_at | CSF2RA | 206765_at | KCNJ2 |
| 211287_x_at | CSF2RA | 206803_at | PDYN |
| 211377_x_at | MYCN | 206880_at | P2RX6 |
| 211526_s_at | RTEL1 /// RTEL1-TNFRSF6B | 206941_x_at | SEMA3E |
| 211533_at | PDGFRA | 207014_at | GABRA2 |
| 211550_at | EGFR | 207035_at | SLC30A3 |
| 211551_at | EGFR | 207055_at | GPR37L1 |
| 211607_x_at | EGFR | 207144_s_at | CITED1 |
| 211675_s_at | MDFIC | 207334_s_at | TGFBR2 |
| 211719_x_at | FN1 | 207388_s_at | PTGES |
| 211872_s_at | RGS11 | 207401_at | PROX1 |
| 211943_x_at | TPT1 | 207455_at | P2RY1 |
| 212012_at | PXDN | 207517_at | LAMC2 |
| 212013_at | PXDN | 207527_at | KCNJ9 |
| 212020_s_at | MKI67 | 207547_s_at | FAM107A |
| 212021_s_at | MKI67 | 207565_s_at | MR1 |
| 212022_s_at | MKI67 | 207566_at | MR1 |
| 212023_s_at | MKI67 | 207589_at | ADRA1B |
| 212284_x_at | TPT1 | 207692_s_at | ACAN |
| 212443_at | NBEAL2 | 207717_s_at | PKP2 |
| 212464_s_at | FN1 | 207767_s_at | EGR4 |
| 212536_at | ATP11B | 207768_at | EGR4 |
| 212556_at | SCRIB | 208123_at | KCNB2 |
| 212619_at | NEMP1 | 208172_s_at | KCNB2 |
| 212621_at | NEMP1 | 208183_at | TACR3 |
| 212841_s_at | PPFIBP2 | 208338_at | P2RX3 |
| 212869_x_at | TPT1 | 208365_s_at | GRK4 |
| 212935_at | MCF2L | 208454_s_at | CPQ |
| 212949_at | NCAPH | 208457_at | GABRD |
| 212962_at | SYDE1 | 208465_at | GRM2 |
| 213116_at | NEK3 | 208550_x_at | KCNG2 |
| 213551_x_at | PCGF2 | 208593_x_at | CRHR1 /// MGC57346 |
| 213707_s_at | DLX5 | 208937_s_at | ID1 |
| 213725_x_at | XYLT1 | 208944_at | TGFBR2 |
| 213829_x_at | RTEL1 /// RTEL1-TNFRSF6B | 208949_s_at | LGALS3 |
| 213837_at | L3MBTL1 | 209034_at | PNRC1 |
| 213920_at | CUX2 | 209074_s_at | FAM107A |
| 213996_at | YPEL1 | 209122_at | PLIN2 |

| | |
|---|---|
| 214121_x_at | PDLIM7 |
| 214122_at | PDLIM7 |
| 214220_s_at | ALMS1 |
| 214221_at | ALMS1 |
| 214239_x_at | PCGF2 |
| 214266_s_at | PDLIM7 |
| 214327_x_at | TPT1 |
| 214455_at | HIST1H2BC |
| 214611_at | GRIK1 |
| 214701_s_at | FN1 |
| 214702_at | FN1 |
| 214707_x_at | ALMS1 |
| 215027_at | RAPGEF3 |
| 215045_at | CELF3 |
| 215218_s_at | WDR62 |
| 215286_s_at | PHTF2 |
| 215305_at | PDGFRA |
| 215310_at | APC |
| 215323_at | LUZP2 |
| 215508_at | BUB1 |
| 215509_s_at | BUB1 |
| 215685_s_at | DLX2 |
| 215717_s_at | FBN2 |
| 215822_x_at | MYT1 |
| 215942_s_at | GTSE1 |
| 216066_at | ABCA1 |
| 216076_at | L3MBTL1 |
| 216077_s_at | L3MBTL1 |
| 216192_at | FABP7 |
| 216237_s_at | MCM5 |
| 216271_x_at | SYDE1 |
| 216272_x_at | SYDE1 |
| 216275_at | BUB1 |
| 216277_at | BUB1 |
| 216442_x_at | FN1 |
| 216493_s_at | IGF2BP3 |
| 216520_s_at | TPT1 |
| 216933_x_at | APC |
| 217066_s_at | DMPK |
| 217097_s_at | PHTF2 |
| 217183_at | SPC24 |
| 217253_at | SH3BP2 |
| 217257_at | SH3BP2 |
| 217599_s_at | MDFIC |
| 217678_at | SLC7A11 |
| 217684_at | TYMS |
| 217733_s_at | TMSB10 |
| 217740_x_at | RPL7A |
| 218148_at | CENPT |
| 218411_s_at | MBIP |
| 218518_at | FAM13B |
| 218585_s_at | DTL |
| 209156_s_at | COL6A2 |
| 209275_s_at | CLN3 |
| 209283_at | CRYAB |
| 209335_at | DCN |
| 209353_s_at | FAM163A |
| 209362_at | MED21 |
| 209363_s_at | MED21 |
| 209536_s_at | EHD4 |
| 209543_s_at | CD34 |
| 209551_at | YIPF4 |
| 209651_at | TGFB1I1 |
| 209676_at | TFPI |
| 209686_at | S100B |
| 209790_s_at | CASP6 |
| 209833_at | CRADD |
| 210042_s_at | CTSZ |
| 210090_at | ARC |
| 210223_s_at | MR1 |
| 210224_at | MR1 |
| 210226_at | NR4A1 |
| 210258_at | RGS13 |
| 210263_at | KCNF1 |
| 210367_s_at | PTGES |
| 210372_s_at | TPD52L1 |
| 210381_s_at | CCKBR |
| 210528_at | MR1 |
| 210600_s_at | GRK4 |
| 210664_s_at | TFPI |
| 210665_at | TFPI |
| 210859_x_at | CLN3 |
| 210912_x_at | GSTM4 |
| 210978_s_at | TAGLN2 |
| 211020_at | GCNT2 |
| 211116_at | SLC9A2 |
| 211143_x_at | NR4A1 |
| 211147_s_at | P2RX6 |
| 211167_s_at | GCK |
| 211421_s_at | RET |
| 211464_x_at | CASP6 |
| 211518_s_at | BMP4 |
| 211538_s_at | HSPA2 |
| 211703_s_at | TM2D1 |
| 211813_x_at | DCN |
| 211896_s_at | DCN |
| 211897_s_at | CRHR1 /// MGC57346 |
| 211986_at | AHNAK |
| 212091_s_at | COL6A1 |
| 212097_at | CAV1 |
| 212203_x_at | IFITM3 |
| 212501_at | CEBPB |
| 212551_at | CAP2 |
| 212554_at | CAP2 |

| | | | |
|---|---|---|---|
| 218586_at | MRGBP | 212562_s_at | CTSZ |
| 218654_s_at | MRPS33 | 212624_s_at | CHN1 |
| 218678_at | NES | 212627_s_at | EXOSC7 |
| 218726_at | HJURP | 212647_at | RRAS |
| 218815_s_at | TMEM51 | 212805_at | PRUNE2 |
| 218821_at | NPEPL1 /// STX16-NPEPL1 | 212806_at | PRUNE2 |
| 218822_s_at | NPEPL1 /// STX16-NPEPL1 | 212937_s_at | COL6A1 |
| 218858_at | DEPTOR | 212938_at | COL6A1 |
| 218875_s_at | FBXO5 | 212939_at | COL6A1 |
| 218913_s_at | GMIP | 212940_at | COL6A1 |
| 219019_at | PIDD1 | 212963_at | TM2D1 |
| 219285_s_at | NIN | 212969_x_at | EML3 |
| 219502_at | NEIL3 | 213134_x_at | BTG3 |
| 219562_at | RAB26 | 213170_at | GPX7 |
| 219621_at | CLSPN | 213182_x_at | CDKN1C |
| 219650_at | ERCC6L | 213228_at | PDE8B |
| 219884_at | LHX6 | 213258_at | TFPI |
| 219888_at | SPAG4 | 213290_at | COL6A2 |
| 220092_s_at | ANTXR1 | 213348_at | CDKN1C |
| 220093_at | ANTXR1 | 213371_at | LDB3 |
| 220124_at | GAN | 213427_at | RPP40 |
| 220429_at | NDST3 | 213428_s_at | COL6A1 |
| 220464_at | MCF2L | 213443_at | TRADD |
| 220476_s_at | FAM212B /// LOC101928718 | 213455_at | FAM114A1 |
| 220492_s_at | OTOF | 213479_at | NPTX2 |
| 220613_s_at | SYTL2 | 213492_at | COL2A1 |
| 220650_s_at | SLC9A5 | 213555_at | RWDD2A |
| 220825_s_at | KIRREL | 213626_at | CBR4 |
| 220997_s_at | DIAPH3 | 213648_at | EXOSC7 |
| 221026_s_at | SCRT1 | 213717_at | LDB3 |
| 221399_at | EDA2R | 213748_at | TRIM66 |
| 221605_s_at | PIPOX | 213779_at | EMID1 |
| 221640_s_at | PIDD1 | 213787_s_at | EBP |
| 221881_s_at | CLIC4 | 213789_at | EBP |
| 222076_at | HBEGF | 213882_at | TM2D1 |
| 31874_at | GAS2L1 | 213883_s_at | TM2D1 |
| 32094_at | CHST3 | 213905_x_at | BGN |
| 32137_at | JAG2 | 213999_at | YIPF4 |
| 34449_at | CASP2 | 214000_s_at | RGS10 |
| 34471_at | MYH8 | 214040_s_at | GSN |
| 35147_at | MCF2L | 214081_at | PLXDC1 |
| 35666_at | SEMA3F | 214091_s_at | GPX3 |
| 35776_at | ITSN1 | 214106_s_at | GMDS |
| 37996_s_at | DMPK | 214116_at | BTD |
| 38037_at | HBEGF | 214117_s_at | BTD |
| 40837_at | TLE2 | 214154_s_at | PKP2 |
| 44702_at | SYDE1 | 214200_s_at | COL6A1 |
| 50965_at | RAB26 | 214218_s_at | XIST |
| 89476_r_at | NPEPL1 | 214247_s_at | DKK3 |
| | | 214378_at | TFPI |
| | | 214613_at | GPR3 |
| | | 214619_at | CRHR1 /// MGC57346- |

| | |
|---|---|
| 214655_at | GPR6 |
| 214703_s_at | MAN2B2 |
| 214729_at | TWISTNB |
| 214733_s_at | YIPF1 |
| 214775_at | N4BP3 |
| 214833_at | TMEM63A |
| 215032_at | RREB1 |
| 215054_at | EPOR /// RGL3 |
| 215104_at | NRIP2 |
| 215425_at | BTG3 |
| 215583_at | TMEM63A |
| 215620_at | RREB1 |
| 215736_at | KCNV1 |
| 215771_x_at | RET |
| 215860_at | SYT12 |
| 215865_at | SYT12 |
| 215880_at | NAGLU |
| 215895_x_at | PLIN2 |
| 216039_at | GABRA2 |
| 216231_s_at | B2M |
| 216248_s_at | NR4A2 |
| 216253_s_at | PARVB |
| 216255_s_at | GRM8 |
| 216256_at | GRM8 |
| 216390_at | LCAT |
| 216434_at | TTC38 |
| 216483_s_at | MYDGF |
| 216648_s_at | RREB1 |
| 216649_at | RREB1 |
| 216768_x_at | TTC38 |
| 216887_s_at | LDB3 |
| 216888_at | LDB3 |
| 216894_x_at | CDKN1C |
| 216904_at | COL6A1 |
| 216992_s_at | GRM8 |
| 217161_x_at | ACAN |
| 217287_s_at | TRPC6 |
| 217294_s_at | ENO1 |
| 217404_s_at | COL2A1 |
| 217411_s_at | RREB1 |
| 217462_at | MYRF |
| 217463_s_at | MYRF |
| 217548_at | ARPIN |
| 217611_at | ERICH1 |
| 217670_at | RPLP2 |
| 217677_at | PLEKHA2 |
| 217700_at | CNPY4 |
| 217794_at | PRR13 |
| 217871_s_at | MIF |
| 217875_s_at | PMEPA1 |
| 217989_at | HSD17B11 |
| 218124_at | RETSAT |

| | |
|---|---|
| 218144_s_at | INF2 |
| 218189_s_at | NANS |
| 218214_at | ATG101 |
| 218232_at | C1QA |
| 218272_at | TTC38 |
| 218322_s_at | ACSL5 |
| 218481_at | EXOSC5 |
| 218500_at | THEM6 |
| 218504_at | FAHD2A |
| 218625_at | NRN1 |
| 218731_s_at | VWA1 |
| 218746_at | TAPBPL |
| 218747_s_at | TAPBPL |
| 218753_at | XKR8 |
| 218773_s_at | MSRB2 |
| 218800_at | SRD5A3 |
| 218831_s_at | FCGRT |
| 218836_at | RPP21 /// TRIM39 /// [illegible] |
| 218996_at | TFPT |
| 219032_x_at | OPN3 |
| 219074_at | TMEM184C |
| 219102_at | RCN3 |
| 219135_s_at | LMF1 |
| 219136_s_at | LMF1 |
| 219140_s_at | RBP4 |
| 219142_at | RASL11B |
| 219263_at | RNF128 |
| 219275_at | PDCD5 |
| 219304_s_at | PDGFD |
| 219416_at | SCARA3 |
| 219429_at | FA2H |
| 219451_at | MSRB2 |
| 219534_x_at | CDKN1C |
| 219561_at | COPZ2 |
| 219569_s_at | SLC35G2 |
| 219655_at | SUGCT |
| 219700_at | PLXDC1 |
| 219771_at | TBC1D8B |
| 219821_s_at | GFOD1 |
| 219896_at | CALY |
| 219914_at | ECEL1 |
| 220016_at | AHNAK |
| 220063_at | GSTCD |
| 220077_at | CCDC134 |
| 220094_s_at | MCUR1 |
| 220230_s_at | CYB5R2 |
| 220294_at | KCNV1 |
| 220345_at | LRRTM4 |
| 220433_at | PRRG3 |
| 220623_s_at | TSGA10 |
| 220631_at | OSGEPL1 |
| 220663_at | IL1RAPL1 |

| | |
|---|---|
| 220732_at | PREX2 |
| 220753_s_at | CRYL1 |
| 220775_s_at | UEVLD |
| 220794_at | GREM2 |
| 220945_x_at | MANSC1 |
| 221041_s_at | SLC17A5 |
| 221049_s_at | POLL |
| 221073_s_at | NOD1 |
| 221126_at | DKK3 |
| 221127_s_at | DKK3 |
| 221142_s_at | PECR |
| 221151_at | PRDM9 |
| 221226_s_at | ASIC4 |
| 221295_at | CIDEA |
| 221363_x_at | GPR25 |
| 221401_at | CACNG5 |
| 221448_s_at | TEX15 |
| 221622_s_at | TMEM126B |
| 221728_x_at | XIST |
| 221739_at | MYDGF |
| 221797_at | OXLD1 |
| 221926_s_at | IL17RC |
| 221967_at | NXPH4 |
| 221991_at | NXPH3 |
| 222049_s_at | RBP4 |
| 222055_at | FAHD2A /// FAHD2B /, |
| 222056_s_at | FAHD2A |
| 222095_s_at | FAM163A |
| 222108_at | AMIGO2 |
| 37022_at | PRELP |
| 37965_at | PARVB |
| 37966_at | PARVB |
| 40093_at | BCAM |
| 46142_at | LMF1 |
| 50374_at | OXLD1 |
| 61734_at | RCN3 |
| 64440_at | IL17RC |
