## Supplementary Data 2 for "Computational analysis of transcriptome signature repurposes low dose trifluoperazine for the treatment of fragile X syndrome in mouse model"

**probes for up-regulated genes**

| Probe_Set_ID | Gene_Symbol |
|---|---|
| 117_at | HSPA6 |
| 200598_s_at | HSP90B1///MIR3652 |
| 200632_s_at | NDRG1 |
| 200825_s_at | HYOU1 |
| 200924_s_at | SLC3A2 |
| 201000_at | AARS |
| 201195_s_at | SLC7A5 |
| 201248_s_at | SREBF2 |
| 201283_s_at | TRAK1 |
| 201471_s_at | SQSTM1 |
| 201531_at | ZFP36 |
| 201625_s_at | INSIG1 |
| 201626_at | INSIG1 |
| 201627_s_at | INSIG1 |
| 201694_s_at | EGR1 |
| 201925_s_at | CD55 |
| 201926_s_at | CD55 |
| 202061_s_at | SEL1L |
| 202284_s_at | CDKN1A |
| 202323_s_at | ACBD3 |
| 202341_s_at | TRIM2 |
| 202402_s_at | CARS |
| 202455_at | HDAC5 |
| 202570_s_at | DLGAP4 |
| 202637_s_at | ICAM1 |
| 202638_s_at | ICAM1 |
| 202655_at | MANF |
| 202665_s_at | WIPF1 |
| 202672_s_at | ATF3 |
| 202679_at | NPC1 |
| 202721_s_at | GFPT1 |
| 202722_s_at | GFPT1 |
| 202769_at | CCNG2 |
| 202828_s_at | MMP14 |
| 202859_x_at | IL8 |
| 202887_s_at | DDIT4 |
| 202889_x_at | MAP7 |
| 203071_at | SEMA3B |
| 203096_s_at | RAPGEF2 |
| 203205_at | KDM4A |
| 203231_s_at | ATXN1 |
| 203252_at | CDK2AP2 |
| 203439_s_at | STC2 |
| 203455_s_at | SAT1 |
| 203665_at | HMOX1 |
| 203752_s_at | JUND |
| 203827_at | WIPI1 |
| 203857_s_at | PDIA5 |
| 203879_at | PIK3CD |
| 204203_at | CEBPG |

**probes for down-regulated genes**

| Probe_Set_ID | Gene_Symbol |
|---|---|
| 200072_s_at | HNRNPM |
| 200659_s_at | PHB |
| 200753_x_at | SRSF2 |
| 200769_s_at | MAT2A |
| 200811_at | CIRBP |
| 201035_s_at | HADH |
| 201129_at | SRSF7 |
| 201282_at | OGDH |
| 201297_s_at | MOB1A |
| 201363_s_at | IVNS1ABP |
| 201370_s_at | CUL3 |
| 201394_s_at | RBM5 |
| 201458_s_at | BUB3 |
| 201478_s_at | DKC1///SNORA56 |
| 201555_at | MCM3 |
| 201614_s_at | RUVBL1 |
| 201677_at | C3orf37 |
| 201742_x_at | SRSF1 |
| 201755_at | MCM5 |
| 201767_s_at | ELAC2 |
| 201869_s_at | TBL1X |
| 201939_at | PLK2 |
| 201969_at | NASP |
| 202032_s_at | MAN2A2 |
| 202094_at | BIRC5 |
| 202107_s_at | MCM2 |
| 202152_x_at | USF2 |
| 202154_x_at | TUBB3 |
| 202320_at | GTF3C1 |
| 202330_s_at | UNG |
| 202353_s_at | PSMD12 |
| 202384_s_at | TCOF1 |
| 202385_s_at | TCOF1 |
| 202444_s_at | ERLIN1 |
| 202474_s_at | HCFC1 |
| 202577_s_at | DDX19A |
| 202669_s_at | EFNB2 |
| 202692_s_at | UBTF |
| 202783_at | NNT |
| 202814_s_at | HEXIM1 |
| 202907_s_at | NBN |
| 202911_at | MSH6 |
| 202993_at | ILVBL |
| 203119_at | CCDC86 |
| 203177_x_at | TFAM |
| 203378_at | PCF11 |
| 203418_at | CCNA2 |
| 203426_s_at | IGFBP5 |
| 203432_at | TMPO |
| 203482_at | FAM178A |

| | | | |
|---|---|---|---|
| 204257_at | FADS3 | 203625_x_at | SKP2 |
| 204284_at | PPP1R3C | 203646_at | FDX1 |
| 204285_s_at | PMAIP1 | 203678_at | FAN1 |
| 204296_at | DCTN1 | 203712_at | KIAA0020 |
| 204351_at | S100P | 203774_at | MTR |
| 204475_at | MMP1 | 203775_at | SLC25A13 |
| 204595_s_at | STC1 | 203817_at | GUCY1B3 |
| 204615_x_at | IDI1 | 203820_s_at | IGF2BP3 |
| 204678_s_at | KCNK1 | 203905_at | PARN |
| 204679_at | KCNK1 | 203921_at | CHST2 |
| 204698_at | ISG20 | 203941_at | INTS9 |
| 204786_s_at | IFNAR2 | 203967_at | CDC6 |
| 204821_at | BTN3A3 | 204108_at | NFYA |
| 204948_s_at | FST | 204178_s_at | RBM14 |
| 204952_at | LYPD3 | 204484_at | PIK3C2B |
| 205032_at | ITGA2 | 204492_at | ARHGAP11A |
| 205047_s_at | ASNS | 204510_at | CDC7 |
| 205076_s_at | MTMR11 | 204521_at | FAM216A |
| 205141_at | ANG | 204603_at | EXO1 |
| 205380_at | PDZK1 | 204666_s_at | SIKE1 |
| 205463_s_at | PDGFA | 204742_s_at | PDS5B |
| 205510_s_at | FLJ10038 | 204767_s_at | FEN1 |
| 205570_at | PIP4K2A | 204768_s_at | FEN1 |
| 205797_s_at | TCP11L1 | 204775_at | CHAF1B |
| 206019_at | RBM19 | 204795_at | PRR3 |
| 206048_at | OVOL2 | 204799_at | ZBED4 |
| 206085_s_at | CTH | 204824_at | ENDOG |
| 206155_at | ABCC2 | 204897_at | PTGER4 |
| 206170_at | ADRB2 | 205085_at | ORC1 |
| 206248_at | PRKCE | 205264_at | CD3EAP |
| 206340_at | NR1H4 | 205293_x_at | BAIAP2 |
| 206416_at | ZNF205 | 205345_at | BARD1 |
| 206474_at | CDK17 | 205698_s_at | MAP2K6 |
| 206509_at | PIP | 205739_x_at | ZNF107 |
| 206569_at | IL24 | 205895_s_at | NOLC1 |
| 206683_at | ZNF165 | 205909_at | POLE2 |
| 206864_s_at | HRK | 205963_s_at | DNAJA3 |
| 206969_at | KRT34///LOC100653049 | 206095_s_at | SRSF10 |
| 207137_at | TONSL | 206432_at | HAS2 |
| 207169_x_at | DDR1///MIR4640 | 206468_s_at | METTL13 |
| 207230_at | CDON | 206920_s_at | GLE1 |
| 207292_s_at | MAPK7 | 207304_at | ZNF45 |
| 207345_at | FST | 207490_at | TUBA4B |
| 207526_s_at | IL1RL1 | 207598_x_at | XRCC2 |
| 207604_s_at | SLC4A7 | 208535_x_at | COL13A1 |
| 207709_at | PRKAA2 | 208541_x_at | TFAM |
| 207849_at | IL2 | 208721_s_at | ANAPC5 |
| 207850_at | CXCL3 | 208913_at | GGA2 |
| 207904_s_at | LNPEP | 208931_s_at | ILF3 |
| 208381_s_at | SGPL1 | 208954_s_at | LARP4B |
| 208434_at | MECOM | 209068_at | HNRPDL |
| 208499_s_at | DNAJC3 | 209085_x_at | RFC1 |

| | | | |
|---|---|---|---|
| 208693_s_at | GARS | 209196_at | WDR46 |
| 208786_s_at | MAP1LC3B | 209336_at | PWP2 |
| 208869_s_at | GABARAPL1 | 209355_s_at | PPAP2B |
| 208881_x_at | IDI1 | 209468_at | LRP5 |
| 208926_at | NEU1 | 209520_s_at | NCBP1 |
| 208960_s_at | KLF6 | 209527_at | EXOSC2 |
| 208961_s_at | KLF6 | 209567_at | RRS1 |
| 208966_x_at | IFI16 | 209602_s_at | GATA3 |
| 209034_at | PNRC1 | 209604_s_at | GATA3 |
| 209102_s_at | HBP1 | 209627_s_at | OSBPL3 |
| 209146_at | MSMO1 | 209629_s_at | NXT2 |
| 209173_at | AGR2 | 209646_x_at | ALDH1B1 |
| 209230_s_at | NUPR1 | 209849_s_at | RAD51C |
| 209344_at | TPM4 | 209859_at | TRIM9 |
| 209357_at | CITED2 | 210008_s_at | MRPS12 |
| 209383_at | DDIT3 | 210078_s_at | KCNAB1 |
| 209386_at | TM4SF1 | 210230_at | |
| 209387_s_at | TM4SF1 | 210338_s_at | HSPA8///SNORD14C///SNORD1[illegible] |
| 209435_s_at | ARHGEF2 | 210347_s_at | BCL11A |
| 209473_at | ENTPD1 | 210469_at | DLG5 |
| 209682_at | CBLB | 210570_x_at | MAPK9 |
| 209732_at | CLEC2B | 211042_x_at | MCAM |
| 209774_x_at | CXCL2 | 211080_s_at | NEK2 |
| 209921_at | SLC7A11 | 211090_s_at | PRPF4B |
| 209925_at | OCLN | 211168_s_at | UPF1 |
| 210041_s_at | PGM3 | 211177_s_at | TXNRD2 |
| 210044_s_at | LYL1 | 211212_s_at | ORC5 |
| 210422_x_at | SLC11A1 | 211450_s_at | MSH6 |
| 210517_s_at | AKAP12 | 211501_s_at | EIF3B |
| 210547_x_at | ICA1 | 211535_s_at | FGFR1 |
| 210592_s_at | SAT1 | 211576_s_at | SLC19A1 |
| 210613_s_at | SYNGR1 | 211713_x_at | KIAA0101 |
| 210689_at | CLDN14 | 211812_s_at | B3GALNT1 |
| 210778_s_at | MIR4800///MXD4 | 211814_s_at | CCNE2 |
| 210827_s_at | ELF3 | 211929_at | HNRNPA3 |
| 210835_s_at | CTBP2 | 211949_s_at | NOLC1 |
| 210942_s_at | ST3GAL6 | 211998_at | H3F3A///H3F3B |
| 210943_s_at | LYST | 212001_at | SUGP2 |
| 210961_s_at | ADRA1D | 212020_s_at | MKI67 |
| 210999_s_at | GRB10 | 212021_s_at | MKI67 |
| 211012_s_at | PML | 212023_s_at | MKI67 |
| 211048_s_at | PDIA4 | 212037_at | PNN |
| 211434_s_at | CCRL2 | 212088_at | PMPCA |
| 211559_s_at | CCNG2 | 212126_at | CBX5 |
| 211620_x_at | LOC100506403///RUNX1 | 212140_at | PDS5A |
| 211936_at | HSPA5 | 212141_at | MCM4 |
| 212236_x_at | JUP///KRT17 | 212147_at | SMG5 |
| 212259_s_at | PBXIP1 | 212168_at | RBM12 |
| 212290_at | SLC7A1 | 212170_at | RBM12 |
| 212345_s_at | CREB3L2 | 212192_at | KCTD12 |
| 212452_x_at | KAT6B | 212226_s_at | PPAP2B |
| 212501_at | CEBPB | 212230_at | PPAP2B |

| | | | |
|---|---|---|---|
| 212531_at | LCN2 | 212242_at | TUBA4A |
| 212717_at | PLEKHM1 | 212267_at | WAPAL |
| 212830_at | MEGF9 | 212299_at | NEK9 |
| 212831_at | MEGF9 | 212303_x_at | KHSRP |
| 212884_x_at | APOE | 212729_at | DLG3 |
| 212902_at | SEC24A | 212751_at | UBE2N |
| 212950_at | GPR116 | 212767_at | MTG1 |
| 213046_at | PABPN1 | 212909_at | LYPD1 |
| 213112_s_at | SQSTM1 | 212949_at | NCAPH |
| 213182_x_at | CDKN1C | 212973_at | RPIA |
| 213242_x_at | KIAA0284 | 213094_at | GPR126 |
| 213258_at | TFPI | 213099_at | ANGEL1 |
| 213261_at | TRANK1 | 213130_at | ZNF473 |
| 213348_at | CDKN1C | 213211_s_at | TAF6L |
| 213418_at | HSPA6 | 213323_s_at | ZC3H7B |
| 213577_at | SQLE | 213333_at | MDH2 |
| 213689_x_at | FAM69A | 213402_at | ZNF787 |
| 213810_s_at | AKIRIN2-AS1 | 213425_at | WNT5A |
| 213836_s_at | WIPI1 | 213605_s_at | |
| 213847_at | PRPH | 213610_s_at | KLHL23///PHOSPHO2-KLHL23 |
| 213988_s_at | SAT1 | 213634_s_at | TRMU |
| 214177_s_at | PBXIP1 | 213647_at | DNA2 |
| 214207_s_at | CARD10 | 213718_at | RBM4 |
| 214212_x_at | FERMT2 | 213785_at | IPO9 |
| 214238_at | | 213826_s_at | H3F3A |
| 214297_at | CSPG4 | 213838_at | NOL7 |
| 214316_x_at | CALR | 213906_at | MYBL1 |
| 214435_x_at | RALA | 213947_s_at | NUP210 |
| 214681_at | GK | 214123_s_at | NOP14-AS1 |
| 215034_s_at | TM4SF1 | 214141_x_at | SRSF7 |
| 215134_at | PI4K2A | 214210_at | SLC25A17 |
| 215447_at | | 214331_at | TSFM |
| 215507_x_at | | 214426_x_at | CHAF1A |
| 215518_at | STXBP5L | 214427_at | NOP2 |
| 215741_x_at | AKAP8L | 214507_s_at | EXOSC2 |
| 215855_s_at | TMF1 | 214537_at | HIST1H1D |
| 216336_x_at | LOC100505584///MT1E | 214649_s_at | MTMR2 |
| 216375_s_at | ETV5 | 214694_at | MPRIP |
| 216450_x_at | HSP90B1 | 214764_at | RRP15 |
| 216867_s_at | PDGFA | 214769_at | CLCN4 |
| 217019_at | | 214831_at | ELK4 |
| 217107_at | | 214876_s_at | TUBGCP5 |
| 217127_at | CTH | 214947_at | FAM105A |
| 217165_x_at | MT1F | 214962_s_at | NUP160 |
| 217168_s_at | HERPUD1 | 215006_at | |
| 217206_at | | 215128_at | |
| 217363_x_at | | 215221_at | |
| 217550_at | ATF6 | 215390_at | |
| 217678_at | SLC7A11 | 215470_at | GTF2H2B |
| 217681_at | WNT7B | 215578_at | |
| 217691_x_at | SLC16A3 | 215629_s_at | DLEU2///DLEU2L |
| 217966_s_at | FAM129A | 215714_s_at | SMARCA4 |

| | | | |
|---|---|---|---|
| 218033_s_at | SNN | 215905_s_at | SNRNP40 |
| 218048_at | COMMD3 | 215908_at | |
| 218088_s_at | RRAGC | 216174_at | HCRP1 |
| 218121_at | HMOX2 | 216211_at | |
| 218145_at | TRIB3 | 216262_s_at | TGIF2 |
| 218237_s_at | SLC38A1 | 216650_at | |
| 218510_x_at | FAM134B | 216958_s_at | IVD |
| 218681_s_at | SDF2L1 | 216969_s_at | KIF22 |
| 218696_at | EIF2AK3 | 217437_s_at | TACC1 |
| 219256_s_at | SH3TC1 | 217791_s_at | ALDH18A1 |
| 219429_at | FA2H | 217884_at | NAT10 |
| 219455_at | C7orf63 | 218114_at | GGA1 |
| 219474_at | C3orf52 | 218119_at | TIMM23 |
| 219477_s_at | MRPS31P3///THSD1///THSD1P | 218156_s_at | TSR1 |
| 219566_at | PLEKHF1 | 218248_at | FAM111A |
| 219600_s_at | TMEM50B | 218308_at | TACC3 |
| 219693_at | AGPAT4 | 218350_s_at | GMNN |
| 219878_s_at | KLF13 | 218414_s_at | NDE1 |
| 219910_at | FICD | 218455_at | NFS1 |
| 220016_at | AHNAK | 218530_at | FHOD1 |
| 220058_at | C17orf39 | 218585_s_at | DTL |
| 220114_s_at | STAB2 | 218630_at | MKS1 |
| 220122_at | MCTP1 | 218663_at | NCAPG |
| 220136_s_at | CRYBA2 | 218691_s_at | PDLIM4 |
| 220559_at | EN1 | 218702_at | FBXO17///SARS2 |
| 220609_at | LOC202181 | 218754_at | NOL9 |
| 220874_at | | 218755_at | KIF20A |
| 220917_s_at | WDR19 | 218787_x_at | CWF19L1 |
| 220987_s_at | AKIP1///NUAK2 | 218884_s_at | GUF1 |
| 221030_s_at | ARHGAP24 | 218888_s_at | NETO2 |
| 221041_s_at | SLC17A5 | 218897_at | TMEM177 |
| 221050_s_at | GTPBP2 | 218948_at | QRSL1 |
| 221577_x_at | GDF15 | 218949_s_at | QRSL1 |
| 221664_s_at | F11R | 218993_at | RNMTL1 |
| 221734_at | PRRC1 | 219000_s_at | DSCC1 |
| 221756_at | PIK3IP1 | 219002_at | FASTKD1 |
| 221778_at | JHDM1D | 219048_at | PIGN |
| 221985_at | KLHL24 | 219083_at | SHQ1 |
| 221986_s_at | KLHL24 | 219149_x_at | DBR1 |
| 222024_s_at | AKAP13 | 219158_s_at | NAA15 |
| 222108_at | AMIGO2 | 219217_at | NARS2 |
| 222121_at | ARHGEF26 | 219224_x_at | ZNF408 |
| 222128_at | NSUN6 | 219328_at | DDX31 |
| 33304_at | ISG20 | 219353_at | NHLRC2 |
| 37028_at | PPP1R15A | 219494_at | FSBP///RAD54B |
| 38241_at | BTN3A3 | 219510_at | POLQ |
| 38521_at | CD22 | 219512_at | DSN1 |
| 39248_at | AQP3 | 219705_at | QSER1 |
| | | 219792_at | AGMAT |
| | | 219936_s_at | GPR87 |
| | | 219990_at | E2F8 |
| | | 220030_at | STYK1 |

| | |
|---|---|
| 220085_at | HELLS |
| 220089_at | L2HGDH |
| 220296_at | |
| 220319_s_at | MYLIP |
| 220353_at | FAM86C1 |
| 220408_x_at | FAM48A |
| 220651_s_at | MCM10 |
| 220865_s_at | PDSS1 |
| 220915_s_at | |
| 221189_s_at | TARS2 |
| 221436_s_at | CDCA3 |
| 221521_s_at | GINS2 |
| 221649_s_at | PPAN///PPAN-P2RY11 |
| 221703_at | BRIP1 |
| 221727_at | SUB1 |
| 221774_x_at | FAM48A |
| 221831_at | LUZP1 |
| 221860_at | HNRNPL |
| 221913_at | SIRT3 |
| 221919_at | HNRNPA1 |
| 222006_at | LETM1 |
| 222036_s_at | MCM4 |
| 222120_at | ZNF764 |
| 222228_s_at | ALKBH4 |
| 222294_s_at | RAB27A |
| 222305_at | HK2 |
| 47105_at | DUS2L |
| 52731_at | AMBRA1 |
