## Supplementary Table 1 for "Computational analysis of transcriptome signature repurposes low dose trifluoperazine for the treatment of fragile X syndrome in mouse model"

| **Rank** | **GO Biology Process** | **Count** | **%** | **P-Value** |
| --- | --- | --- | --- | --- |
| 1 | cell adhesion | 54 | 4.2 | 1.70E-06 |
| 2 | positive regulation of cytosolic calcium ion concentration | 25 | 2 | 1.90E-06 |
| 3 | nervous system development | 41 | 3.2 | 6.10E-05 |
| 4 | negative regulation of neuron apoptotic process | 23 | 1.8 | 7.20E-05 |
| 5 | positive regulation of cell migration | 26 | 2 | 1.50E-04 |
| 6 | cell cycle | 57 | 4.5 | 1.60E-04 |
| 7 | cell division | 39 | 3.1 | 2.20E-04 |
| 8 | cellular response to mechanical stimulus | 14 | 1.1 | 3.20E-04 |
| 9 | mitotic nuclear division | 31 | 2.4 | 3.40E-04 |
| 10 | regulation of G-protein coupled receptor protein signaling pathway | 9 | 0.7 | 4.30E-04 |
| 11 | positive regulation of GTPase activity | 20 | 1.6 | 4.50E-04 |
| 12 | positive chemotaxis | 6 | 0.5 | 9.70E-04 |
| 13 | positive regulation of endothelial cell proliferation | 12 | 0.9 | 1.00E-03 |
| 14 | regulation of ion transmembrane transport | 18 | 1.4 | 1.30E-03 |
| 15 | response to estradiol | 15 | 1.2 | 1.30E-03 |
| 16 | positive regulation of synaptic transmission, GABAergic | 6 | 0.5 | 1.30E-03 |
| 17 | wound healing | 14 | 1.1 | 2.00E-03 |
| 18 | response to mechanical stimulus | 11 | 0.9 | 2.30E-03 |
| 19 | odontogenesis of dentin-containing tooth | 11 | 0.9 | 2.30E-03 |
| 20 | single organismal cell-cell adhesion | 15 | 1.2 | 2.50E-03 |
| 21 | ERK1 and ERK2 cascade | 7 | 0.5 | 2.50E-03 |
| 22 | cell differentiation | 63 | 4.9 | 2.50E-03 |
| 23 | response to oxidative stress | 17 | 1.3 | 2.80E-03 |
| 24 | hepatocyte proliferation | 4 | 0.3 | 3.00E-03 |
| 25 | regulation of axon diameter | 4 | 0.3 | 3.00E-03 |
| 26 | sequestering of actin monomers | 5 | 0.4 | 3.20E-03 |
| 27 | positive regulation of actin filament bundle assembly | 5 | 0.4 | 3.20E-03 |
| 28 | axon guidance | 18 | 1.4 | 3.60E-03 |
| 29 | potassium ion transmembrane transport | 13 | 1 | 3.90E-03 |
| 30 | positive regulation of neuron projection development | 17 | 1.3 | 4.00E-03 |
| 31 | oxidation-reduction process | 55 | 4.3 | 4.20E-03 |
| 32 | positive regulation of vasodilation | 8 | 0.6 | 4.30E-03 |
| 33 | potassium ion transport | 16 | 1.3 | 4.40E-03 |
| 34 | xenophagy | 14 | 1.1 | 4.40E-03 |
| 35 | mitotic chromosome condensation | 5 | 0.4 | 4.40E-03 |
| 36 | positive regulation of cAMP biosynthetic process | 9 | 0.7 | 5.10E-03 |
| 37 | kidney development | 16 | 1.3 | 5.50E-03 |
| 38 | negative regulation of cell migration | 14 | 1.1 | 5.60E-03 |
| 39 | receptor internalization | 8 | 0.6 | 5.80E-03 |
| 40 | response to cold | 8 | 0.6 | 5.80E-03 |
| 41 | cytoskeleton-dependent intracellular transport | 5 | 0.4 | 6.00E-03 |
| 42 | negative regulation of protein kinase activity | 13 | 1 | 6.10E-03 |
| 43 | mitophagy in response to mitochondrial depolarization | 16 | 1.3 | 6.30E-03 |
| 44 | positive regulation of apoptotic process | 31 | 2.4 | 6.50E-03 |
| 45 | positive regulation of protein phosphorylation | 20 | 1.6 | 6.60E-03 |
| 46 | fatty acid biosynthetic process | 11 | 0.9 | 7.40E-03 |
| 47 | neuron projection morphogenesis | 10 | 0.8 | 7.50E-03 |
| 48 | positive regulation of penile erection | 4 | 0.3 | 7.70E-03 |
| 49 | astrocyte cell migration | 4 | 0.3 | 7.70E-03 |
| 50 | response to estrogen | 11 | 0.9 | 8.10E-03 |
| 51 | negative regulation of cell proliferation | 34 | 2.7 | 8.20E-03 |
| 52 | negative regulation of signal transduction | 9 | 0.7 | 8.20E-03 |
| 53 | cellular response to organic substance | 7 | 0.5 | 8.60E-03 |
| 54 | brain development | 22 | 1.7 | 8.70E-03 |
| 55 | negative regulation of MAP kinase activity | 8 | 0.6 | 8.70E-03 |
| 56 | positive regulation of synaptic transmission, glutamatergic | 6 | 0.5 | 9.00E-03 |
| 57 | lung morphogenesis | 6 | 0.5 | 9.00E-03 |
| 58 | chromosome segregation | 12 | 0.9 | 9.90E-03 |
| 59 | response to amphetamine | 7 | 0.5 | 1.00E-02 |
| 60 | extracellular matrix organization | 14 | 1.1 | 1.00E-02 |
| 61 | apoptotic process | 46 | 3.6 | 1.00E-02 |
| 62 | positive regulation of vasculogenesis | 4 | 0.3 | 1.10E-02 |
| 63 | positive regulation of adenylate cyclase activity involved in G-protein coupled receptor signaling pathway | 4 | 0.3 | 1.10E-02 |
| 64 | negative regulation of Wnt signaling pathway | 9 | 0.7 | 1.10E-02 |
| 65 | positive regulation of MAPK cascade | 13 | 1 | 1.20E-02 |
| 66 | positive regulation of osteoblast differentiation | 10 | 0.8 | 1.20E-02 |
| 67 | synaptic vesicle endocytosis | 5 | 0.4 | 1.20E-02 |
| 68 | negative regulation of cell death | 11 | 0.9 | 1.30E-02 |
| 69 | branching morphogenesis of an epithelial tube | 7 | 0.5 | 1.30E-02 |
| 70 | negative regulation of cell cycle | 7 | 0.5 | 1.30E-02 |
| 71 | angiogenesis | 23 | 1.8 | 1.30E-02 |
| 72 | lipid metabolic process | 38 | 3 | 1.40E-02 |
| 73 | positive regulation of synapse assembly | 10 | 0.8 | 1.50E-02 |
| 74 | lipid storage | 6 | 0.5 | 1.50E-02 |
| 75 | skeletal system development | 13 | 1 | 1.50E-02 |
| 76 | mitotic spindle assembly checkpoint | 5 | 0.4 | 1.50E-02 |
| 77 | aging | 18 | 1.4 | 1.50E-02 |
| 78 | cellular response to corticotropin-releasing hormone stimulus | 3 | 0.2 | 1.70E-02 |
| 79 | negative regulation of hydrogen peroxide-mediated programmed cell death | 3 | 0.2 | 1.70E-02 |
| 80 | adherens junction maintenance | 3 | 0.2 | 1.70E-02 |
| 81 | multicellular organism development | 74 | 5.8 | 1.70E-02 |
| 82 | regulation of heart rate by cardiac conduction | 6 | 0.5 | 1.70E-02 |
| 83 | positive regulation of defense response to virus by host | 14 | 1.1 | 1.70E-02 |
| 84 | behavioral response to cocaine | 5 | 0.4 | 1.90E-02 |
| 85 | cell migration | 19 | 1.5 | 1.90E-02 |
| 86 | renal system process | 4 | 0.3 | 2.00E-02 |
| 87 | labyrinthine layer development | 4 | 0.3 | 2.00E-02 |
| 88 | protein localization to plasma membrane | 9 | 0.7 | 2.00E-02 |
| 89 | substrate adhesion-dependent cell spreading | 7 | 0.5 | 2.20E-02 |
| 90 | response to progesterone | 6 | 0.5 | 2.30E-02 |
| 91 | negative regulation of neuron projection development | 9 | 0.7 | 2.40E-02 |
| 92 | calcium ion transport | 15 | 1.2 | 2.40E-02 |
| 93 | embryonic skeletal joint morphogenesis | 4 | 0.3 | 2.50E-02 |
| 94 | actin crosslink formation | 4 | 0.3 | 2.50E-02 |
| 95 | regulation of G2/M transition of mitotic cell cycle | 4 | 0.3 | 2.50E-02 |
| 96 | regulation of cell migration | 10 | 0.8 | 2.60E-02 |
| 97 | negative regulation of GTPase activity | 6 | 0.5 | 2.60E-02 |
| 98 | positive regulation of fibroblast proliferation | 9 | 0.7 | 2.60E-02 |
| 99 | amino acid transport | 7 | 0.5 | 2.70E-02 |
| 100 | regulation of synaptic plasticity | 7 | 0.5 | 2.70E-02 |
| 101 | behavioral fear response | 7 | 0.5 | 2.70E-02 |
| 102 | epithelial cell differentiation | 9 | 0.7 | 3.10E-02 |
| 103 | morphogenesis of embryonic epithelium | 4 | 0.3 | 3.20E-02 |
| 104 | inhibitory postsynaptic potential | 4 | 0.3 | 3.20E-02 |
| 105 | negative regulation of cardiac muscle cell proliferation | 4 | 0.3 | 3.20E-02 |
| 106 | negative regulation of G-protein coupled receptor protein signaling pathway | 4 | 0.3 | 3.20E-02 |
| 107 | embryonic skeletal system development | 7 | 0.5 | 3.30E-02 |
| 108 | positive regulation of cytokinesis | 6 | 0.5 | 3.40E-02 |
| 109 | cellular response to estradiol stimulus | 6 | 0.5 | 3.40E-02 |
| 110 | response to cytokine | 10 | 0.8 | 3.40E-02 |
| 111 | female pregnancy | 10 | 0.8 | 3.40E-02 |
| 112 | positive regulation of angiogenesis | 13 | 1 | 3.60E-02 |
| 113 | positive regulation of vasoconstriction | 7 | 0.5 | 3.60E-02 |
| 114 | ovarian follicle development | 8 | 0.6 | 3.70E-02 |
| 115 | semaphorin-plexin signaling pathway | 6 | 0.5 | 3.80E-02 |
| 116 | positive regulation of neurogenesis | 6 | 0.5 | 3.80E-02 |
| 117 | endothelial cell morphogenesis | 4 | 0.3 | 3.90E-02 |
| 118 | tissue remodeling | 4 | 0.3 | 3.90E-02 |
| 119 | negative regulation of potassium ion transport | 4 | 0.3 | 3.90E-02 |
| 120 | cell activation | 4 | 0.3 | 3.90E-02 |
| 121 | cerebral cortex GABAergic interneuron migration | 3 | 0.2 | 3.90E-02 |
| 122 | intermediate filament bundle assembly | 3 | 0.2 | 3.90E-02 |
| 123 | Toll signaling pathway | 3 | 0.2 | 3.90E-02 |
| 124 | astrocyte activation | 3 | 0.2 | 3.90E-02 |
| 125 | regulation of cell growth | 8 | 0.6 | 4.00E-02 |
| 126 | positive regulation of transcription, DNA-templated | 43 | 3.4 | 4.10E-02 |
| 127 | negative regulation of microtubule depolymerization | 5 | 0.4 | 4.10E-02 |
| 128 | DNA replication initiation | 5 | 0.4 | 4.10E-02 |
| 129 | smoothened signaling pathway | 9 | 0.7 | 4.20E-02 |
| 130 | cellular response to BMP stimulus | 6 | 0.5 | 4.20E-02 |
| 131 | cellular response to DNA damage stimulus | 33 | 2.6 | 4.20E-02 |
| 132 | sensory perception of sound | 14 | 1.1 | 4.20E-02 |
| 133 | neuronal stem cell population maintenance | 5 | 0.4 | 4.70E-02 |
| 134 | negative regulation of peptidyl-serine phosphorylation | 5 | 0.4 | 4.70E-02 |
| 135 | regulation of double-strand break repair via homologous recombination | 4 | 0.3 | 4.70E-02 |
| 136 | osteoblast differentiation | 12 | 0.9 | 4.80E-02 |
| 137 | ion transport | 43 | 3.4 | 4.90E-02 |
| 138 | neuron differentiation | 13 | 1 | 4.90E-02 |
