## Supplementary Table 2 for "Computational analysis of transcriptome signature repurposes low dose trifluoperazine for the treatment of fragile X syndrome in mouse model"

| **Cell adhesion** | | |
| --- | --- | --- |
| **Symbol** | **Gene name** | **Log2 (fold change)** |
| Atp1b2 | ATPase, Na+/K+ transporting, beta 2 polypeptide(Atp1b2) | -0.209579 |
| Cd24a | CD24a antigen(Cd24a) | 0.208692 |
| Cd34 | CD34 antigen(Cd34) | -0.85533 |
| Cd47 | CD47 antigen (Rh-related antigen, integrin-associated signal transducer)(Cd47) | -0.193603 |
| Cd63 | CD63 antigen(Cd63) | -0.22166 |
| Cd9 | CD9 antigen(Cd9) | -0.22731 |
| Cd93 | CD93 antigen(Cd93) | -1.0693 |
| Cd99l2 | CD99 antigen-like 2(Cd99l2) | 0.336254 |
| Epha8 | Eph receptor A8(Epha8) | 0.320632 |
| Fat4 | FAT atypical cadherin 4(Fat4) | 0.230342 |
| Frem2 | Fras1 related extracellular matrix protein 2(Frem2) | 0.265503 |
| Radil | Ras association and DIL domains(Radil) | 0.270472 |
| Adgre5 | adhesion G protein-coupled receptor E5(Adgre5) | -0.580957 |
| Amigo2 | adhesion molecule with Ig like domain 2(Amigo2) | -0.41596 |
| Afdn | afadin, adherens junction formation factor(Afdn) | 0.202055 |
| Acan | aggrecan(Acan) | -0.57369 |
| Bcam | basal cell adhesion molecule(Bcam) | -0.364333 |
| Bcan | brevican(Bcan) | 0.198962 |
| Cdh12 | cadherin 12(Cdh12) | -0.261253 |
| Cdh20 | cadherin 20(Cdh20) | 0.281162 |
| Cdh8 | cadherin 8(Cdh8) | -0.225513 |
| Cdh24 | cadherin-like 24(Cdh24) | 0.489012 |
| Ctnnal1 | catenin (cadherin associated protein), alpha-like 1(Ctnnal1) | 0.270472 |
| Cdon | cell adhesion molecule-related/down-regulated by oncogenes(Cdon) | 0.285659 |
| Cx3cl1 | chemokine (C-X3-C motif) ligand 1(Cx3cl1) | -0.186984 |
| Col5a1 | collagen, type V, alpha 1(Col5a1) | 0.188501 |
| Col6a1 | collagen, type VI, alpha 1(Col6a1) | -0.314202 |
| Col6a2 | collagen, type VI, alpha 2(Col6a2) | -0.668208 |
| Col19a1 | collagen, type XIX, alpha 1(Col19a1) | 0.324061 |
| Col15a1 | collagen, type XV, alpha 1(Col15a1) | 0.699774 |
| Cntn2 | contactin 2(Cntn2) | 0.483002 |
| Cntn3 | contactin 3(Cntn3) | 0.272548 |
| Cntnap5a | contactin associated protein-like 5A(Cntnap5a) | -0.244349 |
| Emb | embigin(Emb) | -0.326487 |
| Fn1 | fibronectin 1(Fn1) | 0.312679 |
| Flrt2 | fibronectin leucine rich transmembrane protein 2(Flrt2) | 0.207083 |
| Gpnmb | glycoprotein (transmembrane) nmb(Gpnmb) | -1.14048 |
| Igsf9b | immunoglobulin superfamily, member 9B(Igsf9b) | -0.206615 |
| Itgav | integrin alpha V(Itgav) | 0.229906 |
| Lama5 | laminin, alpha 5(Lama5) | 0.20888 |
| Lamc2 | laminin, gamma 2(Lamc2) | -0.526442 |
| Lmln | leishmanolysin-like (metallopeptidase M8 family)(Lmln) | 0.211086 |
| Megf10 | multiple EGF-like-domains 10(Megf10) | 0.254789 |
| Nov | nephroblastoma overexpressed gene(Nov) | -0.180737 |
| Negr1 | neuronal growth regulator 1(Negr1) | -0.180157 |
| Parvb | parvin, beta(Parvb) | -0.268398 |
| Pnn | pinin(Pnn) | 0.181257 |
| Pkp4 | plakophilin 4(Pkp4) | -0.250498 |
| Ptpru | protein tyrosine phosphatase, receptor type, U(Ptpru) | 0.2274 |
| Pcdh19 | protocadherin 19(Pcdh19) | 0.217846 |
| Ret | ret proto-oncogene(Ret) | -0.284101 |
| Thbs3 | thrombospondin 3(Thbs3) | 0.384423 |
| Thy1 | thymus cell antigen 1, theta(Thy1) | -0.227688 |
| Zyx | zyxin(Zyx) | -0.210285 |
| Atp1b2 | ATPase, Na+/K+ transporting, beta 2 polypeptide(Atp1b2) | -0.209579 |
| **Positive regulation of cytosolic calcium ion concentration** | | |
| **Symbol** | **Gene name** | **Log2 (fold change)** |
| Cd24a | CD24a antigen(Cd24a) | 0.208692 |
| Gpr6 | G protein-coupled receptor 6(Gpr6) | -0.693168 |
| Gpr3 | G-protein coupled receptor 3(Gpr3) | -0.379597 |
| Swap70 | SWA-70 protein(Swap70) | 0.22232 |
| Adcy5 | adenylate cyclase 5(Adcy5) | 0.18128 |
| Adcyap1 | adenylate cyclase activating polypeptide 1(Adcyap1) | -0.364772 |
| Adra1b | adrenergic receptor, alpha 1b(Adra1b) | -0.34772 |
| Adm | adrenomedullin(Adm) | -0.893345 |
| Agt | angiotensinogen (serpin peptidase inhibitor, clade A, member 8)(Agt) | -0.644347 |
| Cib2 | calcium and integrin binding family member 2(Cib2) | -0.291013 |
| Ccl28 | chemokine (C-C motif) ligand 28(Ccl28) | -1.97529 |
| Cckbr | cholecystokinin B receptor(Cckbr) | -0.562344 |
| F2r | coagulation factor II (thrombin) receptor(F2r) | -0.221243 |
| Crhr1 | corticotropin releasing hormone receptor 1(Crhr1) | -0.311301 |
| Gja1 | gap junction protein, alpha 1(Gja1) | -0.196478 |
| Gck | glucokinase(Gck) | -0.33515 |
| Itgav | integrin alpha V(Itgav) | 0.229906 |
| Npy2r | neuropeptide Y receptor Y2(Npy2r) | -0.230785 |
| Oxtr | oxytocin receptor(Oxtr) | 0.267845 |
| Pth1r | parathyroid hormone 1 receptor(Pth1r) | -0.735892 |
| Pdgfra | platelet derived growth factor receptor, alpha polypeptide(Pdgfra) | 0.288914 |
| P2ry1 | purinergic receptor P2Y, G-protein coupled 1(P2ry1) | -0.317979 |
| P2ry2 | purinergic receptor P2Y, G-protein coupled 2(P2ry2) | -0.415118 |
| Trhr | thyrotropin releasing hormone receptor(Trhr) | -0.253903 |
| Trpc6 | transient receptor potential cation channel, subfamily C, member 6(Trpc6) | -0.440269 |
| **Nervous system development** | | |
| **Symbol** | **Gene name** | **Log2 (fold change)** |
| Cables1 | CDK5 and Abl enzyme substrate 1(Cables1) | -0.281238 |
| Epha8 | Eph receptor A8(Epha8) | 0.320632 |
| Gpsm1 | G-protein signalling modulator 1 (AGS3-like, C. elegans)(Gpsm1) | 0.267821 |
| Lhx6 | LIM homeobox protein 6(Lhx6) | 0.341398 |
| Mdga1 | MAM domain containing glycosylphosphatidylinositol anchor 1(Mdga1) | 0.389912 |
| Rapgef2 | Rap guanine nucleotide exchange factor (GEF) 2(Rapgef2) | 0.18849 |
| Srgap2 | SLIT-ROBO Rho GTPase activating protein 2(Srgap2) | 0.206012 |
| Sox11 | SRY (sex determining region Y)-box 11(Sox11) | 0.227096 |
| Adcyap1 | adenylate cyclase activating polypeptide 1(Adcyap1) | -0.364772 |
| Arx | aristaless related homeobox(Arx) | 0.305155 |
| Bdnf | brain derived neurotrophic factor(Bdnf) | -0.470373 |
| Cdc20 | cell division cycle 20(Cdc20) | 0.353063 |
| Chn1 | chimerin 1(Chn1) | -0.313646 |
| Crmp1 | collapsin response mediator protein 1(Crmp1) | 0.185276 |
| Cntn3 | contactin 3(Cntn3) | 0.272548 |
| Dpysl3 | dihydropyrimidinase-like 3(Dpysl3) | 0.308843 |
| Dpysl5 | dihydropyrimidinase-like 5(Dpysl5) | 0.227129 |
| Dcx | doublecortin(Dcx) | 0.261673 |
| Ect2 | ect2 oncogene(Ect2) | 0.250271 |
| Efnb1 | ephrin B1(Efnb1) | 0.322353 |
| Grik1 | glutamate receptor, ionotropic, kainate 1(Grik1) | 0.285063 |
| Hdac9 | histone deacetylase 9(Hdac9) | -0.278945 |
| Islr2 | immunoglobulin superfamily containing leucine-rich repeat 2(Islr2) | 0.194081 |
| Igsf9b | immunoglobulin superfamily, member 9B(Igsf9b) | -0.206615 |
| Itm2c | integral membrane protein 2C(Itm2c) | -0.254432 |
| Mtss1 | metastasis suppressor 1(Mtss1) | 0.190135 |
| Myt1 | myelin transcription factor 1(Myt1) | 0.266333 |
| Nes | nestin(Nes) | 0.370533 |
| Nrn1 | neuritin 1(Nrn1) | -0.308931 |
| Nav1 | neuron navigator 1(Nav1) | 0.242038 |
| Nrbp2 | nuclear receptor binding protein 2(Nrbp2) | 0.265687 |
| Nr4a2 | nuclear receptor subfamily 4, group A, member 2(Nr4a2) | -0.279153 |
| Olig2 | oligodendrocyte transcription factor 2(Olig2) | 0.245334 |
| Ophn1 | oligophrenin 1(Ophn1) | 0.20499 |
| Plxnb3 | plexin B3(Plxnb3) | -0.774452 |
| Ret | ret proto-oncogene(Ret) | -0.284101 |
| Sema3e | sema domain, immunoglobulin domain (Ig), short basic domain, secreted, (semaphorin) 3E(Sema3e) | -0.349043 |
| Sema4b | sema domain, immunoglobulin domain (Ig), transmembrane domain (TM) and short cytoplasmic domain, (semaphorin) 4B(Sema4b) | 0.206854 |
| Scn2b | sodium channel, voltage-gated, type II, beta(Scn2b) | -0.190638 |
| Stmn1 | stathmin 1(Stmn1) | 0.194703 |
| Zic1 | zinc finger protein of the cerebellum 1(Zic1) | -0.346633 |
| **Negative regulation of neuron apoptotic process** | | |
| **Symbol** | **Gene name** | **Log2 (fold change)** |
| Bok | BCL2-related ovarian killer(Bok) | -0.23007 |
| Cebpb | CCAAT/enhancer binding protein (C/EBP), beta(Cebpb) | -0.387441 |
| Cited1 | Cbp/p300-interacting transactivator with Glu/Asp-rich carboxy-terminal domain 1(Cited1) | -0.51851 |
| Faim2 | Fas apoptotic inhibitory molecule 2(Faim2) | -0.265052 |
| Pcp4 | Purkinje cell protein 4(Pcp4) | -0.301547 |
| Xrcc2 | X-ray repair complementing defective repair in Chinese hamster cells 2(Xrcc2) | 0.269076 |
| Adora2a | adenosine A2a receptor(Adora2a) | -0.284827 |
| Agt | angiotensinogen (serpin peptidase inhibitor, clade A, member 8)(Agt) | -0.644347 |
| Birc5 | baculoviral IAP repeat-containing 5(Birc5) | 0.315857 |
| Bdnf | brain derived neurotrophic factor(Bdnf) | -0.470373 |
| Cln3 | ceroid lipofuscinosis, neuronal 3, juvenile (Batten, Spielmeyer-Vogt disease)(Cln3) | -0.275845 |
| F2r | coagulation factor II (thrombin) receptor(F2r) | -0.221243 |
| Dlx1 | distal-less homeobox 1(Dlx1) | 0.38264 |
| Draxin | dorsal inhibitory axon guidance protein(Draxin) | 0.238541 |
| Itsn1 | intersectin 1 (SH3 domain protein 1A)(Itsn1) | 0.274784 |
| Kif14 | kinesin family member 14(Kif14) | 0.288734 |
| Lgmn | legumain(Lgmn) | -0.203377 |
| Mt1 | metallothionein 1(Mt1) | -0.217934 |
| Msh2 | mutS homolog 2(Msh2) | -0.213602 |
| Nes | nestin(Nes) | 0.370533 |
| Nefl | neurofilament, light polypeptide(Nefl) | -0.24112 |
| Nrbp2 | nuclear receptor binding protein 2(Nrbp2) | 0.265687 |
| Nr4a2 | nuclear receptor subfamily 4, group A, member 2(Nr4a2) | -0.279153 |
| **Positive regulation of cell migration** | | |
| **Symbol** | **Gene name** | **Log2 (fold change)** |
| Ets1 | E26 avian leukemia oncogene 1, 5' domain(Ets1) | 0.253717 |
| Apc | adenomatosis polyposis coli(Apc) | 0.311095 |
| Adra2a | adrenergic receptor, alpha 2a(Adra2a) | 0.205162 |
| Bmp4 | bone morphogenetic protein 4(Bmp4) | -0.338676 |
| Cx3cl1 | chemokine (C-X3-C motif) ligand 1(Cx3cl1) | -0.186984 |
| F2r | coagulation factor II (thrombin) receptor(F2r) | -0.221243 |
| Egfr | epidermal growth factor receptor(Egfr) | 0.283762 |
| Fn1 | fibronectin 1(Fn1) | 0.312679 |
| Gcnt2 | glucosaminyl (N-acetyl) transferase 2, I-branching enzyme(Gcnt2) | -0.495263 |
| Gpnmb | glycoprotein (transmembrane) nmb(Gpnmb) | -1.14048 |
| Hbegf | heparin-binding EGF-like growth factor(Hbegf) | 0.270298 |
| Hgf | hepatocyte growth factor(Hgf) | 0.423115 |
| Itgav | integrin alpha V(Itgav) | 0.229906 |
| Lamc2 | laminin, gamma 2(Lamc2) | -0.526442 |
| Mmp14 | matrix metallopeptidase 14 (membrane-inserted)(Mmp14) | 0.207178 |
| Map2k1 | mitogen-activated protein kinase kinase 1(Map2k1) | -0.23032 |
| Myadm | myeloid-associated differentiation marker(Myadm) | -0.290833 |
| Myo1c | myosin IC(Myo1c) | 0.212708 |
| Mylk | myosin, light polypeptide kinase(Mylk) | -0.397011 |
| Pdgfra | platelet derived growth factor receptor, alpha polypeptide(Pdgfra) | 0.288914 |
| Pdgfd | platelet-derived growth factor, D polypeptide(Pdgfd) | -0.561035 |
| Rack1 | receptor for activated C kinase 1(Rack1) | 1.56605 |
| Ret | ret proto-oncogene(Ret) | -0.284101 |
| Sema3e | sema domain, immunoglobulin domain (Ig), short basic domain, secreted, (semaphorin) 3E(Sema3e) | -0.349043 |
| Sema3f | sema domain, immunoglobulin domain (Ig), short basic domain, secreted, (semaphorin) 3F(Sema3f) | 0.365773 |
| Sema4b | sema domain, immunoglobulin domain (Ig), transmembrane domain (TM) and short cytoplasmic domain, (semaphorin) 4B(Sema4b) | 0.206854 |
| **Cell cycle** | | |
| **Symbol** | **Gene name** | **Log2 (fold change)** |
| Bub1 | BUB1, mitotic checkpoint serine/threonine kinase(Bub1) | 0.446547 |
| Bub1b | BUB1B, mitotic checkpoint serine/threonine kinase(Bub1b) | 0.230577 |
| Cdc14a | CDC14 cell division cycle 14A(Cdc14a) | 0.264674 |
| Cables1 | CDK5 and Abl enzyme substrate 1(Cables1) | -0.281238 |
| Dbf4 | DBF4 zinc finger(Dbf4) | 0.281871 |
| Fbxo5 | F-box protein 5(Fbxo5) | 0.32071 |
| Fancd2 | Fanconi anemia, complementation group D2(Fancd2) | 0.242582 |
| Hjurp | Holliday junction recognition protein(Hjurp) | 0.274537 |
| Mad2l1 | MAD2 mitotic arrest deficient-like 1(Mad2l1) | 0.23892 |
| Nek3 | NIMA (never in mitosis gene a)-related expressed kinase 3(Nek3) | 0.481955 |
| Nek4 | NIMA (never in mitosis gene a)-related expressed kinase 4(Nek4) | -0.209945 |
| Phf13 | PHD finger protein 13(Phf13) | -0.239565 |
| Spc24 | SPC24, NDC80 kinetochore complex component, homolog (S. cerevisiae)(Spc24) | 0.302314 |
| Mki67 | antigen identified by monoclonal antibody Ki 67(Mki67) | 0.323317 |
| Aspm | asp (abnormal spindle)-like, microcephaly associated (Drosophila)(Aspm) | 0.255128 |
| Birc5 | baculoviral IAP repeat-containing 5(Birc5) | 0.315857 |
| Brinp1 | bone morphogenic protein/retinoic acid inducible neural specific 1(Brinp1) | -0.273883 |
| Cdc20 | cell division cycle 20(Cdc20) | 0.353063 |
| Cdc6 | cell division cycle 6(Cdc6) | 0.476019 |
| Cdca5 | cell division cycle associated 5(Cdca5) | 0.531707 |
| Cntrob | centrobin, centrosomal BRCA2 interacting protein(Cntrob) | 0.254391 |
| Cenpt | centromere protein T(Cenpt) | 0.307368 |
| Chek1 | checkpoint kinase 1(Chek1) | 0.258348 |
| Chaf1b | chromatin assembly factor 1, subunit B (p60)(Chaf1b) | 0.557127 |
| Clspn | claspin(Clspn) | 0.324617 |
| Ccna2 | cyclin A2(Ccna2) | 0.261407 |
| Ccnb2 | cyclin B2(Ccnb2) | 0.257691 |
| Ccnd1 | cyclin D1(Ccnd1) | 0.18533 |
| Ccnd2 | cyclin D2(Ccnd2) | 0.353105 |
| Ccne1 | cyclin E1(Ccne1) | -0.260069 |
| Cdkn1c | cyclin-dependent kinase inhibitor 1C (P57)(Cdkn1c) | -0.539388 |
| Ckap2 | cytoskeleton associated protein 2(Ckap2) | 0.243045 |
| Ect2 | ect2 oncogene(Ect2) | 0.250271 |
| Esco2 | establishment of sister chromatid cohesion N-acetyltransferase 2(Esco2) | 0.321325 |
| Ercc6l | excision repair cross-complementing rodent repair deficiency complementation group 6 like(Ercc6l) | 0.335378 |
| Fam83d | family with sequence similarity 83, member D(Fam83d) | 0.45442 |
| Kif20b | kinesin family member 20B(Kif20b) | 0.248936 |
| Kif23 | kinesin family member 23(Kif23) | 0.460478 |
| Knl1 | kinetochore scaffold 1(Knl1) | 0.52736 |
| Lmln | leishmanolysin-like (metallopeptidase M8 family)(Lmln) | 0.211086 |
| Mcm2 | minichromosome maintenance complex component 2(Mcm2) | 0.233332 |
| Mcm5 | minichromosome maintenance complex component 5(Mcm5) | 0.271372 |
| Mcm7 | minichromosome maintenance complex component 7(Mcm7) | 0.316533 |
| Mapk12 | mitogen-activated protein kinase 12(Mapk12) | -0.587148 |
| Msh2 | mutS homolog 2(Msh2) | -0.213602 |
| Mybl2 | myeloblastosis oncogene-like 2(Mybl2) | 0.389053 |
| Ncaph | non-SMC condensin I complex, subunit H(Ncaph) | 0.460362 |
| Ncapd3 | non-SMC condensin II complex, subunit D3(Ncapd3) | 0.226671 |
| Pttg1 | pituitary tumor-transforming gene 1(Pttg1) | -0.268946 |
| Plk1 | polo-like kinase 1(Plk1) | 0.270134 |
| Prc1 | protein regulator of cytokinesis 1(Prc1) | 0.250565 |
| Rack1 | receptor for activated C kinase 1(Rack1) | 1.56605 |
| Rgs2 | regulator of G-protein signaling 2(Rgs2) | -0.201031 |
| Txnip | thioredoxin interacting protein(Txnip) | -0.378476 |
| Usp2 | ubiquitin specific peptidase 2(Usp2) | -0.219692 |
| Usp22 | ubiquitin specific peptidase 22(Usp22) | 0.229866 |
| Zfp830 | zinc finger protein 830(Zfp830) | -0.295318 |
| **Cell division** | | |
| **Symbol** | **Gene name** | **Log2 (fold change)** |
| Bub1 | BUB1, mitotic checkpoint serine/threonine kinase(Bub1) | 0.446547 |
| Bub1b | BUB1B, mitotic checkpoint serine/threonine kinase(Bub1b) | 0.230577 |
| Cdc14a | CDC14 cell division cycle 14A(Cdc14a) | 0.264674 |
| Cables1 | CDK5 and Abl enzyme substrate 1(Cables1) | -0.281238 |
| Fbxo5 | F-box protein 5(Fbxo5) | 0.32071 |
| Mad2l1 | MAD2 mitotic arrest deficient-like 1(Mad2l1) | 0.23892 |
| Nek3 | NIMA (never in mitosis gene a)-related expressed kinase 3(Nek3) | 0.481955 |
| Nek4 | NIMA (never in mitosis gene a)-related expressed kinase 4(Nek4) | -0.209945 |
| Phf13 | PHD finger protein 13(Phf13) | -0.239565 |
| Spc24 | SPC24, NDC80 kinetochore complex component, homolog (S. cerevisiae)(Spc24) | 0.302314 |
| Aspm | asp (abnormal spindle)-like, microcephaly associated (Drosophila)(Aspm) | 0.255128 |
| Birc5 | baculoviral IAP repeat-containing 5(Birc5) | 0.315857 |
| Cdc20 | cell division cycle 20(Cdc20) | 0.353063 |
| Cdc6 | cell division cycle 6(Cdc6) | 0.476019 |
| Cdca5 | cell division cycle associated 5(Cdca5) | 0.531707 |
| Cntrob | centrobin, centrosomal BRCA2 interacting protein(Cntrob) | 0.254391 |
| Cenpt | centromere protein T(Cenpt) | 0.307368 |
| Ccna2 | cyclin A2(Ccna2) | 0.261407 |
| Ccnb2 | cyclin B2(Ccnb2) | 0.257691 |
| Ccnd1 | cyclin D1(Ccnd1) | 0.18533 |
| Ccnd2 | cyclin D2(Ccnd2) | 0.353105 |
| Ccne1 | cyclin E1(Ccne1) | -0.260069 |
| Cdk3-ps | cyclin-dependent kinase 3, pseudogene(Cdk3-ps) | 0.61584 |
| Dynlt1b | dynein light chain Tctex-type 1B(Dynlt1b) | -0.83493 |
| Ect2 | ect2 oncogene(Ect2) | 0.250271 |
| Ercc6l | excision repair cross-complementing rodent repair deficiency complementation group 6 like(Ercc6l) | 0.335378 |
| Fam83d | family with sequence similarity 83, member D(Fam83d) | 0.45442 |
| Kif14 | kinesin family member 14(Kif14) | 0.288734 |
| Kif20b | kinesin family member 20B(Kif20b) | 0.248936 |
| Kif23 | kinesin family member 23(Kif23) | 0.460478 |
| Knl1 | kinetochore scaffold 1(Knl1) | 0.52736 |
| Lmln | leishmanolysin-like (metallopeptidase M8 family)(Lmln) | 0.211086 |
| Mcm5 | minichromosome maintenance complex component 5(Mcm5) | 0.271372 |
| Ncaph | non-SMC condensin I complex, subunit H(Ncaph) | 0.460362 |
| Ncapd3 | non-SMC condensin II complex, subunit D3(Ncapd3) | 0.226671 |
| Pttg1 | pituitary tumor-transforming gene 1(Pttg1) | -0.268946 |
| Plk1 | polo-like kinase 1(Plk1) | 0.270134 |
| Prc1 | protein regulator of cytokinesis 1(Prc1) | 0.250565 |
| Zfp830 | zinc finger protein 830(Zfp830) | -0.295318 |
| **Cellular response to mechanical stimulus** | | |
| **Symbol** | **Gene name** | **Log2 (fold change)** |
| Cradd | CASP2 and RIPK1 domain containing adaptor with death domain(Cradd) | -0.40881 |
| Agt | angiotensinogen (serpin peptidase inhibitor, clade A, member 8)(Agt) | -0.644347 |
| Cnn2 | calponin 2(Cnn2) | -0.858707 |
| Casp2 | caspase 2(Casp2) | 0.310271 |
| Cav1 | caveolin 1, caveolae protein(Cav1) | -0.33376 |
| Chek1 | checkpoint kinase 1(Chek1) | 0.258348 |
| Cyba | cytochrome b-245, alpha polypeptide(Cyba) | -0.513863 |
| Egfr | epidermal growth factor receptor(Egfr) | 0.283762 |
| Gja1 | gap junction protein, alpha 1(Gja1) | -0.196478 |
| Map3k1 | mitogen-activated protein kinase kinase kinase 1(Map3k1) | 0.19665 |
| Nos1 | nitric oxide synthase 1, neuronal(Nos1) | 0.450857 |
| Kcnj2 | potassium inwardly-rectifying channel, subfamily J, member 2(Kcnj2) | -0.526151 |
| Ptgs2 | prostaglandin-endoperoxide synthase 2(Ptgs2) | -0.380668 |
| Tlr3 | toll-like receptor 3(Tlr3) | -0.27512 |
| **Mitotic nuclear division** | | |
| **Symbol** | **Gene name** | **Log2 (fold change)** |
| Bub1 | BUB1, mitotic checkpoint serine/threonine kinase(Bub1) | 0.446547 |
| Bub1b | BUB1B, mitotic checkpoint serine/threonine kinase(Bub1b) | 0.230577 |
| Fbxo5 | F-box protein 5(Fbxo5) | 0.32071 |
| Gem | GTP binding protein (gene overexpressed in skeletal muscle)(Gem) | -0.263064 |
| Mad2l1 | MAD2 mitotic arrest deficient-like 1(Mad2l1) | 0.23892 |
| Nek3 | NIMA (never in mitosis gene a)-related expressed kinase 3(Nek3) | 0.481955 |
| Nek4 | NIMA (never in mitosis gene a)-related expressed kinase 4(Nek4) | -0.209945 |
| Phf13 | PHD finger protein 13(Phf13) | -0.239565 |
| Spc24 | SPC24, NDC80 kinetochore complex component, homolog (S. cerevisiae)(Spc24) | 0.302314 |
| Aspm | asp (abnormal spindle)-like, microcephaly associated (Drosophila)(Aspm) | 0.255128 |
| Birc5 | baculoviral IAP repeat-containing 5(Birc5) | 0.315857 |
| Cdc20 | cell division cycle 20(Cdc20) | 0.353063 |
| Cdc6 | cell division cycle 6(Cdc6) | 0.476019 |
| Cdca5 | cell division cycle associated 5(Cdca5) | 0.531707 |
| Cenpt | centromere protein T(Cenpt) | 0.307368 |
| Ccna2 | cyclin A2(Ccna2) | 0.261407 |
| Ccnb2 | cyclin B2(Ccnb2) | 0.257691 |
| Cdk3-ps | cyclin-dependent kinase 3, pseudogene(Cdk3-ps) | 0.61584 |
| Dynlt1b | dynein light chain Tctex-type 1B(Dynlt1b) | -0.83493 |
| Ercc6l | excision repair cross-complementing rodent repair deficiency complementation group 6 like(Ercc6l) | 0.335378 |
| Fam83d | family with sequence similarity 83, member D(Fam83d) | 0.45442 |
| Kif20b | kinesin family member 20B(Kif20b) | 0.248936 |
| Kif23 | kinesin family member 23(Kif23) | 0.460478 |
| Knl1 | kinetochore scaffold 1(Knl1) | 0.52736 |
| Lmln | leishmanolysin-like (metallopeptidase M8 family)(Lmln) | 0.211086 |
| Map2k1 | mitogen-activated protein kinase kinase 1(Map2k1) | -0.23032 |
| Ncaph | non-SMC condensin I complex, subunit H(Ncaph) | 0.460362 |
| Ncapd3 | non-SMC condensin II complex, subunit D3(Ncapd3) | 0.226671 |
| Pttg1 | pituitary tumor-transforming gene 1(Pttg1) | -0.268946 |
| Plk1 | polo-like kinase 1(Plk1) | 0.270134 |
| Zfp830 | zinc finger protein 830(Zfp830) | -0.295318 |
| **Regulation of G-protein coupled receptor protein signaling pathway** | | |
| **Symbol** | **Gene name** | **Log2 (fold change)** |
| Gpsm1 | G-protein signalling modulator 1 (AGS3-like, C. elegans)(Gpsm1) | 0.267821 |
| Adcyap1 | adenylate cyclase activating polypeptide 1(Adcyap1) | -0.364772 |
| Arrb1 | arrestin, beta 1(Arrb1) | 0.191243 |
| Dynlt1b | dynein light chain Tctex-type 1B(Dynlt1b) | -0.83493 |
| Homer2 | homer scaffolding protein 2(Homer2) | 0.246626 |
| Ramp1 | receptor (calcitonin) activity modifying protein 1(Ramp1) | -0.319947 |
| Ramp2 | receptor (calcitonin) activity modifying protein 2(Ramp2) | -0.524774 |
| Ramp3 | receptor (calcitonin) activity modifying protein 3(Ramp3) | -0.585706 |
| Rgs2 | regulator of G-protein signaling 2(Rgs2) | -0.201031 |
| **Positive regulation of GTPase activity** | | |
| **Symbol** | **Gene name** | **Log2 (fold change)** |
| Dennd1b | DENN/MADD domain containing 1B(Dennd1b) | 0.226369 |
| Rasgrp1 | RAS guanyl releasing protein 1(Rasgrp1) | -0.302872 |
| Rapgef2 | Rap guanine nucleotide exchange factor (GEF) 2(Rapgef2) | 0.18849 |
| Rapgef3 | Rap guanine nucleotide exchange factor (GEF) 3(Rapgef3) | 0.305563 |
| S100a10 | S100 calcium binding protein A10 (calpactin)(S100a10) | -0.323893 |
| Srgap2 | SLIT-ROBO Rho GTPase activating protein 2(Srgap2) | 0.206012 |
| Adcyap1 | adenylate cyclase activating polypeptide 1(Adcyap1) | -0.364772 |
| Afdn | afadin, adherens junction formation factor(Afdn) | 0.202055 |
| Bcas3 | breast carcinoma amplified sequence 3(Bcas3) | -0.206325 |
| Cx3cl1 | chemokine (C-X3-C motif) ligand 1(Cx3cl1) | -0.186984 |
| Ect2 | ect2 oncogene(Ect2) | 0.250271 |
| Lamtor5 | late endosomal/lysosomal adaptor, MAPK and MTOR activator 5(Lamtor5) | -0.245778 |
| Map2k1 | mitogen-activated protein kinase kinase 1(Map2k1) | -0.23032 |
| Pkp4 | plakophilin 4(Pkp4) | -0.250498 |
| Rgl3 | ral guanine nucleotide dissociation stimulator-like 3(Rgl3) | -0.361658 |
| Rack1 | receptor for activated C kinase 1(Rack1) | 1.56605 |
| Rgs2 | regulator of G-protein signaling 2(Rgs2) | -0.201031 |
| Rgs4 | regulator of G-protein signaling 4(Rgs4) | -0.230529 |
| Rgs10 | regulator of G-protein signalling 10(Rgs10) | -0.300101 |
| Thy1 | thymus cell antigen 1, theta(Thy1) | -0.227688 |
| **Positive chemotaxis** | | |
| **Symbol** | **Gene name** | **Log2 (fold change)** |
| Bmp4 | bone morphogenetic protein 4(Bmp4) | -0.338676 |
| Lgals3 | lectin, galactose binding, soluble 3(Lgals3) | -0.512762 |
| Met | met proto-oncogene(Met) | 0.198161 |
| Plxnb3 | plexin B3(Plxnb3) | -0.774452 |
| Scrib | scribbled planar cell polarity(Scrib) | 0.296674 |
| Scg2 | secretogranin II(Scg2) | -0.214127 |
| **positive regulation of endothelial cell proliferation** | | |
| **Symbol** | **Gene name** | **Log2 (fold change)** |
| Bmp4 | bone morphogenetic protein 4(Bmp4) | -0.338676 |
| Cav1 | caveolin 1, caveolae protein(Cav1) | -0.33376 |
| Cyba | cytochrome b-245, alpha polypeptide(Cyba) | -0.513863 |
| Egr3 | early growth response 3(Egr3) | -0.337981 |
| Hmgb2 | high mobility group box 2(Hmgb2) | 0.473187 |
| Mydgf | myeloid derived growth factor(Mydgf) | -0.295379 |
| Nr4a1 | nuclear receptor subfamily 4, group A, member 1(Nr4a1) | -0.315837 |
| Plxnb3 | plexin B3(Plxnb3) | -0.774452 |
| Prox1 | prospero homeobox 1(Prox1) | -0.295314 |
| Scg2 | secretogranin II(Scg2) | -0.214127 |
| Vip | vasoactive intestinal polypeptide(Vip) | -0.406533 |
| Vash2 | vasohibin 2(Vash2) | 0.225852 |
| **Regulation of ion transmembrane transport** | | |
| **Symbol** | **Gene name** | **Log2 (fold change)** |
| Kcnip4 | Kv channel interacting protein 4(Kcnip4) | -0.405868 |
| Cacng3 | calcium channel, voltage-dependent, gamma subunit 3(Cacng3) | -0.419447 |
| Cacng4 | calcium channel, voltage-dependent, gamma subunit 4(Cacng4) | 0.234454 |
| Cacng5 | calcium channel, voltage-dependent, gamma subunit 5(Cacng5) | -0.447424 |
| Clic4 | chloride intracellular channel 4 (mitochondrial)(Clic4) | 0.288359 |
| Kcnv1 | potassium channel, subfamily V, member 1(Kcnv1) | -0.401908 |
| Kcnj2 | potassium inwardly-rectifying channel, subfamily J, member 2(Kcnj2) | -0.526151 |
| Kcnj9 | potassium inwardly-rectifying channel, subfamily J, member 9(Kcnj9) | -0.268833 |
| Kcnb2 | potassium voltage gated channel, Shab-related subfamily, member 2(Kcnb2) | -0.275121 |
| Kcnab1 | potassium voltage-gated channel, shaker-related subfamily, beta member 1(Kcnab1) | -0.230488 |
| Kcnf1 | potassium voltage-gated channel, subfamily F, member 1(Kcnf1) | -0.379904 |
| Kcng1 | potassium voltage-gated channel, subfamily G, member 1(Kcng1) | -0.192533 |
| Kcnh2 | potassium voltage-gated channel, subfamily H (eag-related), member 2(Kcnh2) | -0.220063 |
| Kcnh7 | potassium voltage-gated channel, subfamily H (eag-related), member 7(Kcnh7) | -0.285739 |
| Kcnq3 | potassium voltage-gated channel, subfamily Q, member 3(Kcnq3) | -0.249528 |
| Scn2b | sodium channel, voltage-gated, type II, beta(Scn2b) | -0.190638 |
| Stom | stomatin(Stom) | -0.272398 |
| Tmem109 | transmembrane protein 109(Tmem109) | -0.231289 |
| **Response to estradiol** | | |
| **Symbol** | **Gene name** | **Log2 (fold change)** |
| Ets1 | E26 avian leukemia oncogene 1, 5' domain(Ets1) | 0.253717 |
| Bmp7 | bone morphogenetic protein 7(Bmp7) | 0.244832 |
| Cryab | crystallin, alpha B(Cryab) | -0.391796 |
| Ccnd1 | cyclin D1(Ccnd1) | 0.18533 |
| Gjb2 | gap junction protein, beta 2(Gjb2) | -0.873347 |
| Igfbp2 | insulin-like growth factor binding protein 2(Igfbp2) | 0.316157 |
| Ifi27 | interferon, alpha-inducible protein 27(Ifi27) | -0.447247 |
| Mapk15 | mitogen-activated protein kinase 15(Mapk15) | 0.660586 |
| Oxtr | oxytocin receptor(Oxtr) | 0.267845 |
| Ptgs2 | prostaglandin-endoperoxide synthase 2(Ptgs2) | -0.380668 |
| Ramp2 | receptor (calcitonin) activity modifying protein 2(Ramp2) | -0.524774 |
| Slc6a1 | solute carrier family 6 (neurotransmitter transporter, GABA), member 1(Slc6a1) | 0.201705 |
| Tacr3 | tachykinin receptor 3(Tacr3) | -0.345044 |
| Txnip | thioredoxin interacting protein(Txnip) | -0.378476 |
| Tfpi | tissue factor pathway inhibitor(Tfpi) | -0.683443 |
