## Supplementary Table 3 for "Computational analysis of transcriptome signature repurposes low dose trifluoperazine for the treatment of fragile X syndrome in mouse model"

| **Cell cycle** | | |
| --- | --- | --- |
| **Symbol** | **Gene name** | **Log2 (fold change)** |
| Bub1 | BUB1, mitotic checkpoint serine/threonine kinase(Bub1) | 0.446547 |
| Bub1b | BUB1B, mitotic checkpoint serine/threonine kinase(Bub1b) | 0.230577 |
| Cdc14a | CDC14 cell division cycle 14A(Cdc14a) | 0.264674 |
| Dbf4 | DBF4 zinc finger(Dbf4) | 0.281871 |
| Mad2l1 | MAD2 mitotic arrest deficient-like 1(Mad2l1) | 0.23892 |
| Cdc20 | cell division cycle 20(Cdc20) | 0.353063 |
| Cdc6 | cell division cycle 6(Cdc6) | 0.476019 |
| Chek1 | checkpoint kinase 1(Chek1) | 0.258348 |
| Ccna2 | cyclin A2(Ccna2) | 0.261407 |
| Ccnb2 | cyclin B2(Ccnb2) | 0.257691 |
| Ccnd1 | cyclin D1(Ccnd1) | 0.18533 |
| Ccnd2 | cyclin D2(Ccnd2) | 0.353105 |
| Ccne1 | cyclin E1(Ccne1) | -0.260069 |
| Cdkn1c | cyclin-dependent kinase inhibitor 1C (P57)(Cdkn1c) | -0.539388 |
| Gadd45g | growth arrest and DNA-damage-inducible 45 gamma(Gadd45g) | -0.254925 |
| Mcm2 | minichromosome maintenance complex component 2(Mcm2) | 0.233332 |
| Mcm5 | minichromosome maintenance complex component 5(Mcm5) | 0.271372 |
| Mcm7 | minichromosome maintenance complex component 7(Mcm7) | 0.316533 |
| Pttg1 | pituitary tumor-transforming gene 1(Pttg1) | -0.268946 |
| Plk1 | polo-like kinase 1(Plk1) | 0.270134 |
| **MAPK signaling pathway** | | |
| **Symbol** | **Gene name** | **Log2 (fold change)** |
| Elk1 | ELK1, member of ETS oncogene family(Elk1) | 0.190294 |
| Rasgrp1 | RAS guanyl releasing protein 1(Rasgrp1) | -0.302872 |
| Rapgef2 | Rap guanine nucleotide exchange factor (GEF) 2(Rapgef2) | 0.18849 |
| Arrb1 | arrestin, beta 1(Arrb1) | 0.191243 |
| Relb | avian reticuloendotheliosis viral (v-rel) oncogene related B(Relb) | -0.481266 |
| Bdnf | brain derived neurotrophic factor(Bdnf) | -0.470373 |
| Cacna1i | calcium channel, voltage-dependent, alpha 1I subunit(Cacna1i) | -0.194993 |
| Cacng3 | calcium channel, voltage-dependent, gamma subunit 3(Cacng3) | -0.419447 |
| Cacng4 | calcium channel, voltage-dependent, gamma subunit 4(Cacng4) | 0.234454 |
| Cacng5 | calcium channel, voltage-dependent, gamma subunit 5(Cacng5) | -0.44742 |
| Dusp4 | dual specificity phosphatase 4(Dusp4) | -0.315051 |
| Dusp7 | dual specificity phosphatase 7(Dusp7) | -0.208387 |
| Egfr | epidermal growth factor receptor(Egfr) | 0.283762 |
| Fgf9 | fibroblast growth factor 9(Fgf9) | -0.394831 |
| Flna | filamin, alpha(Flna) | 0.203053 |
| Gadd45g | growth arrest and DNA-damage-inducible 45 gamma(Gadd45g) | -0.254925 |
| Hspa2 | heat shock protein 2(Hspa2) | -0.320115 |
| Mapk12 | mitogen-activated protein kinase 12(Mapk12) | -0.587148 |
| Map2k1 | mitogen-activated protein kinase kinase 1(Map2k1) | -0.23032 |
| Map3k1 | mitogen-activated protein kinase kinase kinase 1(Map3k1) | 0.19665 |
| Map4k4 | mitogen-activated protein kinase kinase kinase kinase 4(Map4k4) | 0.193833 |
| Nr4a1 | nuclear receptor subfamily 4, group A, member 1(Nr4a1) | -0.315837 |
| Pdgfra | platelet derived growth factor receptor, alpha polypeptide(Pdgfra) | 0.288914 |
| Rras | related RAS viral (r-ras) oncogene(Rras) | -0.397615 |
| Rps6ka5 | ribosomal protein S6 kinase, polypeptide 5(Rps6ka5) | 0.239409 |
| Stmn1 | stathmin 1(Stmn1) | 0.194703 |
| Tgfbr2 | transforming growth factor, beta receptor II(Tgfbr2) | -0.402446 |
| **Focal adhesion** | | |
| **Symbol** | **Gene name** | **Log2 (fold change)** |
| Elk1 | ELK1, member of ETS oncogene family(Elk1) | 0.190294 |
| Cav1 | caveolin 1, caveolae protein(Cav1) | -0.33376 |
| Col2a1 | collagen, type II, alpha 1(Col2a1) | -0.495529 |
| Col5a1 | collagen, type V, alpha 1(Col5a1) | 0.188501 |
| Col6a1 | collagen, type VI, alpha 1(Col6a1) | -0.314202 |
| Col6a2 | collagen, type VI, alpha 2(Col6a2) | -0.668208 |
| Ccnd1 | cyclin D1(Ccnd1) | 0.18533 |
| Ccnd2 | cyclin D2(Ccnd2) | 0.353105 |
| Egfr | epidermal growth factor receptor(Egfr) | 0.283762 |
| Fn1 | fibronectin 1(Fn1) | 0.312679 |
| Flna | filamin, alpha(Flna) | 0.203053 |
| Hgf | hepatocyte growth factor(Hgf) | 0.423115 |
| Itgav | integrin alpha V(Itgav) | 0.229906 |
| Lama5 | laminin, alpha 5(Lama5) | 0.20888 |
| Lamc2 | laminin, gamma 2(Lamc2) | -0.526442 |
| Met | met proto-oncogene(Met) | 0.198161 |
| Map2k1 | mitogen-activated protein kinase kinase 1(Map2k1) | -0.23032 |
| Mylk | myosin, light polypeptide kinase(Mylk) | -0.397011 |
| Parvb | parvin, beta(Parvb) | -0.268398 |
| Pdgfra | platelet derived growth factor receptor, alpha polypeptide(Pdgfra) | 0.288914 |
| Pdgfd | platelet-derived growth factor, D polypeptide(Pdgfd) | -0.561035 |
| Thbs3 | thrombospondin 3(Thbs3) | 0.384423 |
| Zyx | zyxin(Zyx) | -0.210285 |
| **p53 signaling pathway** | | |
| **Symbol** | **Gene name** | **Log2 (fold change)** |
| Cd82 | CD82 antigen(Cd82) | -0.584949 |
| Gtse1 | G two S phase expressed protein 1(Gtse1) | 0.388941 |
| Chek1 | checkpoint kinase 1(Chek1) | 0.258348 |
| Ccnb2 | cyclin B2(Ccnb2) | 0.257691 |
| Ccnd1 | cyclin D1(Ccnd1) | 0.18533 |
| Ccnd2 | cyclin D2(Ccnd2) | 0.353105 |
| Ccne1 | cyclin E1(Ccne1) | -0.260069 |
| Gadd45g | growth arrest and DNA-damage-inducible 45 gamma(Gadd45g) | -0.254925 |
| Pidd1 | p53 induced death domain protein 1(Pidd1) | 0.46691 |
| Rrm2 | ribonucleotide reductase M2(Rrm2) | 0.23946 |
| Shisa5 | shisa family member 5(Shisa5) | -0.260195 |
| **Calcium signaling pathway** | | |
| **Symbol** | **Gene name** | **Log2 (fold change)** |
| Htr6 | 5-hydroxytryptamine (serotonin) receptor 6(Htr6) | 0.400014 |
| Atp2b4 | ATPase, Ca++ transporting, plasma membrane 4(Atp2b4) | -0.209647 |
| Adora2a | adenosine A2a receptor(Adora2a) | -0.284827 |
| Adcy8 | adenylate cyclase 8(Adcy8) | -0.219063 |
| Adra1b | adrenergic receptor, alpha 1b(Adra1b) | -0.34772 |
| Cacna1i | calcium channel, voltage-dependent, alpha 1I subunit(Cacna1i) | -0.194993 |
| Cckbr | cholecystokinin B receptor(Cckbr) | -0.562344 |
| F2r | coagulation factor II (thrombin) receptor(F2r) | -0.221243 |
| Egfr | epidermal growth factor receptor(Egfr) | 0.283762 |
| Itpka | inositol 1,4,5-trisphosphate 3-kinase A(Itpka) | -0.304801 |
| Itpkb | inositol 1,4,5-trisphosphate 3-kinase B(Itpkb) | 0.200464 |
| Mylk | myosin, light polypeptide kinase(Mylk) | -0.397011 |
| Nos1 | nitric oxide synthase 1, neuronal(Nos1) | 0.450857 |
| Oxtr | oxytocin receptor(Oxtr) | 0.267845 |
| Plcd3 | phospholipase C, delta 3(Plcd3) | -0.264772 |
| Pdgfra | platelet derived growth factor receptor, alpha polypeptide(Pdgfra) | 0.288914 |
| P2rx3 | purinergic receptor P2X, ligand-gated ion channel, 3(P2rx3) | -0.297367 |
| P2rx6 | purinergic receptor P2X, ligand-gated ion channel, 6(P2rx6) | -0.302309 |
| Tacr3 | tachykinin receptor 3(Tacr3) | -0.345044 |
| Trhr | thyrotropin releasing hormone receptor(Trhr) | -0.253903 |
| **Amino sugar and nucleotide sugar metabolism** | | |
| **Symbol** | **Gene name** | **Log2 (fold change)** |
| Gmds | GDP-mannose 4, 6-dehydratase(Gmds) | -0.273588 |
| Nans | N-acetylneuraminic acid synthase (sialic acid synthase)(Nans) | -0.28157 |
| Cyb5r1 | cytochrome b5 reductase 1(Cyb5r1) | -0.333184 |
| Cyb5r2 | cytochrome b5 reductase 2(Cyb5r2) | -0.686573 |
| Fpgt | fucose-1-phosphate guanylyltransferase(Fpgt) | -0.231752 |
| Gck | glucokinase(Gck) | -0.33515 |
| Hexb | hexosaminidase B(Hexb) | -0.446539 |
| Renbp | renin binding protein(Renbp) | -0.62755 |
| **Cocaine addiction** | | |
| **Symbol** | **Gene name** | **Log2 (fold change)** |
| Gpsm1 | G-protein signalling modulator 1 (AGS3-like, C. elegans)(Gpsm1) | 0.267821 |
| Adcy5 | adenylate cyclase 5(Adcy5) | 0.18128 |
| Bdnf | brain derived neurotrophic factor(Bdnf) | -0.470373 |
| Grm2 | glutamate receptor, metabotropic 2(Grm2) | -0.408961 |
| Maob | monoamine oxidase B(Maob) | -0.607282 |
| Pdyn | prodynorphin(Pdyn) | -0.715908 |
| Ppp1r1b | protein phosphatase 1, regulatory (inhibitor) subunit 1B(Ppp1r1b) | 0.269958 |
| Slc18a2 | solute carrier family 18 (vesicular monoamine), member 2(Slc18a2) | -0.364674 |
| **Rap1 signaling pathway** | | |
| **Symbol** | **Gene name** | **Log2 (fold change)** |
| Rapgef2 | Rap guanine nucleotide exchange factor (GEF) 2(Rapgef2) | 0.18849 |
| Rapgef3 | Rap guanine nucleotide exchange factor (GEF) 3(Rapgef3) | 0.305563 |
| Adora2a | adenosine A2a receptor(Adora2a) | -0.284827 |
| Adcy5 | adenylate cyclase 5(Adcy5) | 0.18128 |
| Adcy8 | adenylate cyclase 8(Adcy8) | -0.219063 |
| Afdn | afadin, adherens junction formation factor(Afdn) | 0.202055 |
| F2r | coagulation factor II (thrombin) receptor(F2r) | -0.221243 |
| Dock4 | dedicator of cytokinesis 4(Dock4) | 0.203538 |
| Egfr | epidermal growth factor receptor(Egfr) | 0.283762 |
| Fgf9 | fibroblast growth factor 9(Fgf9) | -0.394831 |
| Hgf | hepatocyte growth factor(Hgf) | 0.423115 |
| Id1 | inhibitor of DNA binding 1(Id1) | -0.589363 |
| Lpar4 | lysophosphatidic acid receptor 4(Lpar4) | 0.46842 |
| Met | met proto-oncogene(Met) | 0.198161 |
| Mapk12 | mitogen-activated protein kinase 12(Mapk12) | -0.587148 |
| Map2k1 | mitogen-activated protein kinase kinase 1(Map2k1) | -0.23032 |
| Pdgfra | platelet derived growth factor receptor, alpha polypeptide(Pdgfra) | 0.288914 |
| Pdgfd | platelet-derived growth factor, D polypeptide(Pdgfd) | -0.561035 |
| P2ry1 | purinergic receptor P2Y, G-protein coupled 1(P2ry1) | -0.317979 |
| Rras | related RAS viral (r-ras) oncogene(Rras) | -0.397615 |
| **HTLV-I infection** | | |
| **Symbol** | **Gene name** | **Log2 (fold change)** |
| Bub1b | BUB1B, mitotic checkpoint serine/threonine kinase(Bub1b) | 0.230577 |
| Crtc2 | CREB regulated transcription coactivator 2(Crtc2) | 0.228122 |
| Ets1 | E26 avian leukemia oncogene 1, 5' domain(Ets1) | 0.253717 |
| Elk1 | ELK1, member of ETS oncogene family(Elk1) | 0.190294 |
| Mad2l1 | MAD2 mitotic arrest deficient-like 1(Mad2l1) | 0.23892 |
| Apc2 | adenomatosis polyposis coli 2(Apc2) | 0.236055 |
| Apc | adenomatosis polyposis coli(Apc) | 0.311095 |
| Adcy5 | adenylate cyclase 5(Adcy5) | 0.18128 |
| Adcy8 | adenylate cyclase 8(Adcy8) | -0.219063 |
| Relb | avian reticuloendotheliosis viral (v-rel) oncogene related B(Relb) | -0.481266 |
| Cdc20 | cell division cycle 20(Cdc20) | 0.353063 |
| Chek1 | checkpoint kinase 1(Chek1) | 0.258348 |
| Ccnd1 | cyclin D1(Ccnd1) | 0.18533 |
| Ccnd2 | cyclin D2(Ccnd2) | 0.353105 |
| Dvl1 | dishevelled segment polarity protein 1(Dvl1) | 0.183879 |
| H2-T24 | histocompatibility 2, T region locus 24(H2-T24) | -0.347115 |
| Map3k1 | mitogen-activated protein kinase kinase kinase 1(Map3k1) | 0.19665 |
| Mybl2 | myeloblastosis oncogene-like 2(Mybl2) | 0.389053 |
| Pttg1 | pituitary tumor-transforming gene 1(Pttg1) | -0.268946 |
| Pdgfra | platelet derived growth factor receptor, alpha polypeptide(Pdgfra) | 0.288914 |
| Pole | polymerase (DNA directed), epsilon(Pole) | 0.2455 |
| Rras | related RAS viral (r-ras) oncogene(Rras) | -0.397615 |
| Trp53inp1 | transformation related protein 53 inducible nuclear protein 1(Trp53inp1) | 0.269027 |
| Tgfbr2 | transforming growth factor, beta receptor II(Tgfbr2) | -0.402446 |
| **Neuroactive ligand-receptor interaction** | | |
| **Symbol** | **Gene name** | **Log2 (fold change)** |
| Htr6 | 5-hydroxytryptamine (serotonin) receptor 6(Htr6) | 0.400014 |
| Adora2a | adenosine A2a receptor(Adora2a) | -0.284827 |
| Adra1b | adrenergic receptor, alpha 1b(Adra1b) | -0.34772 |
| Adra2a | adrenergic receptor, alpha 2a(Adra2a) | 0.205162 |
| Cckbr | cholecystokinin B receptor(Cckbr) | -0.562344 |
| F2r | coagulation factor II (thrombin) receptor(F2r) | -0.221243 |
| Crhr1 | corticotropin releasing hormone receptor 1(Crhr1) | -0.311301 |
| Gabra2 | gamma-aminobutyric acid (GABA) A receptor, subunit alpha 2(Gabra2) | -1.9779 |
| Gabrd | gamma-aminobutyric acid (GABA) A receptor, subunit delta(Gabrd) | -0.501806 |
| Grik1 | glutamate receptor, ionotropic, kainate 1(Grik1) | 0.285063 |
| Grm2 | glutamate receptor, metabotropic 2(Grm2) | -0.408961 |
| Grm8 | glutamate receptor, metabotropic 8(Grm8) | -0.504899 |
| Lpar4 | lysophosphatidic acid receptor 4(Lpar4) | 0.46842 |
| Npy2r | neuropeptide Y receptor Y2(Npy2r) | -0.230785 |
| Oxtr | oxytocin receptor(Oxtr) | 0.267845 |
| Pth1r | parathyroid hormone 1 receptor(Pth1r) | -0.735892 |
| P2rx3 | purinergic receptor P2X, ligand-gated ion channel, 3(P2rx3) | -0.297367 |
| P2rx6 | purinergic receptor P2X, ligand-gated ion channel, 6(P2rx6) | -0.302309 |
| P2ry1 | purinergic receptor P2Y, G-protein coupled 1(P2ry1) | -0.317979 |
| P2ry2 | purinergic receptor P2Y, G-protein coupled 2(P2ry2) | -0.415118 |
| Sstr3 | somatostatin receptor 3(Sstr3) | -0.225749 |
| Tacr3 | tachykinin receptor 3(Tacr3) | -0.345044 |
| Thrb | thyroid hormone receptor beta(Thrb) | -0.223796 |
| Trhr | thyrotropin releasing hormone receptor(Trhr) | -0.253903 |
| **Pathways in cancer** | | |
| **Symbol** | **Gene name** | **Log2 (fold change)** |
| Rasgrp1 | RAS guanyl releasing protein 1(Rasgrp1) | -0.302872 |
| Arhgef1 | Rho guanine nucleotide exchange factor (GEF) 1(Arhgef1) | 0.236485 |
| Traf3 | TNF receptor-associated factor 3(Traf3) | 0.205892 |
| Apc2 | adenomatosis polyposis coli 2(Apc2) | 0.236055 |
| Apc | adenomatosis polyposis coli(Apc) | 0.311095 |
| Adcy5 | adenylate cyclase 5(Adcy5) | 0.18128 |
| Adcy8 | adenylate cyclase 8(Adcy8) | -0.219063 |
| Birc5 | baculoviral IAP repeat-containing 5(Birc5) | 0.315857 |
| Bmp4 | bone morphogenetic protein 4(Bmp4) | -0.338676 |
| F2r | coagulation factor II (thrombin) receptor(F2r) | -0.221243 |
| Csf2ra | colony stimulating factor 2 receptor, alpha, low-affinity (granulocyte-macrophage)(Csf2ra) | 0.294034 |
| Ccnd1 | cyclin D1(Ccnd1) | 0.18533 |
| Ccne1 | cyclin E1(Ccne1) | -0.260069 |
| Dvl1 | dishevelled segment polarity protein 1(Dvl1) | 0.183879 |
| Egfr | epidermal growth factor receptor(Egfr) | 0.283762 |
| Fgf9 | fibroblast growth factor 9(Fgf9) | -0.394831 |
| Fn1 | fibronectin 1(Fn1) | 0.312679 |
| Gng11 | guanine nucleotide binding protein (G protein), gamma 11(Gng11) | -0.799539 |
| Hgf | hepatocyte growth factor(Hgf) | 0.423115 |
| Itgav | integrin alpha V(Itgav) | 0.229906 |
| Lama5 | laminin, alpha 5(Lama5) | 0.20888 |
| Lamc2 | laminin, gamma 2(Lamc2) | -0.526442 |
| Lpar4 | lysophosphatidic acid receptor 4(Lpar4) | 0.46842 |
| Met | met proto-oncogene(Met) | 0.198161 |
| Map2k1 | mitogen-activated protein kinase kinase 1(Map2k1) | -0.23032 |
| Msh2 | mutS homolog 2(Msh2) | -0.213602 |
| Pdgfra | platelet derived growth factor receptor, alpha polypeptide(Pdgfra) | 0.288914 |
| Ptgs2 | prostaglandin-endoperoxide synthase 2(Ptgs2) | -0.380668 |
| Ret | ret proto-oncogene(Ret) | -0.284101 |
| Runx1t1 | runt-related transcription factor 1; translocated to, 1 (cyclin D-related)(Runx1t1) | 0.218432 |
| Tgfbr2 | transforming growth factor, beta receptor II(Tgfbr2) | -0.402446 |
