## Supplementary Table 4 for "Computational analysis of transcriptome signature repurposes low dose trifluoperazine for the treatment of fragile X syndrome in mouse model"

| Rank | Compound name and cell line | Similarity mean | N | Enrichment | P |
| --- | --- | --- | --- | --- | --- |
| 1 | trichostatin A - PC3 | -0.542 | 55 | -0.661 | 0 |
| 2 | tanespimycin - PC3 | -0.486 | 12 | -0.626 | 0 |
| 3 | 0175029-0000 - PC3 | -0.725 | 4 | -0.931 | 0.00002 |
| 4 | methotrexate - MCF7 | -0.828 | 3 | -0.972 | 0.00008 |
| 5 | etoposide - MCF7 | -0.869 | 2 | -0.993 | 0.00022 |
| 6 | phenoxybenzamine - MCF7 | -0.656 | 3 | -0.947 | 0.00026 |
| 7 | trifluoperazine - PC3 | -0.741 | 3 | -0.917 | 0.00096 |
| 8 | luteolin - MCF7 | -0.817 | 2 | -0.975 | 0.00137 |
| 9 | thioridazine - PC3 | -0.635 | 5 | -0.738 | 0.00248 |
| 10 | Prestwick-642 - MCF7 | 0.608 | 2 | 0.958 | 0.00308 |
| 11 | doxorubicin - MCF7 | -0.744 | 2 | -0.959 | 0.00364 |
| 12 | mefloquine - PC3 | -0.768 | 2 | -0.958 | 0.0038 |
| 13 | piribedil - MCF7 | 0.68 | 2 | 0.954 | 0.00392 |
| 14 | chrysin - MCF7 | -0.731 | 2 | -0.957 | 0.00394 |
| 15 | 5230742 - MCF7 | 0.788 | 2 | 0.953 | 0.00404 |
| 16 | pilocarpine - MCF7 | 0.617 | 2 | 0.951 | 0.00447 |
| 17 | sulfafurazole - MCF7 | 0.312 | 3 | 0.865 | 0.00461 |
| 18 | 0316684-0000 - MCF7 | 0.625 | 2 | 0.948 | 0.00495 |
| 19 | 15-delta prostaglandin J2 - PC3 | -0.641 | 3 | -0.864 | 0.00511 |
| 20 | GW-8510 - PC3 | -0.748 | 2 | -0.947 | 0.00614 |
| 21 | betonicine - MCF7 | 0.41 | 3 | 0.849 | 0.00645 |
| 22 | thiamine - MCF7 | 0.586 | 2 | 0.94 | 0.0068 |
| 23 | lobelanidine - MCF7 | 0.597 | 2 | 0.938 | 0.00742 |
| 24 | cloperastine - MCF7 | -0.614 | 3 | -0.841 | 0.00809 |
| 25 | PF-00562151-00 - PC3 | -0.43 | 4 | -0.737 | 0.00949 |
| 26 | cefazolin - PC3 | -0.621 | 2 | -0.93 | 0.01014 |
| 27 | aztreonam - MCF7 | -0.665 | 2 | -0.927 | 0.01097 |
| 28 | mycophenolic acid - MCF7 | -0.656 | 2 | -0.927 | 0.01109 |
| 29 | alsterpaullone - PC3 | -0.75 | 2 | -0.925 | 0.01159 |
| 30 | thiostrepton - MCF7 | -0.612 | 2 | -0.924 | 0.01181 |
| 31 | PHA-00767505E - MCF7 | 0.543 | 2 | 0.92 | 0.01264 |
| 32 | astemizole - PC3 | -0.795 | 2 | -0.921 | 0.01266 |
| 33 | 0173570-0000 - PC3 | -0.592 | 4 | -0.715 | 0.01337 |
| 34 | nordihydroguaiaretic acid - MCF7 | -0.449 | 8 | -0.526 | 0.0137 |
| 35 | troglitazone - PC3 | -0.434 | 4 | -0.713 | 0.01383 |
| 36 | NU-1025 - MCF7 | 0.537 | 2 | 0.916 | 0.01404 |
| 37 | fenbufen - PC3 | -0.638 | 2 | -0.912 | 0.01555 |
| 38 | geldanamycin - MCF7 | -0.246 | 10 | -0.467 | 0.01617 |
| 39 | perphenazine - PC3 | -0.601 | 2 | -0.909 | 0.0167 |
| 40 | amitriptyline - MCF7 | -0.623 | 3 | -0.796 | 0.01723 |
| 41 | tridihexethyl - MCF7 | 0.542 | 2 | 0.908 | 0.01748 |
| 42 | moroxydine - MCF7 | 0.549 | 2 | 0.907 | 0.01785 |
| 43 | amylocaine - MCF7 | 0.529 | 2 | 0.907 | 0.01801 |
| 44 | chlorcyclizine - MCF7 | -0.506 | 3 | -0.792 | 0.01823 |
| 45 | triamcinolone - MCF7 | -0.607 | 2 | -0.904 | 0.01845 |
| 46 | glafenine - MCF7 | 0.494 | 2 | 0.905 | 0.01847 |
| 47 | levomepromazine - PC3 | -0.653 | 2 | -0.903 | 0.01869 |
| 48 | resveratrol - MCF7 | -0.509 | 6 | -0.582 | 0.01873 |
| 49 | 0175029-0000 - MCF7 | -0.687 | 2 | -0.902 | 0.01899 |
| 50 | etofylline - PC3 | 0.669 | 2 | 0.903 | 0.01905 |
| 51 | monorden - PC3 | -0.466 | 5 | -0.625 | 0.01933 |
| 52 | ivermectin - MCF7 | -0.598 | 2 | -0.901 | 0.01968 |
| 53 | propofol - MCF7 | -0.613 | 2 | -0.901 | 0.01976 |
| 54 | thiethylperazine - MCF7 | -0.596 | 2 | -0.899 | 0.0203 |
| 55 | benfluorex - MCF7 | 0.598 | 2 | 0.9 | 0.0204 |
| 56 | alvespimycin - PC3 | -0.658 | 2 | -0.898 | 0.02066 |
| 57 | dinoprost - MCF7 | 0.544 | 2 | 0.898 | 0.02167 |
| 58 | chloramphenicol - PC3 | 0.509 | 2 | 0.896 | 0.02231 |
| 59 | oxprenolol - MCF7 | -0.639 | 2 | -0.894 | 0.02251 |
| 60 | pentoxifylline - PC3 | 0.483 | 2 | 0.893 | 0.0236 |
| 61 | methylbenzethonium chloride - PC3 | -0.58 | 2 | -0.89 | 0.02394 |
| 62 | withaferin A - PC3 | -0.608 | 2 | -0.888 | 0.02511 |
| 63 | tetrandrine - MCF7 | -0.629 | 2 | -0.886 | 0.02602 |
| 64 | CP-690334-01 - PC3 | -0.281 | 4 | -0.671 | 0.0261 |
| 65 | fulvestrant - PC3 | -0.197 | 12 | -0.407 | 0.0262 |
| 66 | tiletamine - MCF7 | 0.474 | 2 | 0.885 | 0.0272 |
| 67 | levomepromazine - MCF7 | -0.626 | 2 | -0.882 | 0.02791 |
| 68 | paclitaxel - MCF7 | 0.358 | 3 | 0.757 | 0.0285 |
| 69 | canrenoic acid - PC3 | -0.549 | 2 | -0.88 | 0.02907 |
| 70 | acacetin - PC3 | -0.589 | 2 | -0.877 | 0.03044 |
| 71 | triamterene - PC3 | -0.56 | 2 | -0.876 | 0.03094 |
| 72 | sulfadimidine - PC3 | 0.422 | 2 | 0.873 | 0.03245 |
| 73 | metformin - PC3 | 0.567 | 2 | 0.871 | 0.03354 |
| 74 | sulfinpyrazone - MCF7 | -0.635 | 2 | -0.869 | 0.03406 |
| 75 | chlorcyclizine - PC3 | -0.558 | 2 | -0.869 | 0.03438 |
| 76 | thioridazine - HL60 | -0.464 | 4 | -0.652 | 0.03467 |
| 77 | thioguanosine - MCF7 | -0.658 | 2 | -0.868 | 0.03469 |
| 78 | 3-nitropropionic acid - PC3 | 0.469 | 2 | 0.868 | 0.03521 |
| 79 | dobutamine - MCF7 | 0.413 | 2 | 0.868 | 0.03543 |
| 80 | zidovudine - MCF7 | 0.221 | 2 | 0.864 | 0.0376 |
| 81 | propranolol - MCF7 | 0.343 | 2 | 0.864 | 0.03772 |
| 82 | CP-690334-01 - MCF7 | -0.374 | 4 | -0.645 | 0.03796 |
| 83 | parbendazole - PC3 | 0.223 | 2 | 0.86 | 0.03964 |
| 84 | dilazep - PC3 | -0.54 | 2 | -0.857 | 0.04088 |
| 85 | fenoprofen - MCF7 | 0.439 | 3 | 0.723 | 0.04172 |
| 86 | PHA-00745360 - MCF7 | -0.238 | 4 | -0.637 | 0.04182 |
| 87 | probenecid - MCF7 | 0.182 | 2 | 0.856 | 0.04191 |
| 88 | alpha-ergocryptine - MCF7 | -0.458 | 3 | -0.726 | 0.04232 |
| 89 | procarbazine - MCF7 | 0.222 | 2 | 0.855 | 0.04243 |
| 90 | sulfapyridine - MCF7 | 0.273 | 2 | 0.855 | 0.04286 |
| 91 | raubasine - MCF7 | 0.231 | 2 | 0.854 | 0.04308 |
| 92 | fenbendazole - PC3 | 0.26 | 2 | 0.854 | 0.04338 |
| 93 | pioglitazone - PC3 | -0.233 | 5 | -0.57 | 0.04408 |
| 94 | copper sulfate - MCF7 | -0.56 | 2 | -0.851 | 0.04414 |
| 95 | kanamycin - MCF7 | -0.547 | 2 | -0.85 | 0.04481 |
| 96 | amikacin - MCF7 | -0.547 | 2 | -0.85 | 0.04485 |
| 97 | diltiazem - MCF7 | 0.209 | 2 | 0.846 | 0.0477 |
| 98 | cefsulodin - MCF7 | 0.193 | 2 | 0.844 | 0.04871 |
| 99 | quercetin - MCF7 | -0.467 | 4 | -0.625 | 0.04894 |
| 100 | nitrendipine - MCF7 | -0.339 | 3 | -0.711 | 0.04927 |
| 101 | pimozide - MCF7 | -0.584 | 2 | -0.843 | 0.04966 |
| 102 | cyclobenzaprine - MCF7 | -0.662 | 2 | -0.842 | 0.04994 |
