## Supplementary Table 5 for "Computational analysis of transcriptome signature repurposes low dose trifluoperazine for the treatment of fragile X syndrome in mouse model"

| Rank | Compound name and cell line | Similarity mean | N | Enrichment | P |
| --- | --- | --- | --- | --- | --- |
| 1 | fluphenazine - PC3 | 0.558 | 3 | 0.997 | 0 |
| 2 | thioridazine - PC3 | 0.569 | 5 | 0.994 | 0 |
| 3 | vorinostat - MCF7 | 0.333 | 7 | 0.92 | 0 |
| 4 | sirolimus - PC3 | 0.344 | 8 | 0.871 | 0 |
| 5 | prochlorperazine - MCF7 | 0.369 | 9 | 0.867 | 0 |
| 6 | trichostatin A - PC3 | 0.347 | 55 | 0.862 | 0 |
| 7 | trifluoperazine - MCF7 | 0.371 | 9 | 0.787 | 0 |
| 8 | geldanamycin - MCF7 | 0.294 | 10 | 0.753 | 0 |
| 9 | LY-294002 - PC3 | 0.315 | 12 | 0.748 | 0 |
| 10 | trichostatin A - MCF7 | 0.29 | 92 | 0.729 | 0 |
| 11 | thioridazine - MCF7 | 0.373 | 11 | 0.724 | 0 |
| 12 | tanespimycin - MCF7 | 0.273 | 36 | 0.663 | 0 |
| 13 | LY-294002 - MCF7 | 0.166 | 34 | 0.424 | 0 |
| 14 | methylbenzethonium chloride - PC3 | 0.596 | 2 | 0.998 | 0.00002 |
| 15 | astemizole - PC3 | 0.554 | 2 | 0.996 | 0.00002 |
| 16 | mefloquine - PC3 | 0.555 | 2 | 0.996 | 0.00002 |
| 17 | fluphenazine - MCF7 | 0.315 | 10 | 0.724 | 0.00002 |
| 18 | PHA-00846566E - PC3 | -0.864 | 2 | -0.998 | 0.00004 |
| 19 | prochlorperazine - PC3 | 0.525 | 3 | 0.968 | 0.00004 |
| 20 | pyrvinium - MCF7 | 0.374 | 4 | 0.921 | 0.00004 |
| 21 | fulvestrant - MCF7 | 0.207 | 21 | 0.511 | 0.00004 |
| 22 | perphenazine - PC3 | 0.5 | 2 | 0.992 | 0.00006 |
| 23 | tanespimycin - PC3 | 0.272 | 12 | 0.637 | 0.00006 |
| 24 | phenoxybenzamine - MCF7 | 0.35 | 3 | 0.951 | 0.0001 |
| 25 | thioridazine - HL60 | 0.327 | 4 | 0.9 | 0.0001 |
| 26 | AR-A014418 - PC3 | -0.816 | 2 | -0.994 | 0.00012 |
| 27 | alexidine - PC3 | 0.474 | 2 | 0.99 | 0.00012 |
| 28 | promazine - PC3 | 0.486 | 2 | 0.989 | 0.00016 |
| 29 | alvespimycin - MCF7 | 0.271 | 7 | 0.756 | 0.00016 |
| 30 | trichostatin A - HL60 | 0.155 | 34 | 0.368 | 0.0002 |
| 31 | cloperastine - MCF7 | 0.364 | 3 | 0.946 | 0.00022 |
| 32 | loperamide - MCF7 | 0.369 | 3 | 0.943 | 0.00022 |
| 33 | niclosamide - PC3 | 0.452 | 2 | 0.988 | 0.00026 |
| 34 | lycorine - MCF7 | -0.638 | 3 | -0.94 | 0.00032 |
| 35 | wortmannin - MCF7 | 0.244 | 10 | 0.608 | 0.00038 |
| 36 | terfenadine - MCF7 | 0.441 | 2 | 0.985 | 0.0004 |
| 37 | pioglitazone - PC3 | -0.422 | 5 | -0.823 | 0.00042 |
| 38 | 15-delta prostaglandin J2 - MCF7 | 0.301 | 8 | 0.672 | 0.00042 |
| 39 | CAY-10397 - PC3 | -0.734 | 2 | -0.985 | 0.00048 |
| 40 | troglitazone - MCF7 | 0.256 | 7 | 0.699 | 0.0005 |
| 41 | fluspirilene - MCF7 | 0.423 | 2 | 0.983 | 0.00056 |
| 42 | 5707885 - PC3 | 0.417 | 2 | 0.982 | 0.00062 |
| 43 | cloperastine - PC3 | 0.438 | 2 | 0.981 | 0.00062 |
| 44 | sirolimus - MCF7 | 0.18 | 25 | 0.389 | 0.00066 |
| 45 | trifluoperazine - HL60 | 0.292 | 4 | 0.853 | 0.00068 |
| 46 | astemizole - MCF7 | 0.428 | 2 | 0.98 | 0.0007 |
| 47 | rottlerin - MCF7 | 0.362 | 3 | 0.927 | 0.00072 |
| 48 | beta-escin - PC3 | 0.422 | 2 | 0.98 | 0.00074 |
| 49 | suloctidil - MCF7 | 0.427 | 2 | 0.979 | 0.00076 |
| 50 | resveratrol - MCF7 | 0.288 | 6 | 0.744 | 0.00077 |
| 51 | withaferin A - PC3 | 0.403 | 2 | 0.978 | 0.00078 |
| 52 | chlorcyclizine - PC3 | 0.423 | 2 | 0.978 | 0.00082 |
| 53 | Prestwick-675 - PC3 | -0.835 | 2 | -0.98 | 0.00085 |
| 54 | perhexiline - MCF7 | 0.402 | 2 | 0.977 | 0.00087 |
| 55 | clotrimazole - MCF7 | 0.345 | 3 | 0.924 | 0.00092 |
| 56 | benzethonium chloride - MCF7 | 0.4 | 2 | 0.974 | 0.00107 |
| 57 | azacyclonol - PC3 | 0.435 | 2 | 0.97 | 0.00159 |
| 58 | methylbenzethonium chloride - MCF7 | 0.355 | 3 | 0.909 | 0.00164 |
| 59 | semustine - PC3 | 0.381 | 2 | 0.969 | 0.00165 |
| 60 | prenylamine - MCF7 | 0.392 | 2 | 0.969 | 0.00165 |
| 61 | mefloquine - MCF7 | 0.381 | 2 | 0.967 | 0.00181 |
| 62 | clozapine - PC3 | 0.325 | 3 | 0.903 | 0.00182 |
| 63 | ivermectin - PC3 | 0.394 | 2 | 0.966 | 0.00191 |
| 64 | niclosamide - MCF7 | 0.384 | 2 | 0.966 | 0.00205 |
| 65 | Chicago Sky Blue 6B - MCF7 | -0.67 | 2 | -0.968 | 0.00225 |
| 66 | diprophylline - PC3 | -0.683 | 2 | -0.966 | 0.00262 |
| 67 | amitriptyline - PC3 | 0.422 | 2 | 0.96 | 0.00266 |
| 68 | thiostrepton - MCF7 | 0.376 | 2 | 0.958 | 0.00304 |
| 69 | pimozide - MCF7 | 0.368 | 2 | 0.958 | 0.00308 |
| 70 | 15-delta prostaglandin J2 - HL60 | 0.308 | 3 | 0.884 | 0.00318 |
| 71 | pyrithyldione - MCF7 | -0.629 | 2 | -0.961 | 0.00334 |
| 72 | lomustine - PC3 | 0.379 | 2 | 0.955 | 0.00358 |
| 73 | monorden - MCF7 | 0.197 | 12 | 0.489 | 0.00381 |
| 74 | wortmannin - PC3 | 0.389 | 2 | 0.95 | 0.00455 |
| 75 | amiodarone - MCF7 | 0.285 | 3 | 0.865 | 0.00461 |
| 76 | scriptaid - PC3 | 0.345 | 2 | 0.949 | 0.00481 |
| 77 | mebendazole - PC3 | 0.357 | 2 | 0.947 | 0.00499 |
| 78 | nortriptyline - MCF7 | 0.373 | 2 | 0.947 | 0.00505 |
| 79 | nordihydroguaiaretic acid - MCF7 | 0.235 | 8 | 0.57 | 0.00536 |
| 80 | iohexol - MCF7 | -0.604 | 2 | -0.95 | 0.00547 |
| 81 | miconazole - PC3 | 0.358 | 2 | 0.945 | 0.00577 |
| 82 | arecoline - MCF7 | -0.579 | 2 | -0.948 | 0.00588 |
| 83 | disulfiram - PC3 | 0.35 | 2 | 0.944 | 0.00596 |
| 84 | sulconazole - MCF7 | 0.346 | 2 | 0.943 | 0.00606 |
| 85 | MG-262 - PC3 | 0.352 | 2 | 0.943 | 0.0062 |
| 86 | tonzonium bromide - MCF7 | 0.339 | 2 | 0.942 | 0.0063 |
| 87 | ketorolac - MCF7 | -0.564 | 2 | -0.946 | 0.00632 |
| 88 | isoconazole - PC3 | 0.343 | 2 | 0.942 | 0.00632 |
| 89 | metolazone - MCF7 | -0.573 | 2 | -0.945 | 0.00666 |
| 90 | bepridil - MCF7 | 0.344 | 2 | 0.94 | 0.0068 |
| 91 | thioguanosine - MCF7 | 0.355 | 2 | 0.94 | 0.00704 |
| 92 | MS-275 - PC3 | 0.358 | 2 | 0.938 | 0.00755 |
| 93 | loperamide - PC3 | 0.377 | 2 | 0.937 | 0.00765 |
| 94 | vorinostat - PC3 | 0.37 | 2 | 0.936 | 0.00781 |
| 95 | fendiline - MCF7 | 0.354 | 2 | 0.934 | 0.00829 |
| 96 | withaferin A - MCF7 | 0.341 | 2 | 0.934 | 0.00833 |
| 97 | clemizole - PC3 | 0.332 | 2 | 0.933 | 0.00849 |
| 98 | clopamide - MCF7 | -0.527 | 2 | -0.935 | 0.00885 |
| 99 | rescinnamine - MCF7 | 0.348 | 2 | 0.931 | 0.00915 |
| 100 | perphenazine - MCF7 | 0.363 | 2 | 0.93 | 0.00942 |
| 101 | 5224221 - MCF7 | 0.349 | 2 | 0.929 | 0.00962 |
| 102 | pivmecillinam - MCF7 | -0.553 | 2 | -0.93 | 0.01028 |
| 103 | metergoline - MCF7 | 0.347 | 2 | 0.926 | 0.01046 |
| 104 | protriptyline - MCF7 | 0.373 | 2 | 0.926 | 0.01058 |
| 105 | ionomycin - MCF7 | 0.339 | 3 | 0.825 | 0.01074 |
| 106 | quinostatin - MCF7 | 0.33 | 2 | 0.924 | 0.01127 |
| 107 | isocarboxazid - MCF7 | -0.506 | 2 | -0.923 | 0.01205 |
| 108 | isometheptene - MCF7 | -0.509 | 2 | -0.923 | 0.01213 |
| 109 | 0297417-0002B - MCF7 | 0.318 | 2 | 0.92 | 0.01247 |
| 110 | clozapine - MCF7 | 0.136 | 10 | 0.478 | 0.01264 |
| 111 | nadide - MCF7 | -0.507 | 2 | -0.921 | 0.01268 |
| 112 | cyproheptadine - PC3 | 0.363 | 2 | 0.919 | 0.01278 |
| 113 | trimethoprim - PC3 | -0.639 | 2 | -0.92 | 0.01292 |
| 114 | fursultiamine - MCF7 | -0.559 | 2 | -0.92 | 0.01302 |
| 115 | homochlorcyclizine - MCF7 | 0.371 | 2 | 0.919 | 0.01308 |
| 116 | fluphenazine - HL60 | 0.231 | 4 | 0.716 | 0.01319 |
| 117 | mianserin - PC3 | 0.339 | 2 | 0.918 | 0.01336 |
| 118 | tetrandrine - MCF7 | 0.334 | 2 | 0.917 | 0.0136 |
| 119 | disopyramide - MCF7 | -0.507 | 2 | -0.916 | 0.01424 |
| 120 | CP-320650-01 - PC3 | -0.423 | 4 | -0.71 | 0.01428 |
| 121 | harmol - MCF7 | -0.494 | 2 | -0.916 | 0.01431 |
| 122 | econazole - MCF7 | 0.325 | 2 | 0.915 | 0.01435 |
| 123 | proadifen - MCF7 | 0.337 | 2 | 0.914 | 0.01443 |
| 124 | miconazole - MCF7 | 0.341 | 2 | 0.913 | 0.01531 |
| 125 | lanatoside C - MCF7 | 0.257 | 3 | 0.802 | 0.01576 |
| 126 | colecalciferol - MCF7 | -0.488 | 2 | -0.911 | 0.01591 |
| 127 | PNU-0251126 - MCF7 | -0.434 | 2 | -0.91 | 0.01632 |
| 128 | carbimazole - MCF7 | -0.484 | 2 | -0.908 | 0.01686 |
| 129 | syrosingopine - MCF7 | 0.334 | 2 | 0.908 | 0.01752 |
| 130 | sulfaphenazole - MCF7 | -0.541 | 2 | -0.905 | 0.01819 |
| 131 | puromycin - MCF7 | 0.317 | 2 | 0.906 | 0.01825 |
| 132 | fenbendazole - PC3 | 0.347 | 2 | 0.906 | 0.01827 |
| 133 | deptropine - MCF7 | 0.312 | 2 | 0.904 | 0.01873 |
| 134 | geldanamycin - PC3 | 0.322 | 2 | 0.903 | 0.01932 |
| 135 | flupentixol - MCF7 | 0.305 | 2 | 0.903 | 0.01934 |
| 136 | iloprost - MCF7 | -0.443 | 2 | -0.902 | 0.01936 |
| 137 | bucladesine - MCF7 | -0.218 | 4 | -0.69 | 0.02003 |
| 138 | parthenolide - MCF7 | 0.332 | 2 | 0.9 | 0.02058 |
| 139 | flumetasone - PC3 | -0.324 | 2 | -0.897 | 0.02117 |
| 140 | antimycin A - MCF7 | 0.314 | 2 | 0.898 | 0.02169 |
| 141 | alverine - PC3 | 0.31 | 2 | 0.897 | 0.02215 |
| 142 | carbamazepine - MCF7 | 0.229 | 5 | 0.623 | 0.02221 |
| 143 | demeclocycline - PC3 | -0.3 | 2 | -0.893 | 0.02286 |
| 144 | dicoumarol - PC3 | -0.347 | 2 | -0.893 | 0.02316 |
| 145 | flunixin - PC3 | -0.306 | 2 | -0.892 | 0.02338 |
| 146 | chlorpromazine - PC3 | 0.354 | 4 | 0.677 | 0.02379 |
| 147 | oxolinic acid - PC3 | -0.301 | 2 | -0.891 | 0.02386 |
| 148 | haloperidol - PC3 | 0.233 | 6 | 0.567 | 0.02427 |
| 149 | geldanamycin - HL60 | 0.253 | 3 | 0.769 | 0.02458 |
| 150 | calmidazolium - MCF7 | 0.346 | 2 | 0.891 | 0.02463 |
| 151 | metitepine - MCF7 | 0.295 | 2 | 0.89 | 0.02515 |
| 152 | oxetacaine - PC3 | 0.293 | 2 | 0.889 | 0.02529 |
| 153 | nordihydroguaiaretic acid - PC3 | 0.293 | 2 | 0.887 | 0.0263 |
| 154 | phentolamine - PC3 | 0.297 | 2 | 0.887 | 0.0264 |
| 155 | sertaconazole - MCF7 | 0.287 | 2 | 0.887 | 0.02648 |
| 156 | pizotifen - MCF7 | 0.29 | 2 | 0.886 | 0.02672 |
| 157 | clomifene - MCF7 | 0.355 | 2 | 0.885 | 0.02714 |
| 158 | dequalinium chloride - MCF7 | 0.297 | 2 | 0.885 | 0.02732 |
| 159 | norcyclobenzaprine - MCF7 | 0.343 | 2 | 0.885 | 0.02734 |
| 160 | thapsigargin - MCF7 | 0.306 | 2 | 0.882 | 0.02851 |
| 161 | hexetidine - MCF7 | 0.332 | 2 | 0.881 | 0.02875 |
| 162 | ivermectin - MCF7 | 0.308 | 2 | 0.88 | 0.02893 |
| 163 | amoxapine - PC3 | 0.343 | 2 | 0.88 | 0.02899 |
| 164 | felodipine - MCF7 | 0.247 | 5 | 0.605 | 0.02938 |
| 165 | etofylline - PC3 | -0.341 | 2 | -0.879 | 0.02946 |
| 166 | methotrexate - MCF7 | 0.248 | 3 | 0.753 | 0.02946 |
| 167 | butoconazole - MCF7 | 0.285 | 2 | 0.879 | 0.0295 |
| 168 | piperacillin - PC3 | -0.398 | 2 | -0.878 | 0.0298 |
| 169 | oxaprozin - PC3 | -0.278 | 2 | -0.877 | 0.03024 |
| 170 | oxetacaine - MCF7 | 0.291 | 2 | 0.877 | 0.03064 |
| 171 | valproic acid - PC3 | 0.152 | 10 | 0.436 | 0.03115 |
| 172 | prochlorperazine - HL60 | 0.246 | 4 | 0.655 | 0.03268 |
| 173 | gossypol - MCF7 | 0.272 | 3 | 0.741 | 0.03379 |
| 174 | helveticoside - PC3 | 0.296 | 2 | 0.869 | 0.03477 |
| 175 | oleandomycin - PC3 | -0.249 | 2 | -0.867 | 0.03501 |
| 176 | aminophenazone - PC3 | -0.346 | 2 | -0.867 | 0.03507 |
| 177 | nicergoline - MCF7 | 0.286 | 2 | 0.868 | 0.03539 |
| 178 | dexamethasone - PC3 | -0.46 | 2 | -0.867 | 0.03541 |
| 179 | homatropine - PC3 | -0.446 | 2 | -0.867 | 0.03549 |
| 180 | thiamazole - PC3 | -0.275 | 2 | -0.867 | 0.03569 |
| 181 | clorgiline - MCF7 | 0.292 | 2 | 0.868 | 0.03591 |
| 182 | naltrexone - PC3 | -0.359 | 2 | -0.866 | 0.03626 |
| 183 | Prestwick-559 - MCF7 | 0.273 | 2 | 0.865 | 0.03694 |
| 184 | dilazep - PC3 | 0.274 | 2 | 0.865 | 0.03712 |
| 185 | cyanocobalamin - MCF7 | -0.328 | 2 | -0.863 | 0.03774 |
| 186 | Prestwick-1080 - MCF7 | -0.32 | 2 | -0.862 | 0.03825 |
| 187 | medrysone - MCF7 | 0.231 | 3 | 0.731 | 0.03834 |
| 188 | Prestwick-674 - PC3 | -0.257 | 2 | -0.861 | 0.03849 |
| 189 | quinisocaine - MCF7 | 0.278 | 2 | 0.861 | 0.03909 |
| 190 | meclozine - MCF7 | 0.251 | 3 | 0.729 | 0.0393 |
| 191 | 6-bromoindirubin-3'-oxime - MCF7 | -0.418 | 3 | -0.732 | 0.04002 |
| 192 | rifabutin - MCF7 | 0.273 | 2 | 0.857 | 0.04123 |
| 193 | verteporfin - MCF7 | 0.273 | 2 | 0.856 | 0.04219 |
| 194 | clomipramine - MCF7 | 0.322 | 2 | 0.855 | 0.04251 |
| 195 | genistein - HL60 | 0.22 | 3 | 0.721 | 0.04266 |
| 196 | helveticoside - MCF7 | 0.24 | 3 | 0.721 | 0.04293 |
| 197 | erastin - PC3 | 0.273 | 2 | 0.854 | 0.04318 |
| 198 | metixene - MCF7 | 0.297 | 2 | 0.852 | 0.04394 |
| 199 | nialamide - MCF7 | -0.28 | 2 | -0.851 | 0.04436 |
| 200 | sulfadiazine - PC3 | -0.367 | 2 | -0.851 | 0.04449 |
| 201 | allantoin - PC3 | -0.38 | 2 | -0.85 | 0.04519 |
| 202 | azacitidine - MCF7 | 0.27 | 2 | 0.85 | 0.04529 |
| 203 | pimethixene - MCF7 | 0.281 | 2 | 0.847 | 0.04736 |
| 204 | ouabain - MCF7 | 0.269 | 2 | 0.847 | 0.0475 |
| 205 | beta-escin - MCF7 | 0.246 | 3 | 0.71 | 0.04761 |
| 206 | sulfamethoxazole - MCF7 | -0.325 | 2 | -0.846 | 0.04766 |
| 207 | 16-phenyltetranorprostaglandin E2 - PC3 | 0.267 | 2 | 0.845 | 0.04823 |
| 208 | 2,6-dimethylpiperidine - PC3 | -0.294 | 2 | -0.845 | 0.04829 |
| 209 | todralazine - PC3 | -0.376 | 2 | -0.845 | 0.04841 |
| 210 | azathioprine - MCF7 | 0.212 | 4 | 0.626 | 0.04882 |
| 211 | nicergoline - PC3 | 0.283 | 2 | 0.844 | 0.04889 |
| 212 | tranylcypromine - PC3 | 0.276 | 2 | 0.844 | 0.04911 |
| 213 | tolfenamic acid - PC3 | 0.269 | 2 | 0.844 | 0.04915 |
